# CD56^dim^CD16^dim^ NK cells are the dominant effector cells against HIV-infected primary T-cells

**DOI:** 10.64898/2026.06.26.734820

**Authors:** William Howell, Courtney Branch, Jeffrey Ward, Zachary Davis, Emma Geatches, Edward Barker

## Abstract

Despite being rare among circulating natural killer (NK) cells and expressing 10-fold less CD16 than the predominant CD56^dim^CD16^bright^ population, CD56^dim^CD16^dim^ NK cells are expanded in HIV long-term elite controllers, yet their capacity to kill HIV-infected cells remained untested. Here, we show that these rare cells are the dominant effectors against HIV-infected T-cells, mediating approximately 4-fold higher direct cytotoxicity and 3-4-fold higher antibody-dependent cellular cytotoxicity (ADCC) than CD56^dim^CD16^bright^ cells, and serially engaging multiple targets. This advantage is intrinsic, unexplained by cytotoxic granule content or inhibitory receptors recognizing MHC class I. Direct killing depends on NKG2D recognition of Vpr-induced ligands, with NKG2D elevated on CD56^dim^CD16^dim^ cells; ADCC requires both NKG2D and ADAM17-mediated CD16 turnover for serial engagement. These findings explain the elite-controller reorganization, reveal that NK effector dominance is target-tuned rather than fixed (CD56^dim^CD16^negative^ cells dominate against K562 cells), and identify high-NKG2D CD56^dim^CD16^dim^ cells as the effector population HIV therapies should reproduce.

**Impact Statement:** A rare natural killer cell subset that makes up only a few percent of circulating NK cells, yet is enriched in the people who control HIV for years without medication, turns out to be the dominant killer of HIV-infected cells, far outperforming the common subset long assumed to do the job, and this work shows how these cells recognize and repeatedly attack infected targets, pointing to the specific effector that future cell-based therapies might use to help people with HIV stay healthy without lifelong drugs.

## Introduction

HIV affects approximately 40.8 million people worldwide [37.0-45.6 million] (UNAIDS 2024), causing a chronic infection that requires antiretroviral therapy (ART) to prevent progression to AIDS. Although ART effectively suppresses viral plasma levels (Antela et al., 2021; Puertas et al., 2025; Sarin & Goldstein, 1995), it cannot eradicate HIV-infected reservoirs that persist in anatomical sites, especially in gut-associated lymphoid tissue and lymph nodes, where ART’s access to infected cells appears limited (Belmonte et al., 2007; Chun et al., 2008; Estes et al., 2017; Fletcher et al., 2014; North et al., 2010; Thompson et al., 2017). Moreover, latent reservoirs cause viral rebound within weeks of stopping ART, making lifelong therapy necessary (Finzi et al., 1997; Magombedze et al., 2025; Puertas et al., 2025). Long-term ART has side effects, including accelerated aging, metabolic issues, loss of bone integrity, and cardiovascular problems (Ghandakly et al., 2025; Lagathu et al., 2019; Milburn et al., 2017; Olali et al., 2022; Vos & Venter, 2021). These limitations of ART highlight the urgent need for new, sustainable therapeutic strategies that can target and eliminate infected cells.

Natural killer (NK) cells are cytotoxic lymphocytes that eliminate virus-infected cells through direct cytotoxicity and antibody-dependent cellular cytotoxicity (ADCC) without prior sensitization (Lanier, Chang, et al., 1991; Lanier et al., 1983; Lanier, Yu, et al., 1991; Perussia et al., 1984). NK cell-based immunotherapies have shown success in treating leukemia, and haploidentical NK cell adoptive transfer is safe, does not cause graft-versus-host disease, cytokine release syndrome, or immune effector cell–associated neurotoxicity syndrome, and, under certain circumstances, can be sustained (Bednarski et al., 2022; Bjorklund et al., 2018; Cichocki et al., 2020; Cooley et al., 2019; Geller & Miller, 2011; Miller et al., 2005; Myers & Miller, 2021). However, NK cell-based therapeutic approaches to HIV have demonstrated limited effectiveness (Abeynaike et al., 2023; Kim et al., 2022), possibly due to uncertainty about which NK cell subset provides the best antiviral activity (Alter et al., 2005; Florez-Alvarez et al., 2020; Hu et al., 1995; Ivison et al., 2022; Lichtfuss et al., 2012; Mavilio et al., 2003; Pohlmeyer et al., 2019; Sanchez-Gaona et al., 2024).

Human NK cells comprise distinct subsets with varying functional capacities. The CD56^dim^CD16^bright^ population, representing ∼85% of circulating NK cells, has traditionally been considered the primary cytotoxic subset because of its high CD16 expression, which enables robust ADCC. (Cooper et al., 2001; Jacobs et al., 2001; Nagler et al., 1989). Cells with intermediate CD16 expression have been characterized as transitional or exhausted states in studies of tumors and other contexts, and their low frequency in peripheral blood meant that their potential contribution to HIV-specific responses had not been directly examined (Romee et al., 2013; Yang et al., 2019).

Together with the low frequency of CD56^dim^CD16^dim^ cells in the peripheral blood of uninfected individuals (<6%), this characterization led therapeutic development to focus on the more abundant CD56^dim^CD16^bright^ cell subset. However, recent observations have prompted a reexamination of this characterization (Amand et al., 2017; Rallon et al., 2024). Rallon et al. reported that long-term (>10 years since initial infection) HIV elite controllers (LTEC), a rare (<1%) subset of infected individuals who suppress HIV without ART, maintain expanded CD56^dim^CD16^dim^ populations, suggesting that these cells may contribute to viral control (Rallon et al., 2024). Moreover, the frequency of CD56^dim^CD16^bright^ NK cells is lower in LTEC than in uninfected controls and in people living with HIV who are on ART. Although this study reports a higher prevalence of CD56^dim^CD16^dim^ cells in LTEC, the functional capacity of these cells to eliminate HIV-infected targets has not been directly assessed (Rallon et al., 2024).

Here, we systematically compared the ability of NK cell subsets defined by CD16 expression to eliminate HIV-infected cells. We show that CD56^dim^CD16^dim^ cells, despite their rarity and lower CD16 density, are the dominant NK cell population that targets HIV-infected primary T-cells. This dominance reflects integrated NKG2D and ADAM17-dependent CD16 signaling, which supports both per-cell efficiency and serial engagement of multiple infected targets. These findings identify the natural effector cells configuration that cellular therapies aimed at an HIV functional cure can build toward.

## Results

### CD56^dim^CD16^dim^ NK cells show target-dependent enhanced functional responses

To understand why CD56^dim^CD16^dim^ NK cell frequencies correlate with HIV clinical outcomes (Amand et al., 2017; Rallon et al., 2024), even though prior work has not shown them to be universally superior effectors against tumor targets, we first validated these findings using the chronic myeloid leukemia cell line, K562, as the target cells (Amand et al., 2017; Herberman & Ortaldo, 1981; Ortaldo et al., 1977). We then specifically examined NK cell responses to autologous HIV-infected T-cells.

We began by characterizing the baseline distribution of NK cell subsets in healthy, uninfected donors. The CD56^negative^ NK cells were identified by NKp80 expression (Figure 1-figure supplement 1). In line with the established literature (Amand et al., 2017; Rallon et al., 2024), CD56^dim^ NK cells constituted the majority [mean=93.38% ± standard deviation (SD)= 0.96%] of circulating NK cells, with CD56^bright^ NK cells (4.46% ± 0.73%) and CD56^negative^ NK cells (2.04% ± 0.32%) accounting for a small fraction (Figure 1-figure supplement 1). Within the CD56^dim^ NK cell population, CD16^bright^ cells were predominant (91.84% ± 0.04%). In comparison, CD16^dim^ cells (4.56% ± 0.24%) and CD16^negative^ cells (3.82% ± 0.22%) were minor CD56^dim^ NK cell subsets (Figure 1-figure supplement 1).

To confirm the previous observations (Amand et al., 2017) regarding CD56^dim^CD16^dim^ functional capacity, we assessed NK cell degranulation responses against the chronic myeloid leukemia K562 cell line at an effector-to-target (E:T) ratio of 1:1. Consistent with previous findings (Amand et al., 2017), CD56^dim^CD16^negative^ NK cells showed significantly higher degranulation (surface CD107a^positive^) than CD56^dim^CD16^dim^ NK cells (Figure 1—figure supplement 2; donor 1, p=0.0018; donor 2, p<0.0001), with CD56^dim^CD16^bright^ cells showing the lowest responses (donor 1: 20.34 ± 2.39; donor 2: 12.24 ± 0.33; all between-subset comparisons within each donor p≤0.0018 by one-way ANOVA with Tukey’s multiple comparisons test; Supplemental Table 1). CD56^dim^CD16^negative^ NK cells showed the highest degranulation in donors 1 (85.63% ± 1.38%) and 2 (58.78% ± 1.13%), followed by CD56^dim^CD16^dim^ NK cells in donors 1 (71.17% ± 3.81%) and 2 (48.38% ± 0.56%). These results confirmed (Amand et al., 2017) that CD56^dim^CD16^dim^ NK cells do not exhibit a functional advantage over CD56^dim^CD16^negative^ NK cells in response to K562 target cells; on the contrary, CD56^dim^CD16^negative^ NK cells significantly outperform CD56^dim^CD16^dim^ NK cells in this context.

Given this knowledge gap between clinical relevance against HIV (Amand et al., 2017; Rallon et al., 2024) and functional capacity against tumor targets (Amand et al., 2017), we hypothesized that CD56^dim^CD16^dim^ cells might possess HIV-specific functional capacity. To test this, we examined NK cell responses against autologous HIV-infected T-cells using the same degranulation assay at an E:T ratio of 1:1. Notably, despite a low frequency (4.26% ± 0.24%) of CD56^dim^CD16^dim^ cells amongst the total population of NK cells (Figure 1A), a dramatically higher frequency of CD56^dim^CD16^dim^ cells that degranulated against HIV-infected targets (25.47% ± 3.75%) compared to all other NK cell subsets, which showed markedly lower responses (0.63% to 13.19%) (Figure 1B; one-way ANOVA F(6, 14) = 85.96, p<0.0001; Dunnett’s multiple comparisons test against CD56^dim^CD16^dim^, all p<0.0001; Supplemental Table 2).

**Figure 1:**
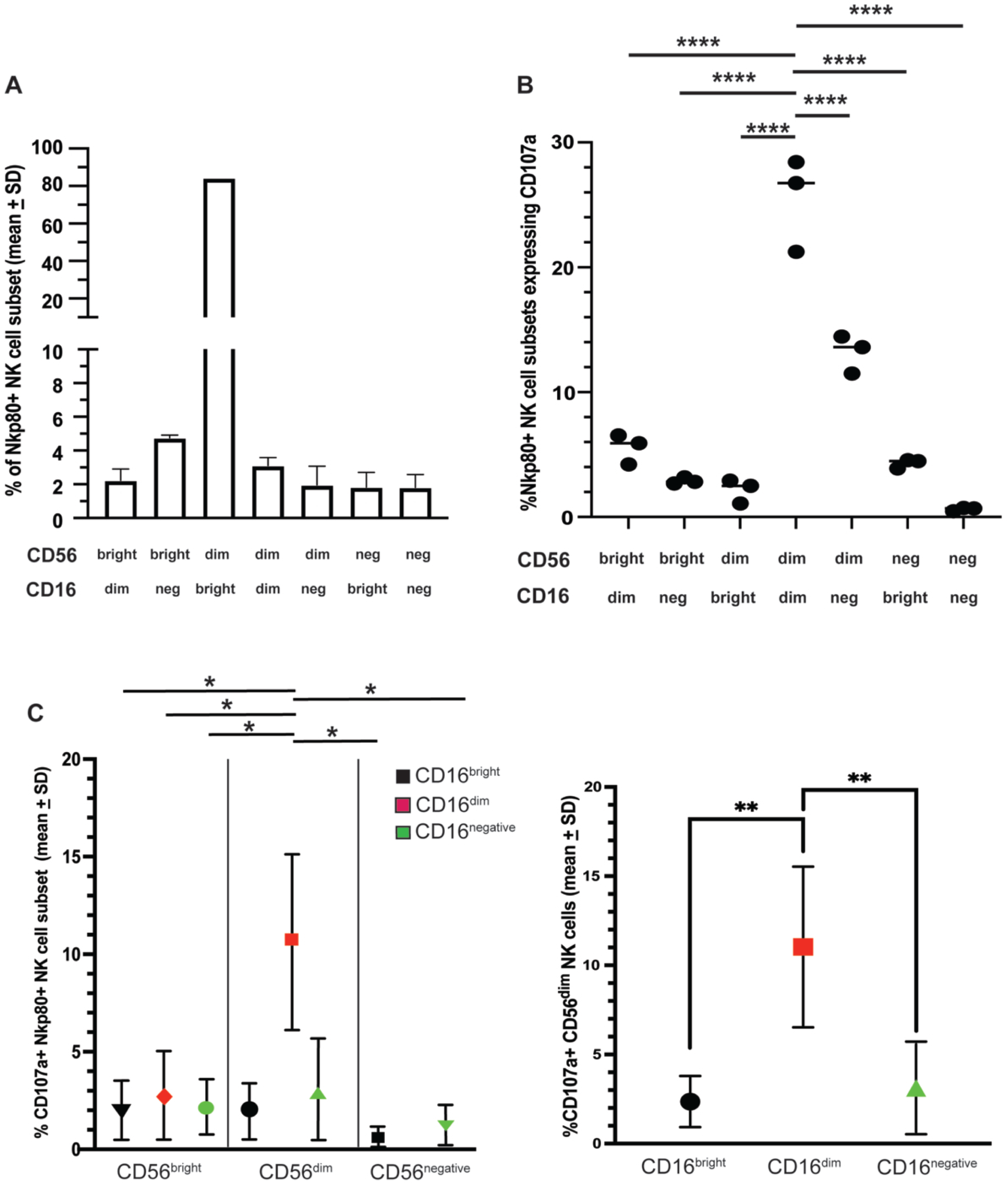
Degranulation of NK cell subsets based on differential expression of CD56 and CD16 in response to purified autologous HIV productively infected T-cells. (A) Frequency of NK cell subsets [mean + standard deviation (SD)] within total peripheral blood NK cells of an HIV-uninfected donor [the donor used for (B)], based on differential CD56 and CD16 expression. NKp80 was used to identify CD56^negative^ NK cell subsets. (B) Frequency of NK cell subsets expressing CD107a following a 4-hour exposure at a 1:1 effector cell to target cell (E:T) ratio to purified autologous productively HIV-1_SHM-1_ infected T-cells. Each circle represents a technical replicate; lines indicate mean values. All values are background-subtracted (target-exposed minus unexposed control). (C) Frequency of CD107a expression (mean + SD) on NK cell subsets across six different donors. The CD56^negative^ CD16^dim^ subset was excluded because its frequency in peripheral blood (<0.2% of NK cells; Figure 1-figure supplement 1B) yields event counts insufficient for reliable measurement. (D) Frequency of CD56^dim^ NK cell subsets [from (C)] that were also CD16^bright^, CD16^dim^, or CD16^negative^ across the same six donors. Panels A and B show data from one representative donor with three technical replicates; panels C and D show data from six donors analyzed in separate experiments, with each donor evaluated independently to avoid confounding from KIR and HLA polymorphism. Statistical analyses: one-way ANOVA with Dunnett’s post-hoc against CD56^dim^CD16^dim^ for panel B; repeated-measures one-way ANOVA with donor matching and Dunnett’s post-hoc for panels C and D. Full ANOVA tables and post-hoc results are in Supplemental Tables 2 and 3. Significance: * p < 0.05, ** p < 0.01, *** p < 0.001, **** p < 0.0001.

### The superiority of CD56^dim^CD16^dim^ to degranulate in response to HIV-infected cells is reproducible and intrinsic

To determine whether the HIV-specific functional superiority of CD56^dim^CD16^dim^ NK cells was reproducible, we examined NK cell degranulation responses to HIV-infected cells at a 1:1 E:T ratio across six healthy, uninfected donors. The NK cell subsets evaluated were based on nine different populations defined by gates set using fluorescence-minus-one (FMO) staining controls for CD56 and CD16, with gates for CD56^bright^CD16^negative^ and CD56^dim^CD16^bright^ NK cells used to determine “bright” NK cell subpopulations, as shown in Figure 1-figure supplement 1 A.

Analysis of NK cell responses to autologous HIV productively infected targets across multiple donors (Figure 1C) confirmed the consistent superiority of CD56^dim^CD16^dim^ NK cells in the present study. The CD56^negative^CD16^dim^ subset was excluded from this analysis because its very low frequency in peripheral blood (<0.2% of NK cells; Figure 1-figure supplement 1B) resulted in insufficient event counts for reliable measurement. Within the CD56^dim^ population, a significantly higher frequency of CD16^dim^ cells showed degranulation responses than CD56^bright^CD16^bright^ (p=0.0110), CD56^bright^CD16^dim^ (p=0.0200), and CD56^bright^CD16^negative^ (p=0.0192) NK cells. In other comparisons, CD56^dim^CD16^dim^ cells showed a significantly higher frequency of degranulating cells than CD56^negative^CD16^bright^ (p=0.0104) and CD56^negative^CD16^negative^ (p=0.0145) cells (repeated-measures one-way ANOVA F(1.941, 9.705) = 17.88, p=0.0006; Dunnett’s multiple comparisons test against CD56^dim^CD16^dim^; Supplemental Table 3).

To specifically examine the role of CD16 expression levels within the CD56^dim^ population, we directly compared degranulation responses across the CD16^bright^, CD16^dim^, and CD16^negative^ subsets. CD56^dim^CD16^dim^ cells exhibited significantly greater HIV-specific degranulation compared to both CD56^dim^CD16^bright^ (p=0.0043) and CD56^dim^CD16^negative^ cells (p=0.0040) (Figure 1D). The mean ± SD frequency of degranulating CD56^dim^CD16^dim^ cells was 11.03 ± 4.51%, which was more than 4-fold higher than the frequency of degranulating CD56^dim^CD16^bright^ cells (2.36 ± 1.44%) and more than 3-fold higher than CD56^dim^CD16^negative^ cells (2.93 ± 2.79%).

### CD56^dim^CD16^dim^ cells demonstrate greater cytotoxicity even when purified

To verify that the observed HIV-specific functional superiority was inherent to CD56^dim^CD16^dim^ NK cells and not influenced by differences in subset frequency in mixed populations, we performed cytotoxicity assays using purified CD56^dim^CD16^bright^ and CD56^dim^CD16^dim^ NK cell subsets.

Direct comparison of purified NK cell subsets revealed that CD56^dim^CD16^dim^ NK cells exhibited significantly higher cytotoxic activity against autologous HIV-infected targets at three of four E:T ratios tested (Figure 2). At the lowest E:T ratio (1:8), CD56^dim^CD16^dim^ cells showed a 2-fold higher mean ± SD of specific lysis (14.49% ± 2.90%) compared to CD56^dim^CD16^bright^ cells (7.50% ± 1.05%). This difference approached statistical significance but did not reach it (p=0.0579). At the 1:4 ratio, the functional advantage of CD56^dim^CD16^dim^ cells reached significance, with 32.63% ± 7.45% specific lysis compared to 16.15% ± 3.16% for CD56^dim^CD16^bright^ cells (p=0.0002). The advantage increased progressively at higher E:T ratios. At the 1:2 ratio, CD56^dim^CD16^dim^ cells achieved 59.96% ± 7.39% specific lysis compared to 26.31% ± 2.18% for CD56^dim^CD16^bright^ cells (p<0.0001). At the 1:1 ratio, CD56^dim^CD16^dim^ cells achieved 77.58% ± 1.35% specific lysis compared to 39.18% ± 2.05% for CD56^dim^CD16^bright^ cells (p<0.0001). The overall analysis confirmed significant effects of NK cell subset (F(1, 16) = 195.0, p<0.0001), E:T ratio (F(3, 16) = 148.1, p<0.0001), and their interaction (F(3, 16) = 18.42, p<0.0001), by two-way ANOVA with Šidák’s multiple comparisons test (Supplemental Table 4).

**Figure 2:**
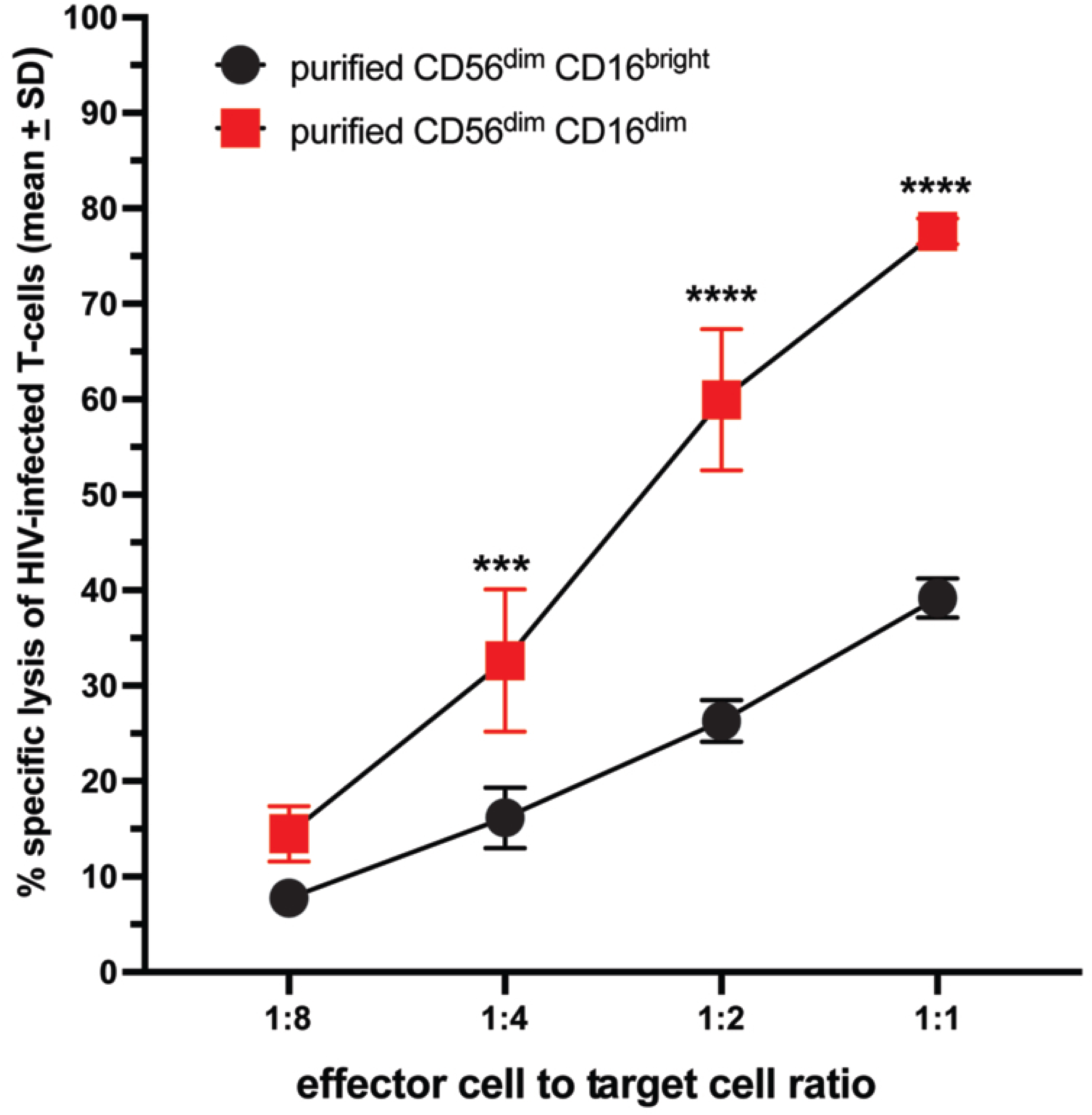
Lysis of autologous HIV-productively infected T-cells by purified CD56^dim^CD16^dim^ and CD56^dim^CD16^bright^ NK cells. Purified NK cells from an uninfected individual were sorted into CD56^dim^CD16^dim^ and CD56^dim^CD16^bright^ NK cell subsets. Carboxyfluorescein succinimidyl ester (CFSE)-labeled purified autologous productively HIV-1_SHM-1_ infected T-cells generated in vitro were added to the sorted NK cell subsets at various effector-to-target cell (E:T) ratios in triplicate. After 4 hours, cells were surface-stained for CD56 and CD3, then exposed to 7-amino-actinomycin D (7AAD). Mean percent specific lysis ± standard deviation (SD) was calculated as (target cell-exposed minus mean unexposed target cells) / (mean positive control minus mean unexposed target cells) × 100, based on 7AAD staining of CFSE-labeled CD3^positive^CD56^negative^ cells. The data shown are from one representative donor and are representative of two independent sort experiments performed with separate donors, each yielding similar results. Statistical analysis: two-way ANOVA (E:T ratio × subset) with Šidák’s multiple comparisons test between subsets at each E:T ratio. Full ANOVA table and post-hoc results are in Supplemental Table 4 for Figure 2. Significance: * p < 0.05, ** p < 0.01, *** p < 0.001, **** p < 0.0001.

### CD56^dim^CD16^dim^ cells exhibit greater degranulation and killing frequency in mixed NK cell populations

To comprehensively characterize the functional differences between CD56^dim^CD16^bright^ and CD56^dim^CD16^dim^ cells within mixed NK cell populations, we analyzed both degranulation responses and killing frequency across a range of E:T ratios.

Analysis of the overall NK cell population revealed distinct patterns of degranulation and cytotoxicity as the E:T ratio increased (Figure 3A). Degranulation of CD56^dim^CD16^positive^ NK cells against autologous HIV-infected cells showed a characteristic decrease from 3.23% ± 0.49% to 1.61% ± 0.05% as E:T ratios increased from 1:4 to 10:1. Specific lysis demonstrated the opposite pattern, rising from 6.23% ± 2.14% to 60.63% ± 1.39% across the same range.

**Figure 3:**
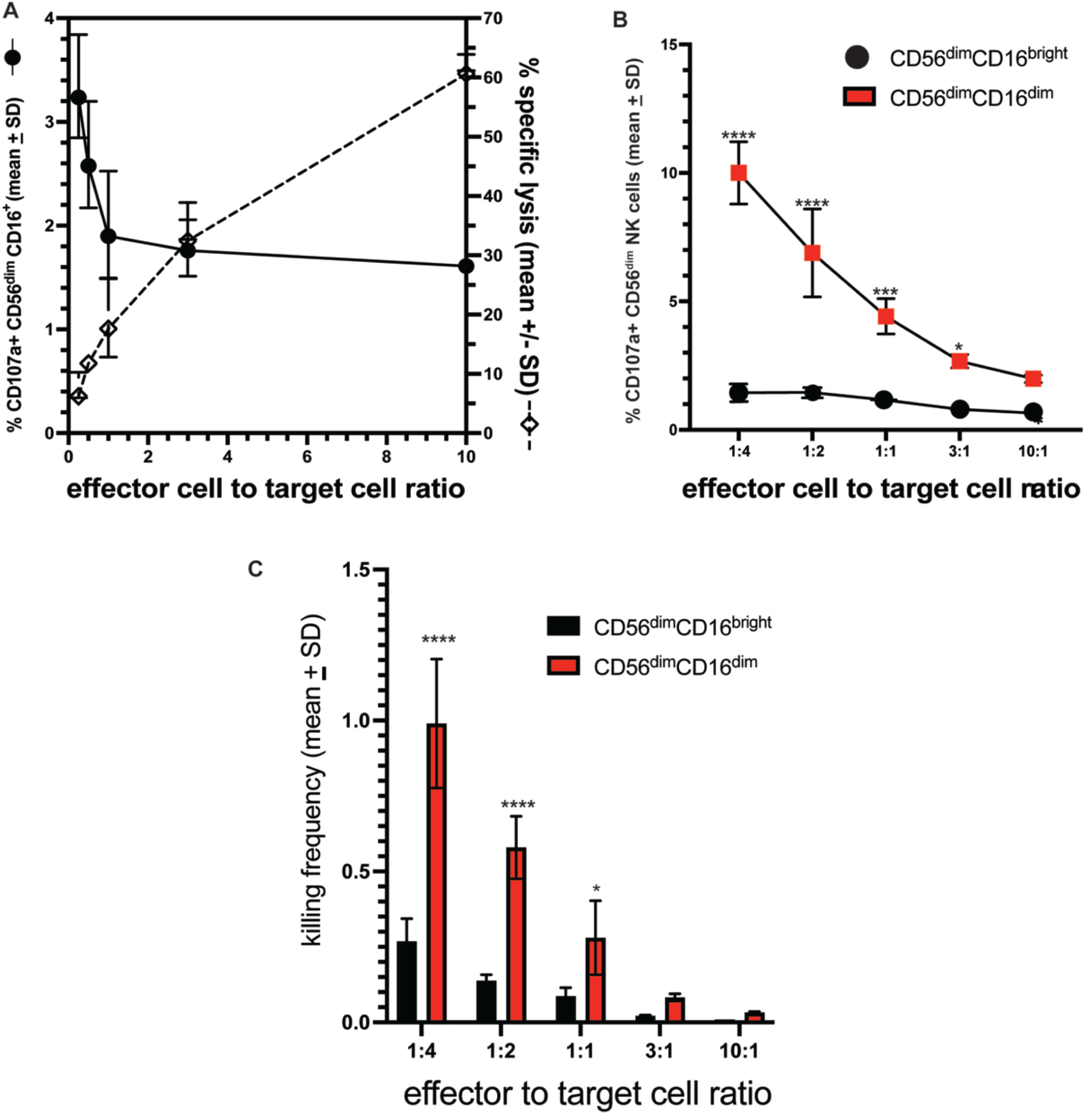
Killing frequency of autologous HIV-productively infected T-cells by CD56^dim^CD16^dim^ and CD56^dim^CD16^bright^ NK cells as they naturally exist within the total NK cell population. (A) Percentage of specific lysis and percentage of CD107a^+^ CD56^dim^CD16^+^ NK cells when exposed to purified autologous HIV-1_SHM-1_ productively infected T-cells across increasing effector cell to target cell (E:T) ratios. Values represent mean ± standard deviation (SD) from three technical replicates. (B) Percentage of CD107a expressing CD56^dim^CD16^bright^ and CD56^dim^CD16^dim^ NK cells when exposed to autologous HIV-1_SHM-1_ productively infected T-cells across increasing E:T ratios. (C) NK cell killing frequency (killing frequency = killed targets / active effectors, where killed targets = total targets × % specific lysis / 100 and active effectors = total effectors × % degranulation / 100) calculated from data in panels A and B. Statistical analyses: two-way ANOVA (E:T ratio × subset) with Šidák’s multiple comparisons test between subsets at each E:T ratio for panels B and C. Full ANOVA tables and post-hoc results are in Supplemental Tables 5 and 6 for Figure 3B and 3C. Significance: * p < 0.05, ** p < 0.01, *** p < 0.001, **** p < 0.0001.

Subset-specific analysis revealed dramatic functional differences between CD56^dim^CD16^bright^ and CD56^dim^CD16^dim^ cells (Figure 3B). A minimal percentage of CD56^dim^CD16^bright^ NK cells degranulated in response to purified productively HIV-infected T-cells (1.57% ± 0.21% to 0.55% ± 0.01%) across all E:T ratios tested. In contrast, a substantially greater percentage of CD56^dim^CD16^dim^ cells degranulated, peaking at the lowest E:T ratio (10.00% ± 1.21% at 1:4; p<0.0001) and decreasing progressively as the E:T ratio increased. The CD56^dim^CD16^dim^ advantage remained significant at 1:2 (p<0.0001), 1:1 (p=0.0001), and 3:1 (p=0.0107), but was no longer significant at 10:1 (1.99% ± 0.15%; p=0.0824). The overall analysis confirmed significant effects of NK cell subset (F(1, 19) = 254.1, p<0.0001), E:T ratio (F(4, 19) = 41.18, p<0.0001), and their interaction (F(4, 19) = 23.94, p<0.0001), by two-way ANOVA with Šidák’s multiple comparisons test (Supplemental Table 5).

Two-way ANOVA of killing frequency showed a significant effect of CD56^dim^ NK cell subsets (F(1,20)=79.47, p<0.0001, 22.20% of variation), a significant effect of E:T ratio (F(4,20)=48.23, p<0.0001, 53.89%), and a significant subset × E:T interaction (F(4,20)=16.39, p<0.0001, 18.32%, Supplemental Table 6). At every E:T ratio tested, CD56^dim^CD16^dim^ cells achieved a higher killing frequency than CD56^dim^CD16^bright^ cells: at 1:4, 0.990 + 0.214 versus 0.269 + 0.075 (mean diff −0.7215, p<0.0001); at 1:2, 0.579 + 0.104 versus 0.138 + 0.020 (mean diff −0.4411, p<0.0001); at 1:1, 0.280 + 0.122 versus 0.088 + 0.027 (mean diff −0.1928, p=0.0147); at 3:1, 0.083 + 0.012 versus 0.022 + 0.002 (mean diff −0.06046, p=0.4124, ns); and at 10:1, 0.033 + 0.003 versus 0.009 + 0.000 (mean diff −0.02375, p=0.7457, ns, Figure 3C, Supplemental Table 6). Within CD56^dim^CD16^bright^ cells, killing frequency at 1:4 was higher than at 3:1 (mean diff 0.2463, p=0.0274) and at 10:1 (mean diff 0.2594, p=0.0181), with no significant differences among other within-subset E:T comparisons (Figure 3C, Supplemental Table 6). Within CD56^dim^CD16^dim^ cells, killing frequency at 1:4 was higher than at 1:2 (mean diff 0.4105, p=0.0001), 1:1 (mean diff 0.7096, p<0.0001), 3:1 (mean diff 0.9073, p<0.0001), and 10:1 (mean diff 0.9571, p<0.0001); killing frequency at 1:2 was higher than at 1:1 (mean diff 0.2991, p=0.0050), 3:1 (mean diff 0.4968, p<0.0001), and 10:1 (mean diff 0.5466, p<0.0001); and killing frequency at 1:1 was higher than at 10:1 (mean diff 0.2476, p=0.0263), with no significant difference between 1:1 vs 3:1 (mean diff 0.1978, p=0.1197) or 3:1 vs 10:1 (mean diff 0.04979, p=0.9990, Figure 3C, Supplemental Table 6).

### In response to HIV-infected cells, CD56^dim^CD16^dim^ NK cells exhibit greater degranulation and serial degranulation than CD56^dim^CD16^bright^ NK cells

Having established that CD56^dim^CD16^dim^ NK cells mediate enhanced killing of HIV-infected targets within mixed NK cell populations, we sought to characterize the kinetics and serial killing capacity of each subset. We measured degranulation over time by exposing purified NK cells to autologous HIV-infected CD4+ T-cells and quantifying CD107a surface expression at multiple time points.

In extended time course experiments (Figure 4-figure supplement 1A), CD56^dim^CD16^dim^ cells demonstrated significantly greater degranulation than CD56^dim^CD16^bright^ cells at 30 minutes, 1 hr., 2 hr., and 4 hr. (all p<0.0001). CD56^dim^CD16^dim^ cells reached peak degranulation at 47% at 1 hour, compared to only 11% for CD56^dim^CD16^bright^ cells (p<0.0001). Degranulation declined in both subsets after the 1 hr. time point, with CD56^dim^CD16^dim^ cells maintaining approximately 31.9% ± 1.0% CD107a positivity at 4 hrs. compared to 5.9% ± 0.29% for CD56^dim^CD16^bright^ cells (p<0.0001) (Figure 4-figure supplement 1A; two-way ANOVA: subset F(1, 20) = 2664, p<0.0001; time F(4, 20) = 463.3, p<0.0001; interaction F(4, 20) = 181.3, p<0.0001; Šidák’s multiple comparisons test; Supplemental Table 7).

Early kinetic analysis (Figure 4-figure supplement 1B) revealed that CD56^dim^CD16^dim^ cells began responding by 30 minutes, reaching approximately 15.6% ± 1.3% degranulation by 1 hour, whereas CD56^dim^CD16^bright^ cells showed minimal early responses, reaching only 2.1% ± 0.4% by 1 hour (p<0.0001). These data indicate that CD56^dim^CD16^dim^ cells respond more rapidly and with greater magnitude to HIV-infected targets (two-way ANOVA: subset F(1, 20) = 438.3, p<0.0001; time F(4, 20) = 203.4, p<0.0001; interaction F(4, 20) = 120.7, p<0.0001; Šidák’s multiple comparisons test; Supplemental Table 8).

To determine whether individual NK cells could sequentially engage multiple targets, we employed a serial degranulation assay that distinguishes cells undergoing zero, one, two, or three degranulation events within 60 minutes (Niemann et al., 2025). This analysis revealed striking differences between the CD16 subsets (Figure 4). The majority of CD56^dim^CD16^bright^ cells (75.9% *±* 1.0%) did not degranulate during the assay period, compared to only 35.2% ± 0.8% of CD56^dim^CD16^dim^ cells (p<0.0001). Among responding cells, the distribution of degranulation events differed markedly across subsets. 9.40% *±* 0.01% of CD56^dim^CD16^bright^ cells exhibited a single degranulation, with only 3.00% ± 0.23% exhibiting two and 3.89% ± 0.54% exhibiting three degranulations.

**Figure 4:**
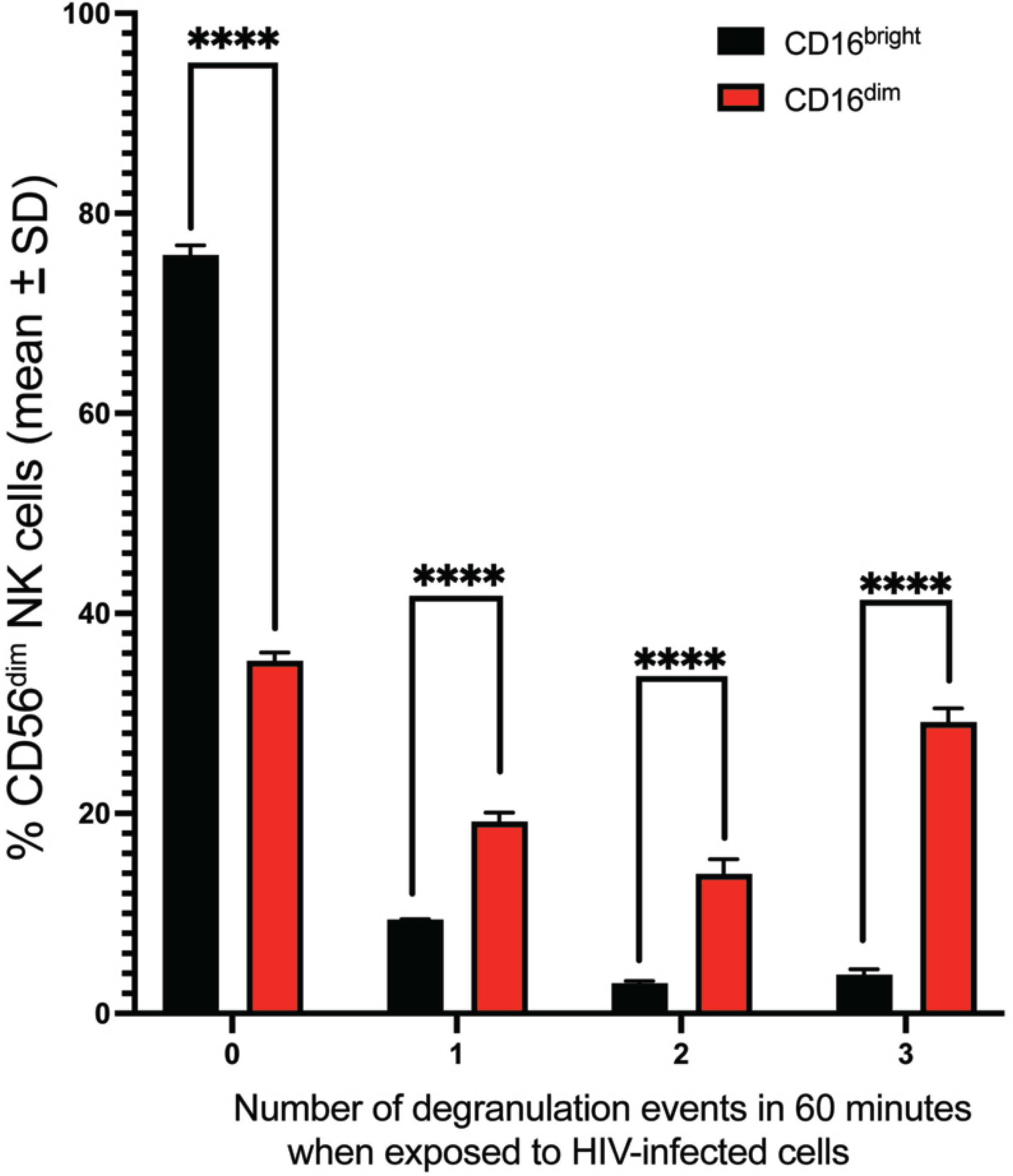
The number of NK cells that degranulated when exposed to HIV-infected cells over a 60-minute period. Purified NK cells exposed to autologous HIV-1_SHM-1_ productively infected T-cells at an effector cell-to-target cell ratio of 1:1 were treated with a different fluorophore-labeled anti-CD107a antibody every 15 minutes for 60 minutes, blocked with excess unlabeled anti-CD107a, and washed between each labeling step. The number of fluorophores accumulated on CD56^dim^CD16^bright^ and CD56^dim^CD16^dim^ NK cells determined how many times each NK cell degranulated during the 60-minute exposure. SD = standard deviation. Statistical analysis: two-way ANOVA (subset × number of degranulation events) with Šidák’s multiple comparisons test between subsets at each event category. Full ANOVA tables and post-hoc results are in Supplemental Table 7 for Figure 4. Significance: * p < 0.05, ** p < 0.01, *** p < 0.001, **** p < 0.0001.

In contrast, CD56^dim^CD16^dim^ cells showed a broader distribution: 19.18% ± 0.89% achieved one degranulation (p<0.0001), 13.94% ± 1.47% achieved two degranulations (p<0.0001), and 29.13% ± 1.36% achieved three degranulations (p<0.0001) (two-way ANOVA: subset F(1, 16) = 12.80, p=0.0025; events F(3, 16) = 3287, p<0.0001; interaction F(3, 16) = 1478, p<0.0001; Šidák’s multiple comparisons test; Supplemental Table 9).

### CD56^dim^CD16^dim^ cells exhibit an enhanced activation state and display HIV-specific noncytolytic responses

To investigate the basis of CD56^dim^CD16^dim^ functional superiority, we examined whether these cells exhibited enhanced HIV-specific non-cytolytic responsiveness compared to CD56^dim^CD16^bright^ cells and baseline activation. CD56^dim^CD16^positive^ NK cells can secrete inflammatory cytokines in addition to lysing target cells (De Maria et al., 2011) and are governed by the same mechanisms that control the lysis of target cells (Bryceson et al., 2006; Mandelboim et al., 1998).

IFN-γ production patterns followed a similar trend to the degranulation response (Figure 1-figure supplement 3). A higher frequency of CD56^dim^CD16^dim^ NK cells produced IFN-γ than CD56^dim^CD16^bright^ NK cells across all E:T ratios tested (Figure 1-figure supplement 3A). CD56^dim^CD16^dim^ cells produced IFN-γ at the 1:4 ratio (4.53 ± 0.58%), 1:2 (3.13 ± 0.60%), 1:1 (2.10 ± 0.51%), and 2:1 (0.87 ± 0.34%) ratios. CD56^dim^CD16^bright^ cells expressed IFN-γ at 1:4 (0.38 ± 0.09%), 1:2 (0.31 ± 0.03%), 1:1 (0.10 ± 0.01%), and 2:1 (0.12 ± 0.19%). Between-subset differences were significant at all E:T ratios (1:4, 1:2, and 1:1, all p<0.0001; 2:1, p=0.0263; two-way ANOVA: subset F(1, 16) = 251.4, p<0.0001; E:T ratio F(3, 16) = 30.08, p<0.0001; interaction F(3, 16) = 21.77, p<0.0001; Šidák’s multiple comparisons test; Supplemental Table 10).

At the same time IFN-γ was measured, we also evaluated the same NK cells for their ability to degranulate (Figure 1-figure supplement 3B). CD56^dim^CD16^dim^ cells degranulated at the 1:4 ratio (15.05 ± 1.70%), 1:2 (11.43 ± 0.91%), 1:1 (7.84 ± 0.30%), and 2:1 (2.56 ± 0.71%) ratios. CD56^dim^CD16^bright^ cells degranulated at 1:4 (2.91 ± 0.16%), 1:2 (1.92 ± 0.06%), 1:1 (1.16 ± 0.11%), and 2:1 (0.37 ± 0.04%). Between-subset differences were significant at all E:T ratios (1:4, 1:2, and 1:1, all p<0.0001; 2:1, p=0.0086; two-way ANOVA: subset F(1, 16) = 649.1, p<0.0001; E:T ratio F(3, 16) = 114.1, p<0.0001; interaction F(3, 16) = 50.49, p<0.0001; Šidák’s multiple comparisons test; Supplemental Table 11).

To determine the baseline activation of CD56^dim^CD16^dim^ NK cell responses, we compared degranulation of CD56^dim^ subsets against uninfected and HIV-infected target cells (Figure 1—figure supplement 3C). At the 1:4 ratio, 21.32 ± 3.78% of CD56^dim^CD16^dim^ cells degranulated against HIV-infected targets and 9.19 ± 1.95% degranulated against uninfected targets. CD56^dim^CD16^dim^ cells responded to HIV-infected cells at 2.3- to 7.3-fold higher percentages than to uninfected cells across all E:T ratios. At the 1:4 ratio, 4.58 ± 0.65% of CD56^dim^CD16^bright^ NK cells degranulated against infected cells and 1.28 ± 0.04% degranulated against uninfected cells [two-way ANOVA: condition F(3, 40) = 329.9, p<0.0001, 67.31% of variation; E:T ratio F(4, 40) = 64.82, p<0.0001, 17.63%; interaction F(12, 40) = 15.13, p<0.0001, 12.34%; Šidák’s multiple comparisons test; Supplemental Table 12].

### CD56^dim^CD16^dim^ NK cell functional superiority is independent of cytotoxic granule protein content and NK cell inhibitory receptors that recognize MHC class I molecules

To determine whether the functional superiority of CD56^dim^CD16^dim^ cells was due to higher levels of cytotoxic effector molecules, we examined the expression of perforin, granzyme A, and granzyme B in the CD56^dim^ NK cell subsets.

Flow cytometric analysis showed that the expression of cytotoxic proteins was broadly similar between CD56^dim^CD16^bright^ and CD56^dim^CD16^dim^ NK cells (Figure 5-figure supplement 1). Perforin was present in nearly all cells of both subsets, with CD56^dim^CD16^bright^ and CD56^dim^CD16^dim^ cells exhibiting similar frequencies of perforin-positive cells (99.99% ± 0.01% and 99.27 ± 0.27% cells expressing perforin, respectively). Likewise, granzyme B expression was consistently high across both subsets (100% of cells expressing granzyme B), indicating that all CD56^dim^ NK cells possess this essential cytotoxic protease, regardless of CD16 levels.

Granzyme A expression followed a similar pattern (Figure 5-figure supplement 1), with both CD56^dim^CD16^bright^ and CD56^dim^CD16^dim^ NK cells showing similar expression frequencies (99.40 ± 0.31 and 96.84 ± 0.54% of cells expressing granzyme A, respectively). In contrast, CD56^dim^CD16^negative^ NK cells had slightly lower perforin (87.04 ± 1.95% of cells expressing perforin) and granzyme A expressing cells (74.96 ± 3.04 % of cells expressing granzyme A), while maintaining high granzyme B expression (100% cells expressing granzyme B).

To determine whether CD56^dim^CD16^dim^ functional superiority resulted from differential KIR3DL1 expression, we evaluated KIR3DL1 frequency and density [geometric mean fluorescent intensity (gMFI)] on CD56^dim^ subsets following 4-hour exposure to purified autologous productively HIV-1_SHM-1_ infected T-cells.

Two-way ANOVA of KIR3DL1^positive^ frequency showed a significant effect of CD16 subset (F(2,18)=54.34, p<0.0001, 77.50% of variation), with no significant effect of E:T ratio (F(2,18)=1.99, p=0.1661) or subset × E:T interaction (F(4,18)=2.39, p=0.0888, Supplemental Table 13). Across the three E:T ratios, KIR3DL1^positive^ frequency was 21.86 ± 0.20% on CD56^dim^CD16^bright^, 20.16 ± 0.89% on CD56^dim^CD16^dim^, and 23.37 ± 0.18% on CD56^dim^CD16^negative:^ cells (Figure 5-figure supplement 2A). At no targets, CD56^dim^CD16^bright^ differed from CD56^dim^CD16^dim^ (mean diff 2.400, p=0.0008), CD56^dim^CD16^bright^ differed from CD56^dim^CD16^negative:^ (mean diff −1.533, p=0.0301), and CD56^dim^CD16^dim^ differed from CD56^dim^CD16^negative:^ (mean diff −3.933, p<0.0001). At the 3:1 E:T ratio, CD56^dim^CD16^bright^ vs CD56^dim^CD16^negative:^ (mean diff −1.700, p=0.0153) and CD56^dim^CD16^dim^ vs CD56^dim^CD16^negative:^ (mean diff −2.167, p=0.0022) differed, while CD56^dim^CD16^bright^ vs CD56^dim^CD16^dim^ did not (mean diff 0.467, p=0.7769). At the 1:1 E:T ratio, CD56^dim^CD16^bright^ vs CD56^dim^CD16^dim^ (mean diff 2.233, p=0.0017) and CD56^dim^CD16^dim^ vs CD56^dim^CD16^negative:^ (mean diff −3.533, p<0.0001) differed, while CD56^dim^CD16^bright^ vs CD56^dim^CD16^negative:^ did not (mean diff −1.300, p=0.0746, Figure 5-figure supplement 2A, Supplemental Table 13).

Two-way ANOVA of KIR3DL1 density showed a significant effect of CD16 subset (F(2,18)=3970, p<0.0001, 98.68% of variation), a significant effect of E:T ratio (F(2,18)=10.70, p=0.0009, 0.27%), and a significant subset × E:T interaction (F(4,18)=16.78, p<0.0001, 0.83%, Supplemental Table 13). Across the three E:T ratios, KIR3DL1 density on KIR3DL1^positive^ cells was 12686.67 ± 107.31 gMFI for CD56^dim^CD16^bright^, 8607.44 ± 203.91 gMFI for CD56^dim^CD16^dim^, and 4630.44 ± 20.36 gMFI for CD56^dim^CD16^negative:^ (Figure 5-figure supplement 2B). At no targets, CD56^dim^CD16^bright^ vs CD56^dim^CD16^dim^ (mean diff 5025, p<0.0001), CD56^dim^CD16^bright^ vs CD56^dim^CD16^negative:^ (mean diff 8146, p<0.0001), and CD56^dim^CD16^dim^ vs CD56^dim^CD16^negative:^ (mean diff 3121, p<0.0001) all differed. The same pattern was observed at the 3:1 E:T ratio (CD56^dim^CD16^bright^ vs CD56^dim^CD16^dim^ mean diff 3666, p<0.0001; CD56^dim^CD16^bright^ vs CD56^dim^CD16^negative:^ mean diff 8004, p<0.0001; CD56^dim^CD16^dim^ vs CD56^dim^CD16^negative:^ mean diff 4338, p<0.0001) and at the 1:1 E:T ratio (CD56^dim^CD16^bright^ vs CD56^dim^CD16^dim^ mean diff 3547, p<0.0001; CD56^dim^CD16^bright^ vs CD56^dim^CD16^negative:^ mean diff 8018, p<0.0001; CD56^dim^CD16^dim^ vs CD56^dim^CD16^negative:^ mean diff 4472, p<0.0001, Figure 5-figure supplement 2B, Supplemental Table 13).

According to NK licensing theory, KIR3DL1^positive^ cells should show enhanced killing when HLA-Bw4 is downmodulated by HIV-1 Nef (Bonaparte & Barker, 2004; Boudreau et al., 2016; Davis et al., 2016). Two-way ANOVA of KIR3DL1-stratified degranulation showed a significant effect of CD16 subset (F(2,12)=80.34, p<0.0001, 78.25% of variation), a significant effect of KIR3DL1 expression (F(1,12)=12.90, p=0.0037, 6.28%), and a significant subset × KIR3DL1 interaction (F(2,12)=9.88, p=0.0029, 9.62%, Supplemental Table 13). Within CD56^dim^CD16^dim^ cells, the frequency of KIR3DL1^positive^ cells that degranulated was higher than the frequency of KIR3DL1− cells that degranulated (8.32 ± 1.54% vs 4.50 ± 0.53%, mean diff 3.812, p=0.0001, Figure 5-figure supplement 2C). No difference was observed within CD56^dim^CD16^bright^ (1.53 ± 0.18% vs 1.63 ± 0.28%, mean diff −0.102, p=0.8822) or CD56^dim^CD16^negative^ (1.11 ± 0.70% vs 0.63 ± 0.89%, mean diff 0.472, p=0.4958, Figure 5-figure supplement 2C). Among KIR3DL1^positive^ cells, CD56^dim^CD16^dim^ cells degranulated at a higher frequency than CD56^dim^CD16^bright^ (mean diff −6.783, p<0.0001) and CD56^dim^CD16^negative^ (mean diff 7.208, p<0.0001) cells, with no difference between CD56^dim^CD16^bright^ and CD56^dim^CD16^negative^ (mean diff 0.426, p=0.9019). Among KIR3DL1− cells, CD56^dim^CD16^dim^ cells also degranulated at a higher frequency than CD56^dim^CD16^bright^ (mean diff −2.870, p=0.0033) and CD56^dim^CD16^negative^ (mean diff 3.869, p=0.0003) cells, with no difference between CD56^dim^CD16^bright^ and CD56^dim^CD16^negative^ (mean diff 1.000, p=0.4135, Figure 5-figure supplement 2C, Supplemental Table 13).

Since in our study we used the HIV-1_SHM-1_ (a primary isolate strain), which does not down-modulate HLA-C (Davis et al., 2016), we wanted to see whether the greater capacity of CD56^dim^CD16^dim^ NK cells to degranulate was due to fewer CD56^dim^CD16^dim^ NK cells expressing KIR2DLs or to lower KIR2DL density than the other CD16 subsets. Two-way ANOVA of KIR2DL2/3^positive^ frequency showed a significant effect of CD16 subset (F(2,18)=54.51, p<0.0001, 81.90% of variation), with no significant effect of E:T ratio (F(2,18)=2.18, p=0.1417) or subset × E:T interaction (F(4,18)=1.18, p=0.7757, Supplemental Table 14). CD56^dim^CD16^bright^ cells had the highest frequency of KIR2DL2/3^positive^ cells (21.04 ± 1.20%) compared to CD56^dim^CD16^dim^ (14.47 ± 0.38%, p=0.0002) and CD56^dim^CD16^negative^ (13.90 ± 1.13%, p<0.0001), with no significant difference between CD56^dim^CD16^dim^ and CD56^dim^CD16^negative^ (p=0.9367, Figure 5-figure supplement 3A, Supplemental Table 14). For KIR2DL2/3 density, two-way ANOVA showed a significant effect of CD16 subset (F(2,18)=58.68, p<0.0001, 84.76% of variation), with no significant effect of E:T ratio (F(2,18)=0.65, p=0.5363) or interaction (F(4,18)=0.45, p=0.7685, Supplemental Table 14). KIR2DL2/3 density was significantly higher on CD56^dim^CD16^bright^ cells (1969.11 ± 55.51 gMFI) compared to CD56^dim^CD16^negative^ (1340.56 ± 20.71 gMFI, p<0.0001), and higher on CD56^dim^CD16^dim^ (1665.11 ± 40.84 gMFI) compared to CD56^dim^CD16^negative^ (p=0.0183), with no significant difference between CD56^dim^CD16^bright^ and CD56^dim^CD16^dim^ (p=0.0804, Figure 5-figure supplement 3A, Supplemental Table 14). Two-way ANOVA of KIR2DL1^positive^ frequency showed a significant effect of CD16 subset (F(2,18)=376.8, p<0.0001, 97.02% of variation), with no significant effect of E:T ratio (F(2,18)=1.40, p=0.2732) or interaction (F(4,18)=0.59, p=0.6753, Supplemental Table 15). KIR2DL1^positive^ cells were most frequent in CD56^dim^CD16^bright^ (4.69 ± 0.33%) versus CD56^dim^CD16^dim^ (1.78 ± 0.27%, p<0.0001) and CD56^dim^CD16^negative^ (0.60 ± 0.10%, p<0.0001) populations, with CD56^dim^CD16^dim^ higher than CD56^dim^CD16^negative^ (p=0.0019, Figure 5-figure supplement 4A, Supplemental Table 15). For KIR2DL1 density, two-way ANOVA showed a significant effect of CD16 subset (F(2,18)=151.7, p<0.0001, 87.06% of variation), a significant effect of E:T ratio (F(2,18)=5.82, p=0.0112, 3.35%), and a significant subset × E:T interaction (F(4,18)=3.82, p=0.0204, 4.40%, Supplemental Table 15). Densities were comparable between CD56^dim^CD16^bright^ (2121.11 ± 10.41 gMFI) and CD56^dim^CD16^dim^ (2097.45 ± 38.66 gMFI, p=0.1517) when expressing KIR2DL1, with both higher than CD56^dim^CD16^negative^ (1793.22 ± 17.39 gMFI, p<0.0001 for both comparisons, Figure 5-figure supplement 4B, Supplemental Table 15).

Functionally, two-way ANOVA of KIR2DL2/3-stratified degranulation showed a significant effect of CD16 subset (F(2,12)=67.44, p<0.0001, 83.31% of variation), a significant effect of KIR2DL2/3 expression (F(1,12)=6.95, p=0.0217, 4.29%), and a significant subset × KIR2DL2/3 interaction (F(2,12)=4.03, p=0.0457, 4.98%, Supplemental Table 14). The frequency of KIR2DL2/3^positive^ CD56^dim^CD16^dim^ cells degranulating was enhanced compared to the frequency of KIR2DL2/3^negative^ counterparts that degranulated (11.16 ± 1.06% vs 7.64 ± 0.70%, p=0.0023, Figure 5-figure supplement 3C, Supplemental Table 14), while only minimal percentages of CD56^dim^CD16^bright^ (KIR2DL2/3^positive^ 3.65 ± 1.15% vs KIR2DL2/3^negative^ 3.28 ± 0.90%, p=0.6958) and CD56^dim^CD16^negative^ (KIR2DL2/3^positive^ 2.56 ± 1.80% vs KIR2DL2/3^negative^ 2.27 ± 0.77%, p=0.7512) cells degranulated regardless of KIR2DL2/3 status (Figure 5-figure supplement 3C, Supplemental Table 14). For KIR2DL1-stratified degranulation, two-way ANOVA showed a significant effect of CD16 subset (F(2,12)=46.19, p<0.0001, 80.39% of variation), no significant effect of KIR2DL1 expression (F(1,12)=1.03, p=0.3304), and a significant subset × KIR2DL1 interaction (F(2,12)=4.75, p=0.0302, 8.27%, Supplemental Table 15). The frequency of CD56^dim^CD16^negative^ cells expressing KIR2DL1 showed paradoxically higher degranulation than the frequency of KIR2DL1^negative^ cells that degranulated (2.02 ± 0.59% vs 0.11 ± 0.11%, p=0.0105, Figure 5-figure supplement 4C, Supplemental Table 15). No significant differences in frequency of cells degranulating were observed between KIR2DL1^positive^ and KIR2DL1^negatve^ cells within CD56^dim^CD16^bright^ (1.23 ± 0.74% vs 1.31 ± 0.72%, p=0.9063) or CD56^dim^CD16^dim^ (4.50 ± 1.21% vs 5.23 ± 0.82%, p=0.2711) subsets (Figure 5-figure supplement 4C, Supplemental Table 15). Critically, a significantly higher percentage of CD56^dim^CD16^dim^ cells degranulated in response to HIV-infected cells compared to both CD56^dim^CD16^bright^ and CD56^dim^CD16^negative^ populations within each KIR2DL2/3 stratum (KIR2DL2/3^positive^: vs CD16^bright^ p<0.0001, vs CD16^negative^ p<0.0001; KIR2DL2/3^negative^: vs CD16^bright^ p=0.0014, vs CD16^negative^ p=0.0002, Figure 5-figure supplement 3C, Supplemental Table 14) and within each KIR2DL1 stratum (KIR2DL1^positive^: vs CD16bright p=0.0007, vs CD16^negative^ p=0.0057; KIR2DL1^negative^: vs CD16bright p=0.0001, vs CD16^negative^ p<0.0001, Figure 5-figure supplement 4C, Supplemental Table 15).

To determine whether CD56^dim^CD16^dim^ functional superiority resulted from differential NKG2A expression, we evaluated NKG2A frequency and density on CD56^dim^ subsets following 4-hour exposure to autologous HIV-infected cells.

Two-way ANOVA of NKG2A^positive^ frequency showed a significant effect of CD16 subset (F(2,18)=7.362, p=0.0046, 37.33% of variation), with no significant effect of E:T ratio (F(2,18)=0.48, p=0.6248) or subset × E:T interaction (F(4,18)=1.44, p=0.2616, Supplemental Table 16). Across the three E:T ratios, NKG2A^positive^ frequency was 99.54 ± 0.04% on CD56^dim^CD16^bright^, 99.61 ± 0.05% on CD56^dim^CD16^dim^, and 99.80 ± 0.03% on CD56^dim^CD16^negative^ cells (Figure 5-figure supplement 5A). At no targets, CD56^dim^CD16^bright^ differed from CD56^dim^CD16^negative^ (mean diff −0.367, p=0.0199), with no significant differences between CD56^dim^CD16^bright^ and CD56^dim^CD16^dim^ (mean diff −0.167, p=0.4500) or between CD56^dim^CD16^dim^ and CD56^dim^CD16^negative^ (mean diff −0.200, p=0.2997). At the 3:1 E:T ratio, no between-subset comparisons reached significance (CD56^dim^CD16^bright^ vs CD56^dim^CD16^dim^ p=0.4500; CD56^dim^CD16^bright^ vs CD56^dim^CD16^negative^ p=0.2997; CD56^dim^CD16^dim^ vs CD56^dim^CD16^negative^ p=0.9899). At the 1:1 E:T ratio, CD56^dim^CD16^dim^ differed from CD56^dim^CD16^negative^ (mean diff −0.333, p=0.0382), with no significant differences between CD56^dim^CD16^bright^ and CD56^dim^CD16^dim^ (mean diff 0.133, p=0.6265) or between CD56^dim^CD16^bright^ and CD56^dim^CD16^negative^ (mean diff −0.200, p=0.2997, Figure 5-figure supplement 5A, Supplemental Table 16).

Two-way ANOVA of NKG2A density on NKG2A^positive^ cells showed no significant effect of CD16 subset (F(2,18)=1.98, p=0.1676), E:T ratio (F(2,18)=1.07, p=0.3630), or subset × E:T interaction (F(4,18)=0.10, p=0.9804, Supplemental Table 16). Across the three E:T ratios, NKG2A density on NKG2A^positive^ cells was 4046.89 ± 161.10 gMFI for CD56^dim^CD16^bright^, 4403.56 ± 114.74 gMFI for CD56^dim^CD16^dim^, and 4155.44 ± 88.41 gMFI for CD56^dim^CD16^negative^ (Figure 5-figure supplement 5B). No between-subset comparisons reached significance at any E:T ratio (Figure 5-figure supplement 5B, Supplemental Table 16).

Two-way ANOVA of NKG2A-stratified degranulation showed a significant effect of CD16 subset (F(2,12)=25.34, p<0.0001, 78.52% of variation), with no significant effect of NKG2A expression (F(1,12)=0.11, p=0.7470) or subset × NKG2A interaction (F(2,12)=0.88, p=0.4410, Supplemental Table 16). Within each CD16 subset, the frequency of NKG2A^positive^ cells degranulating did not differ from the frequency of NKG2A− cells degranulating (CD56^dim^CD16^bright^ 3.10 ± 1.31% vs 3.81 ± 0.96%, mean diff −0.708, p=0.5963; CD56^dim^CD16^dim^ 9.26 ± 2.83% vs 8.12 ± 0.61%, mean diff 1.133, p=0.4007; CD56^dim^CD16^negative^ 2.09 ± 0.72% vs 3.25 ± 1.94%, mean diff −1.169, p=0.3863, Figure 5-figure supplement 5C). Among NKG2A^positive^ cells, CD56^dim^CD16^dim^ cells degranulated at a higher frequency than CD56^dim^CD16^bright^ (mean diff −6.157, p=0.0015) and CD56^dim^CD16^negative^ (mean diff 7.172, p=0.0004) cells, with no difference between CD56^dim^CD16^bright^ and CD56^dim^CD16^negative^ (mean diff 1.015, p=0.8339).

Among NKG2A− cells, CD56^dim^CD16^dim^ cells also degranulated at a higher frequency than CD56^dim^CD16^bright^ (mean diff −4.316, p=0.0182) and CD56^dim^CD16^negative^ (mean diff 4.870, p=0.0084) cells, with no difference between CD56^dim^CD16^bright^ and CD56^dim^CD16^negative^ (mean diff 0.553, p=0.9666, Figure 5-figure supplement 5C, Supplemental Table 16).

### HIV Vpr induces NKG2D ligands on HIV-infected cells and triggers degranulation responses in autologous CD56^dim^CD16^dim^ NK cells through NKG2D

We and others previously demonstrated that NK cell lysis of HIV-infected cells depends on NKG2D (Cerboni et al., 2007; Ward et al., 2007). To understand how HIV-specific recognition occurs in CD56^dim^ NK cell subsets with differential CD16 expression, we examined whether HIV infection induces the expression of specific ligands recognized by the activating receptor NKG2D. Analysis of NKG2D ligand expression on HIV-infected cells using a soluble NKG2D IgG1 Fc chimeric protein revealed that HIV infection induces expression of these stress-induced molecules (Figure 5A) on the infected cell surface, as previously demonstrated by us and others (Cerboni et al., 2007; Richard et al., 2010; Ward et al., 2009). Uninfected CD4^positive^ T-cells exhibited minimal NKG2D ligand expression, as indicated by overlapping histograms for NKG2D IgG1 Fc chimeric protein staining (blue line) and staining control (red line). Cells infected with wild-type (WT) HIV displayed a rightward shift in NKG2D ligand expression (blue line). Cells infected with HIV lacking the Vpr gene (ΔVpr) had lower NKG2D ligand levels, with histograms resembling those of uninfected cells.

**Figure 5:**
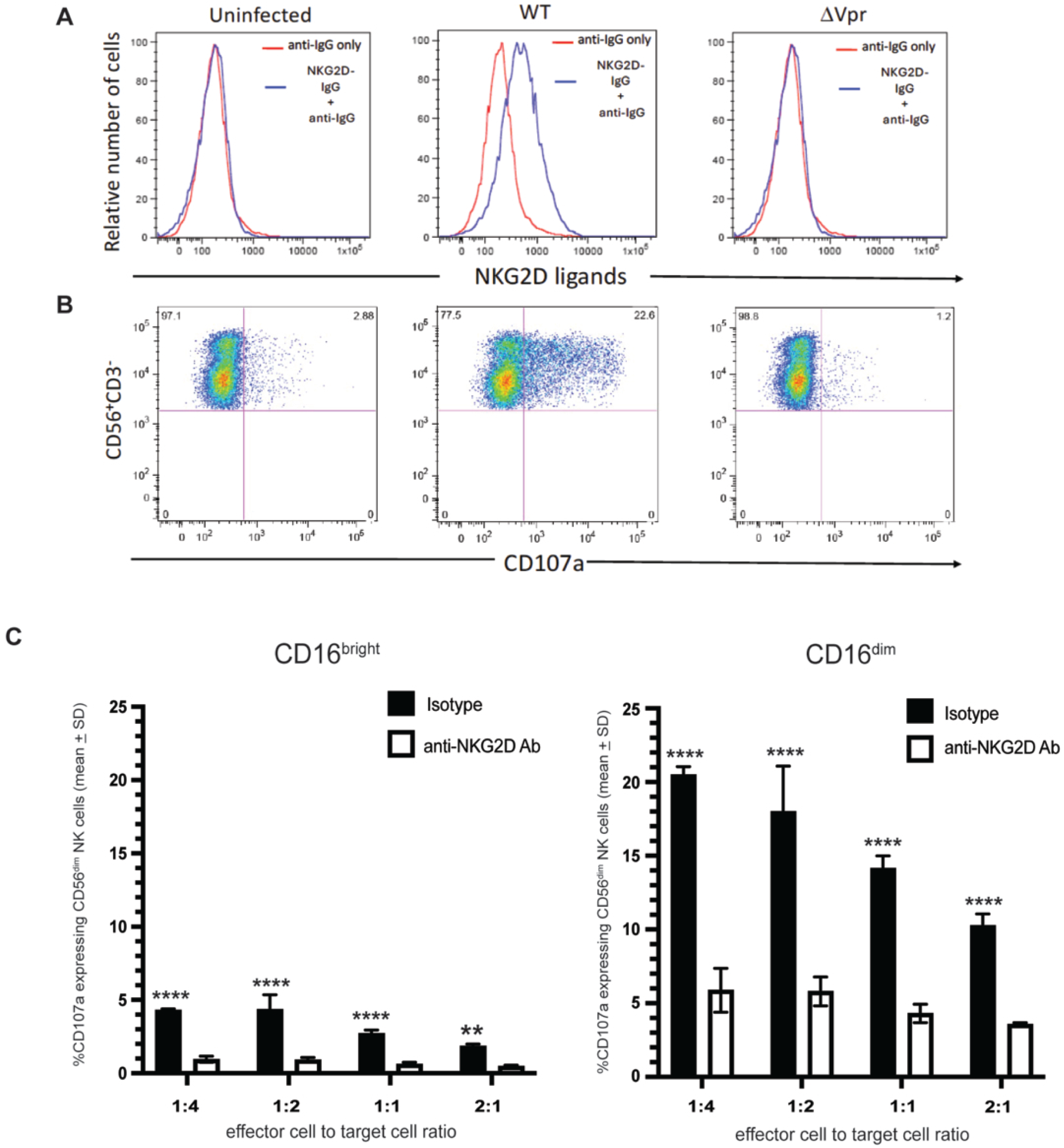
Role of NKG2D in the ability of NK cell subsets to degranulate in response to autologous HIV-productively infected T-cells. (A) CD3 and CD28-stimulated CD4^+^ T-cells remained uninfected or were infected with VSV-G pseudotyped DHIV3, which possessed (wild-type; WT) or lacked (ΔVpr) Vpr. Following infection, cells were stained with a recombinant human soluble NKG2D/IgG1 Fc chimera protein, followed by a fluorochrome-conjugated anti-human IgG1 Fc-specific antibody (Ab). The staining control consisted of cells stained only with the anti-human IgG1 Fc-specific Ab. Cells were then stained with fluorochrome-conjugated antibodies against surface CD4 and intracellular HIV-1 p24 capsid and analyzed by flow cytometry. (B) NK cells were isolated from healthy uninfected donors by negative selection from peripheral blood mononuclear cells and exposed for 4 hours to uninfected CD4^positive^ T-cells or purified HIV-infected cells possessing or lacking Vpr. Cells were stained with fluorochrome-conjugated antibodies against CD56, CD3, and CD107a and analyzed by flow cytometry. Dot plots show CD107a versus CD56 expression on CD3− gated cells, with quadrants set by FMO controls; the upper-right quadrant indicates degranulating (CD107a+) CD56+ NK cells, with the percentage shown. (C) NK cells in triplicate wells were treated with blocking antibodies to NKG2D or an isotype-specific control antibody, then exposed to purified autologous HIV-1_SHM-1_ productively infected T-cells at various effector-to-target (E:T) ratios for 4 hours. Cells were stained with antibodies against CD107a, CD56, CD16, and CD3 and analyzed by flow cytometry. All values represent background-subtracted percentages (target-exposed minus mean unexposed controls). Statistical analysis: separate two-way ANOVAs (E:T ratio × treatment) for CD56^dim^CD16^bright^ and CD56^dim^CD16^dim^ subsets, with Šidák’s multiple comparisons test between isotype and anti-NKG2D treatment at each E:T ratio. Full ANOVA tables and post-hoc results are in Supplemental Table 17 for Figure 5C. Significance: * p < 0.05, ** p < 0.01, *** p < 0.001, **** p < 0.0001.

NK cell degranulation occurred only when NKG2D ligands were present, unlike target T-cells infected with Vpr-lacking HIV, which do not express NKG2D ligands (Figure 5B). Flow cytometry confirmed that NKG2D ligand expression correlated with NK cell degranulation (Figure 5A).

To assess the contribution of NKG2D-mediated recognition to direct killing capacity, CD56^dim^ NK cell subsets defined by CD16 expression were tested against HIV-infected targets across E:T ratios (1:4 to 2:1) in the presence of an NKG2D-blocking Ab (Figure 5C). NKG2D blocking reduced degranulation in CD56^dim^CD16^dim^ cells across all E:T ratios tested: 1:4 (p<0.0001), 1:2 (p<0.0001), 1:1 (p<0.0001), and 2:1 (p<0.0001), with degranulation decreasing from 20.49 ± 0.56% to 5.88 ± 0.98% CD107a^positive^ at the lowest E:T ratio tested (two-way ANOVA: treatment F(1, 16) = 543.0, p<0.0001; E:T ratio F(3, 16) = 36.30, p<0.0001; interaction F(3, 16) = 13.18, p=0.0001; Šidák’s multiple comparisons test; Supplemental Table 17). NKG2D blocking also reduced degranulation in CD56^dim^CD16^bright^ cells across all E:T ratios: 1:4 (p<0.0001), 1:2 (p<0.0001), 1:1 (p<0.0001), and 2:1 (p=0.0018), with responses decreasing from 4.30 ± 0.09% to 0.95 ± 0.21% CD107a^positive^ at the 1:4 E:T ratio (two-way ANOVA: treatment F(1, 16) = 266.4, p<0.0001; E:T ratio F(3, 16) = 21.54, p<0.0001; interaction F(3, 16) = 10.06, p=0.0006; Šidák’s multiple comparisons test; Supplemental Table 17).

### NKG2D expression levels correlate with subset-specific degranulation capacity

To determine whether differential NKG2D receptor expression on CD56^dim^CD16^dim^ and CD56^dim^CD16^bright^ cells could account for the NKG2D-dependent subset-specific degranulation responses to HIV-infected cells, we measured NKG2D surface expression (gMFI) following 4-hour exposure to HIV-infected targets across a range of E:T ratios (Figure 6A). Two-way ANOVA of NKG2D gMFI showed a significant effect of CD56^dim^ NK cell subset (F(1,20)=1897, p<0.0001, 70.97% of variation), a significant effect of E:T ratio (F(4,20)=139.0, p<0.0001, 20.80%), and a significant subset × E:T interaction (F(4,20)=49.98, p<0.0001, 7.48%, Supplemental Table 18). CD56^dim^CD16^dim^ cells showed significantly higher NKG2D gMFI than CD56^dim^CD16^bright^ cells at every E:T ratio tested: at no targets (mean diff −1182, p<0.0001), 1:2 (mean diff −501.0, p<0.0001), 1:1 (mean diff −558.3, p<0.0001), 2:1 (mean diff −679.3, p<0.0001), and 4:1 (mean diff −826.0, p<0.0001, Figure 6A, Supplemental Table 18). Across the five E:T ratios, CD56^dim^CD16^dim^ NKG2D gMFI was 2521.5 ± 17.35, and CD56^dim^CD16^bright^ NKG2D gMFI was 1772.2 ± 9.07.

**Figure 6:**
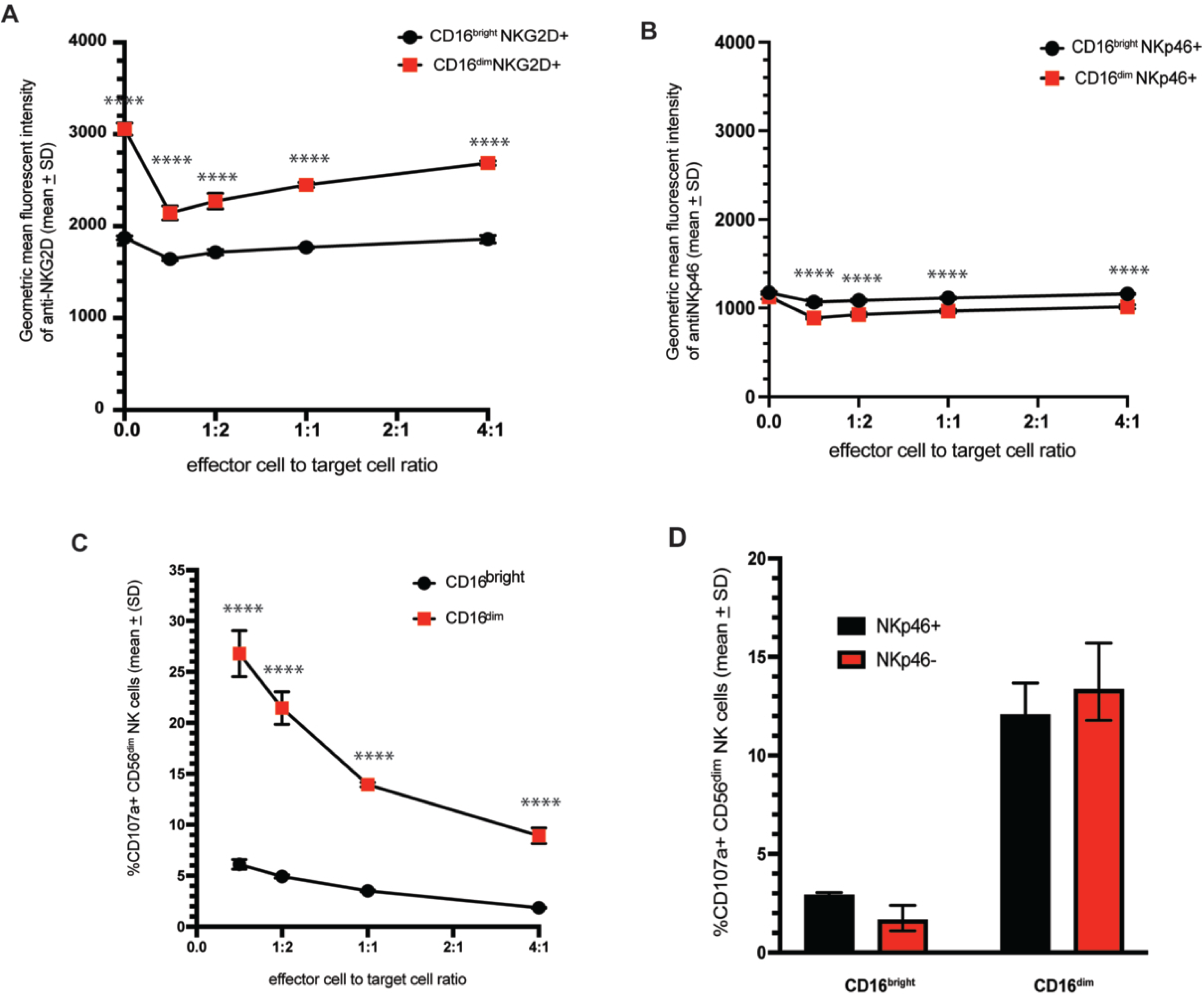
NKG2D and NKp46 surface expression and degranulation responses of CD56^dim^CD16^bright^ and CD56^dim^CD16^dim^ NK cell subsets following HIV-infected T-cell recognition. (A) Geometric mean fluorescence intensity (gMFI) of NKG2D surface expression on CD56^dim^CD16^bright^ and CD56^dim^CD16^dim^ NK cells following 4-hour exposure to purified autologous HIV-1_SHM-1_ productively infected T-cells across different effector cell-to-target cell ratios. (B) gMFI of NKp46 surface expression on the NKp46-positive cells within the CD56^dim^CD16^bright^ and CD56^dim^CD16^dim^ NK cell subsets under the same conditions as (A). (C) CD107a degranulation responses of CD56^dim^CD16^bright^ and CD56d^im^CD16^dim^ NK cells from (A) across different effector cell-to-target cell ratios. (D) CD107a degranulation responses of CD56^dim^CD16^bright^ and CD56^dim^CD16^dim^ NK cells stratified by NKp46 expression (NKp46+ vs NKp46−) at a 1:1 effector-to-target cell ratio. Values represent mean ± standard deviation (SD). All CD107a values represent background-subtracted percentages (target-exposed minus mean unexposed controls). Statistical analyses: two-way ANOVA (effector cell-to-target cell ratio × subset) with Šidák’s multiple comparisons test between subsets at each effector cell-to-target cell ratio for panels A, B, and C; two-way ANOVA (subset × NKp46 status) with Šidák’s multiple comparisons test for panel D. Full ANOVA tables and post-hoc results are in Supplemental Table 18 for Figure 6. Significance: * p < 0.05, ** p < 0.01, *** p < 0.001, **** p < 0.0001.

To verify that the differential NKG2D expression observed between CD16 subsets was receptor-specific and not due to general differences in surface receptor levels, we measured NKp46 expression on the same CD56^dim^CD16^bright^ and CD56^dim^CD16^dim^ NK cells from identical experimental wells (Figure 6B). NKp46 was selected because it does not participate in NK cell lysis of HIV-infected cells (Ward et al., 2007). Two-way ANOVA of NKp46 gMFI showed a significant effect of CD56^dim^ NK cell subset (F(1,20)=321.1, p<0.0001, 50.30% of variation), a significant effect of E:T ratio (F(4,20)=64.89, p<0.0001, 40.66%), and a significant subset × E:T interaction (F(4,20)=9.432, p=0.0002, 5.91%, Supplemental Table 18). In contrast to NKG2D,

NKp46 gMFI did not differ between CD56^dim^CD16^dim^ and CD56^dim^CD16^bright^ cells at no targets (mean diff 45.67, p=0.0668), and showed smaller absolute differences at the higher E:T ratios than those observed for NKG2D (1:2 mean diff 178.3, p<0.0001; 1:1 mean diff 160.7, p<0.0001; 2:1 mean diff 148.3, p<0.0001; 4:1 mean diff 144.3, p<0.0001, Figure 6B, Supplemental Table 18). Across the five E:T ratios, CD56^dim^CD16^bright^ NKp46 gMFI was 1119.7 ± 11.58, and CD56^dim^CD16^dim^ NKp46 gMFI was 984.2 ± 11.83.

As all CD56^dim^CD16^bright^ and CD56^dim^CD16^dim^ NK cells expressed NKG2D receptors, we examined the proportion of cells within each subset that degranulated after exposure to HIV-infected targets (Figure 6C). Two-way ANOVA of % CD107a^positive^ cells showed a significant effect of CD56^dim^ NK cell subset (F(1,16)=1434, p<0.0001, 65.24% of variation), a significant effect of E:T ratio (F(3,16)=179.7, p<0.0001, 24.53%), and a significant subset × E:T interaction (F(3,16)=69.57, p<0.0001, 9.50%, Supplemental Table 18). At every E:T ratio tested, a higher percentage of CD56^dim^CD16^dim^ cells degranulated than CD56^dim^CD16^bright^ cells: at 1:4, 26.53 ± 1.67% of CD56^dim^CD16^dim^ cells degranulated compared to 6.11 ± 0.47% of CD56^dim^CD16^bright^ cells (mean diff −20.42, p<0.0001); at 1:2, 21.46 ± 1.59% versus 4.92 ± 0.17% (mean diff −16.54, p<0.0001); at 1:1, 13.96 ± 0.17% versus 3.51 ± 0.08% (mean diff −10.45, p<0.0001); and at 2:1, 8.91 ± 0.78% versus 1.86 ± 0.05% (mean diff −7.047, p<0.0001, Figure 6C, Supplemental Table 18). Both subsets showed decreasing degranulation frequency as E:T ratios increased. To evaluate whether NKp46 expression affects the differential degranulation capacities observed among CD16 subsets, we analyzed degranulation responses according to NKp46 status within each CD56^dim^ subset at the 1:1 E:T ratio (Figure 6D). Two-way ANOVA showed a significant effect of CD56^dim^ NK cell subset (F(1,8)=176.2, p<0.0001, 94.96% of variation), with no significant effect of NKp46 expression status (F(1,8)=0.963, p=0.3552) or subset × NKp46 interaction (F(1,8)=0.388, p=0.5507, Supplemental Table 18). Within CD56^dim^CD16^bright^ cells, NKp46^positive^ cells exhibited 2.857 ± 0.119% CD107a^positive^ compared to 1.669 ± 0.622% CD107a^positive^ in NKp46− cells, with no significant difference between groups (mean diff 1.188, p=0.4952). Within CD56^dim^CD16^dim^ cells, NKp46^positive^ cells showed 12.227 ± 1.594% CD107a^positive^ compared to 11.961 ± 1.910% in NKp46^negative^ cells, again with no significant difference (mean diff 0.265, p=0.9625, Figure 6D, Supplemental Table 18).

### CD56^dim^CD16^dim^ NK cells exhibit an enhanced antibody-dependent cell-mediated cytotoxic response against anti-gp120-coated HIV-infected cells

To thoroughly evaluate the functional capacity of CD56^dim^CD16^dim^ NK cells, we assessed their ability to mediate ADCC using anti-gp120 broadly neutralizing Ab (clone VRC01)-coated HIV-1∼_SHM-1_ productively infected autologous primary CD4^positive^ T-cells, and compared these responses with the patterns of direct cytotoxicity.

Dose-response analysis of ADCC against anti-gp120 Ab-coated targets at a 1:1 E:T ratio showed that a higher frequency of CD56^dim^CD16^dim^ NK cells exhibited Ab-mediated degranulation responses across all tested VRC01 concentrations (Figure 7A). Two-way ANOVA showed a significant effect of NK cell subset (F(1,16)=424.0, p<0.0001, 85.91% of variation), a significant effect of VRC01 concentration (F(3,16)=16.74, p<0.0001, 10.18%), and no significant subset × VRC01 interaction (F(3,16)=1.112, p=0.3732, 0.68%, Supplemental Table 19). At 0 µg/mL VRC01, CD56^dim^CD16^dim^ cells degranulated at 12.19 cells ± 0.97% CD107a^positive^ compared to 2.92 + 0.06% CD107a^positive^ for CD56^dim^CD16^bright^ (mean diff −9.268, p<0.0001). At 0.5 µg/mL, CD56^dim^CD16^dim^ cells reached 13.17 ± 1.04% versus 3.99 ± 0.06% for CD56^dim^CD16^bright^ (mean diff −9.174, p<0.0001). At 1 µg/mL, CD56^dim^CD16^dim^ cells reached 15.22 ± 2.19% versus 4.94 ± 0.60% for CD56^dim^CD16^bright^ cells (mean diff −10.29, p<0.0001). At 2 µg/mL, CD56^dim^CD16^dim^ cells reached 17.82 ± 1.86% versus 6.45 ± 0.88% for CD56^dim^CD16^bright^ (mean diff −11.37, p<0.0001, Figure 7A, Supplemental Table 19).

**Figure 7:**
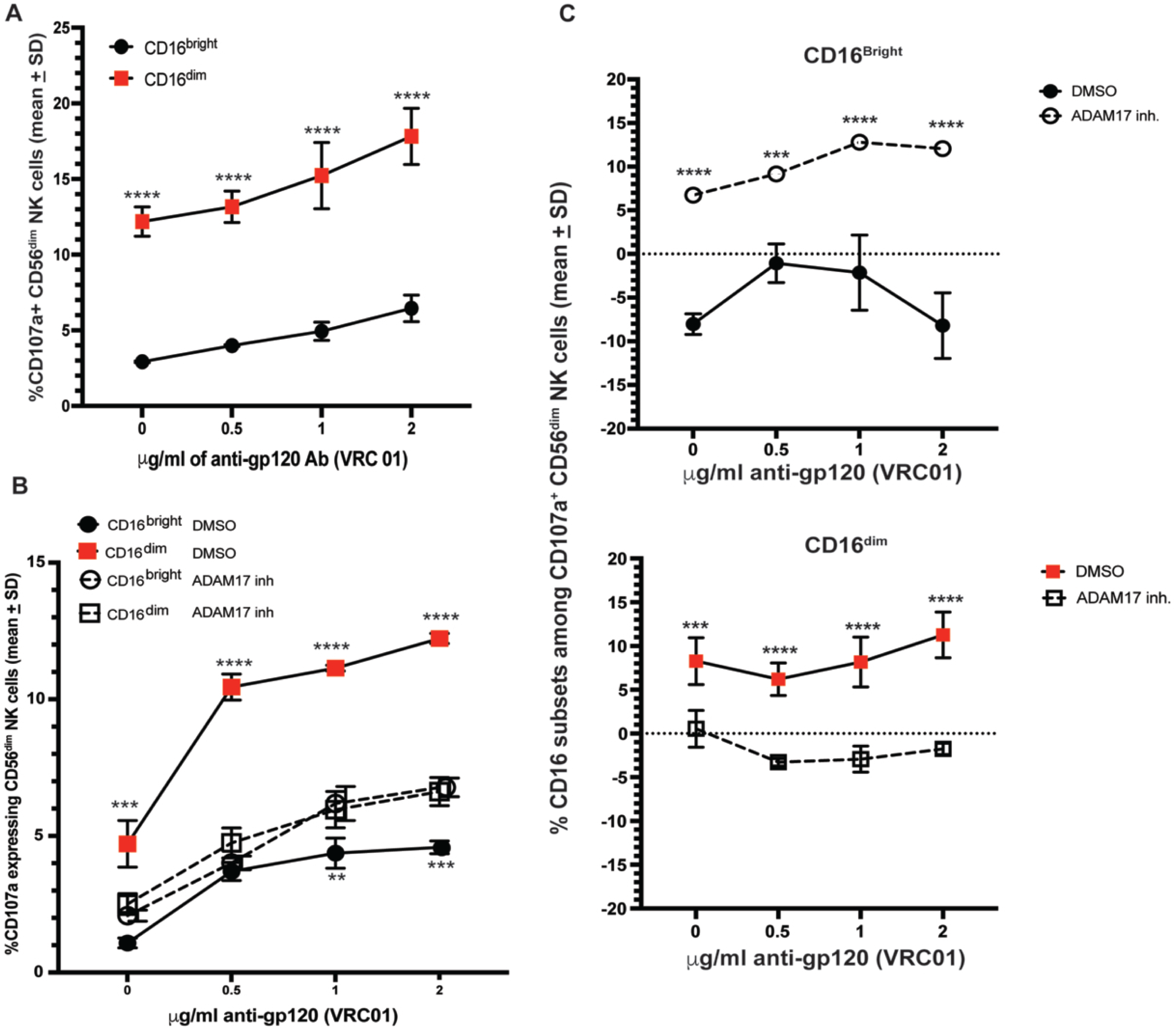
Degranulation of NK cell subsets in response to anti-gp120 labeled autologous HIV productively infected T-cells in the presence or absence of ADAM17 inhibition. (A) Mean percentage ± standard deviation (SD) of CD56dimCD16bright and CD56dimCD16dim NK cells degranulating following 4-hour exposure at an effector-to-target ratio of 1:1 to purified autologous HIV-1SHM-1 productively infected T-cells pretreated with varying concentrations of anti-gp120 broadly neutralizing antibody (VRC01). All values represent background-subtracted percentages (target-exposed minus mean unexposed controls). (B) CD107a degranulation of CD56dimCD16bright (upper sub-panel) and CD56dimCD16dim (lower sub-panel) NK cells following exposure to VRC01-coated autologous HIV-1SHM-1 productively infected T-cells in the presence of an ADAM17 inhibitor in DMSO or DMSO vehicle alone. p-values next to the symbols are comparisons of the vehicle versus an ADAM17 inhibitor for each VRC01 concentration. (C) Change in the percentage of each CD16 subset following 4-hour exposure to purified autologous HIV-1_SHM-1_ productively infected T-cells across VRC01 concentrations, calculated as the percentage of the subset under target exposure minus the percentage of the same subset without target exposure, under either DMSO vehicle or ADAM17 inhibition, shown for CD56^dim^CD16^bright^ (upper sub-panel) and CD56^dim^CD16^dim^ (lower sub-panel) NK cells. p-values next to the symbols are comparisons between the vehicle and the ADAM17 inhibitor at each VRC01 concentration. Statistical analyses: two-way ANOVA (subset × VRC01 concentration) with Šidák’s multiple comparisons test between subsets at each VRC01 concentration for panel A; three-way ANOVA (subset × treatment × VRC01 concentration) with Šidák’s multiple comparisons test between DMSO and ADAM17 inhibition at each VRC01 concentration within each sub-panel for panel B; separate two-way ANOVAs (treatment × VRC01 concentration) for the CD56^dim^CD16^bright^ and CD56^dim^CD16^dim^ sub-panels, with Šidák’s multiple comparisons test between DMSO and ADAM17 inhibition at each VRC01 concentration, for panel C. Full ANOVA tables and post-hoc results are in Supplemental Tables 19, 21, and 22 for Figures 7A, 7B, and 7C. Significance: * p < 0.05, ** p < 0.01, *** p < 0.001, **** p < 0.0001.

To confirm that CD56^dim^CD16^bright^ cells retain the capacity for CD16-mediated degranulation despite their limited response against antibody-coated HIV-infected cells, we tested both subsets under direct CD16 cross-linking conditions using anti-CD16 antibody (clone 3G8). One-way ANOVA across the four treatment conditions (CD56^dim^CD16^bright^ isotype, CD56^dim^CD16^bright^ 3G8, CD56^dim^CD16^dim^ isotype, CD56^dim^CD16^dim^ 3G8) showed a significant effect of treatment (F(3,8)=1887, p<0.0001, R² = 0.9986, Supplemental Table 20). Both subsets demonstrated robust degranulation in response to 3G8, with CD56^dim^CD16^bright^ cell reaching 41.00 ± 2.08% CD107a^positive^ compared to 0.12 ^±^ 0.02% with isotype (mean diff −40.89, p<0.0001) and CD56^dim^CD16^dim^ cells reaching 61.27 ± 0.30% CD107a+ compared to 3.43 ± 1.08% with isotype (mean diff −57.84, p<0.0001). CD56^dim^CD16^dim^ cells degranulated at a higher frequency than CD56^dim^CD16^bright^ cell under 3G8 cross-linking (mean diff −20.27, p<0.0001, Figure 7-figure supplement 1, Supplemental Table 20).

### ADAM17 enhances the ability of CD56^dim^CD16^dim^ NK cells to degranulate in response to anti-HIV-1 gp120 antibody-coated HIV-infected cells

ADAM17 mediates CD16 shedding from the NK cell surface following engagement of antibody-coated targets, thereby influencing the NK cell’s serial killing capacity (Lajoie et al., 2014; Romee et al., 2013). Thus, we investigated how ADAM17’s impact on CD16 shedding affects the function of CD56^dim^CD16^dim^ NK cells against anti-HIV-1 gp120 Ab-labeled HIV-infected targets (Figure 7B). We also determined the extent of changes in the frequency of CD16^bright^ and CD16^dim^ among the CD56^dim^ that degranulated when these cells were exposed to anti-gap120 Ab-coated HIV-infected cells in the absence or presence of ADAM17 inhibition.

Three-way ANOVA of CD107a^positive^ degranulation responses (NK cell subset × treatment × VRC01 concentration) showed a significant effect of NK cell subset (F(1,16)=557.9, p<0.0001, 26.25% of variation), treatment (F(1,16)=178.1, p<0.0001, 7.18%), and VRC01 concentration (F(3,16)=309.0, p<0.0001, 37.38%), as well as a significant treatment × subset interaction (F(1,16)=492.6, p<0.0001, 23.18%), VRC01 × treatment interaction (F(3,16)=11.68, p=0.0003, 1.41%), VRC01 × subset interaction (F(3,16)=8.962, p=0.0010, 1.27%), and VRC01 × treatment × subset interaction (F(3,16)=13.72, p<0.0001, 1.94%, Supplemental Table 21). Within CD56^dim^CD16^bright^ cell, ADAM17 inhibition did not significantly change the percentage of cells degranulating at 0 µg/mL VRC01 (DMSO 1.082 ± 0.181% vs ADAM17 inhibitor 2.084 ± 0.204%, mean diff −1.001, p=0.7204) or at 0.5 µg/mL (DMSO 3.700 ± 0.330% vs ADAM17 inhibitor 4.013 ± 0.251%, mean diff −0.313, p>0.9999), but significantly increased degranulation at 1.0 µg/mL (DMSO 4.371 ± 0.553% vs ADAM17 inhibitor 6.188 ± 0.622%, mean diff −1.817, p=0.0029) and at 2.0 µg/mL (DMSO 4.582 ± 0.231% vs ADAM17 inhibitor 6.776 ± 0.340%, mean diff −2.194, p=0.0002, Figure 7B, Supplemental Table 21). Conversely, within CD56^dim^CD16^dim^ cells, ADAM17 inhibition significantly decreased the percentage of cells degranulating at every VRC01 concentration tested: at 0 µg/mL (DMSO 4.711 ± 0.853% vs ADAM17 i inhibition 2.509 ± 0.378%, mean diff 2.202, p=0.0001), 0.5 µg/mL (DMSO 10.446 ± 0.475% vs ADAM17 inhibition 4.742 ± 0.550%, mean diff 5.704, p<0.0001), 1.0 µg/mL (DMSO 11.141 ± 0.108% vs ADAM17 inhibition 5.959 ± 0.665%, mean diff 5.182, p<0.0001), and 2.0 µg/mL (DMSO 12.226 + 0.183% vs ADAM17 inhibition 6.621 ± 0.516%, mean diff 5.605, p<0.0001, Figure 7B, Supplemental Table 21).

Ab concentration-dependent analysis revealed ADAM17-mediated phenotypic transitions among CD56^dim^ NK cell subsets (Figure 7C). For CD56^dim^CD16^bright^ cells, two-way ANOVA (treatment × VRC01 concentration) showed a significant effect of treatment (F(1,16)=179.7, p<0.0001, 80.93% of variation), a significant effect of VRC01 concentration (F(3,16)=5.437, p=0.0090, 7.35%), and a significant treatment × concentration interaction (F(3,16)=3.338, p=0.0459, 4.51%, Supplemental Table 22).

Under DMSO conditions, the change in CD56^dim^CD16^bright^ frequency upon exposure to Ab-coated targets was negative across the concentration range (−8.03 ± 1.18% at 0 µg/mL, −1.07 ± 2.21% at 0.5 µg/mL, −2.13 ± 4.31% at 1.0 µg/mL, and −8.20 ± 3.76% at 2.0 µg/mL), whereas ADAM17 inhibition shifted these values positive (6.73 ± 3.76%, 9.17 ± 0.17%, 12.80 ± 2.10%, and 12.07 ± 1.73%, respectively). The difference between DMSO and ADAM17 inhibitor was significant at every concentration: 0 µg/mL (mean diff −14.77, p<0.0001), 0.5 µg/mL (mean diff −10.23, p=0.0003), 1.0 µg/mL (mean diff −14.93, p<0.0001), and 2.0 µg/mL (mean diff −20.27, p<0.0001, Figure 7C, Supplemental Table 22).

For CD56^dim^CD16^dim^ cells, two-way ANOVA showed a significant effect of treatment (F(1,16)=156.3, p<0.0001, 82.96% of variation), a significant effect of VRC01 concentration (F(3,16)=3.495, p=0.0402, 5.57%), and no significant treatment × concentration interaction (F(3,16)=1.871, p=0.1751, Supplemental Table 22). Under DMSO conditions, the change in CD56^dim^CD16^dim^ frequency upon exposure to Ab-coated targets was positive across the concentration range (8.27 ± 2.68% at 0 µg/mL, 6.20 ± 1.85% at 0.5 µg/mL, 8.17 ± 2.85% at 1.0 µg/mL, and 11.27 ± 2.61% at 2.0 µg/mL), whereas ADAM17 inhibition held these values at or below zero (0.53 ± 2.11%, −3.28 ± 0.43%, −2.93 ± 1.49%, and −1.77 ± 0.66%, respectively). The difference between DMSO and ADAM17 inhibitor was significant at every concentration: 0 µg/mL (mean diff 7.733, p=0.0003), 0.5 µg/mL (mean diff 9.483, p<0.0001), 1.0 µg/mL (mean diff 11.10, p<0.0001), and 2.0 µg/mL (mean diff 13.03, p<0.0001, Figure 7C, Supplemental Table 22).

### CD56^dim^CD16^dim^ NK cells require coordinated NKG2D signaling and ADAM17 activity for optimal anti-HIV response during antibody-dependent T-cell cytotoxicity of HIV-1-infected T-cells

To evaluate the functional impact of NKG2D and/or ADAM17 on CD56^dim^ NK cell subsets during ADCC, we analyzed the degranulation responses of CD56^dim^CD16^bright^ and CD56^dim^CD16^dim^ NK cells against HIV-infected T-cells opsonized with 1 µg/mL of anti-gp120 Ab (VRC01) at a 1:1 E:T ratio (Figure 8). Two distinct response patterns were observed among the CD56^dim^ NK cell subsets.

**Figure 8:**
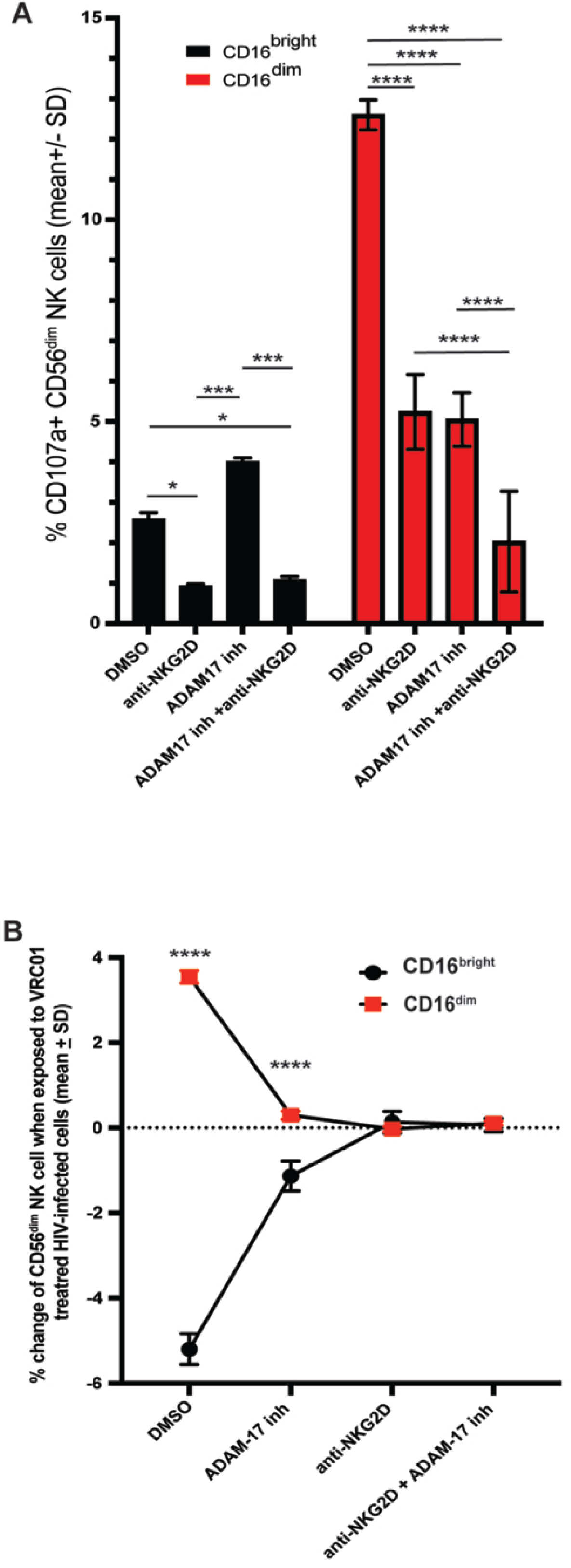
Degranulation of NK cell subsets in response to anti-gp120 labeled autologous HIV productively infected T-cells in the presence or absence of NKG2D blocking antibodies and/or ADAM17 inhibition. (A) Mean percentage ± standard deviation (SD) of CD56^dim^CD16^bright^ and CD56^dim^CD16^dim^ NK cells degranulating following 4-hour exposure at an effector-to-target ratio of 1:1 to purified autologous HIV-1_SHM-1_ productively infected T-cells pretreated with 1 µg/mL of anti-gp120 broadly neutralizing antibody (VRC01). NK cells were treated with DMSO vehicle, anti-NKG2D blocking antibody, ADAM17 inhibitor, or anti-NKG2D blocking antibody combined with ADAM17 inhibitor before being added to VRC01-coated HIV-infected T-cells. All values represent background-subtracted percentages (target-exposed minus mean unexposed controls). (B) Change in the frequency of CD56^dim^CD16^bright^ and CD56^dim^CD16^dim^ NK cells under each treatment condition, calculated as the percentage of the subset following 4-hour exposure to VRC01-coated HIV-infected T-cells minus the percentage of the same subset under matched treatment in the absence of target cells and antibody. Statistical analyses: two-way ANOVA (subset × treatment) with Tukey’s multiple comparisons test for panel A; two-way ANOVA (subset × treatment) with Šidák’s multiple comparisons test for panel B. Full ANOVA tables and post-hoc results are in Supplemental Tables 23 and 24 for Figures 8A and 8B. Significance: * p < 0.05, ** p < 0.01, *** p < 0.001, **** p < 0.0001

Two-way ANOVA of CD107a^positive^ degranulation showed a significant effect of CD56^dim^ NK cell subset (F(1,16)=245.2, p<0.0001, 32.59% of variation), a significant effect of treatment (F(3,16)=97.33, p<0.0001, 38.81%), and a significant subset × treatment interaction (F(3,16)=66.39, p<0.0001, 26.47%, Supplemental Table 23). Under DMSO control conditions, CD56^dim^CD16^dim^ cells degranulated at 12.604 ± 0.376% CD107a^positive^ compared to 2.583 ± 0.163% for CD56^dim^CD16^bright^ cell (mean diff −10.02, p<0.0001, Figure 8A). Within CD56^dim^CD16^dim^ cells, all three perturbations reduced degranulation relative to DMSO: anti-NKG2D blockade reduced degranulation to 5.241 ± 0.926% (mean diff 7.363, p<0.0001), ADAM17 inhibition reduced degranulation to 5.048 ± 0.814% (mean diff 7.556, p<0.0001), and the ADAM17 inhibitor + anti-NKG2D combination reduced degranulation to 2.026 ± 1.249% (mean diff 10.58, p<0.0001). The combination treatment reduced degranulation further than either single perturbation: ADAM17 inhibition vs combination mean diff 3.022, p=0.0001; anti-NKG2D vs combination mean diff 3.215, p<0.0001. Anti-NKG2D and ADAM17 inhibition alone did not differ from each other (mean diff 0.193, p=0.9821, Figure 8A, Supplemental Table 23).

Within CD56^dim^CD16^bright^ cell, the pattern differed. Anti-NKG2D blockade reduced degranulation from 2.583 ± 0.163% to 0.918 ± 0.060% (mean diff 1.666, p=0.0263). ADAM17 inhibition increased degranulation to 3.993 ± 0.113% (mean difference −1.410, p = 0.0679, ns). The ADAM17 inhibitor ± anti-NKG2D combination reduced degranulation to 1.074 ± 0.084% (DMSO vs combination mean diff 1.509, p=0.0472; ADAM17 inhibition vs combination mean diff 2.919, p=0.0002). Anti-NKG2D alone and the combination did not differ (mean diff −0.157, p=0.9903, Figure 8A, Supplemental Table 23). Between subsets, CD56^dim^CD16^dim^ cells degranulated at higher frequencies than CD56^dim^CD16^bright^ cell under DMSO (mean diff −10.02, p<0.0001) and anti-NKG2D (mean diff −4.323, p<0.0001), with no significant difference under ADAM17 inhibition (mean diff −1.055, p=0.0604) or under the combination treatment (mean diff −0.952, p=0.0871, Figure 8A, Supplemental Table 23).

Analysis of the change in CD56^dim^ NK cell subset frequency when exposed to Ab-coated HIV-infected T-cells (relative to no-target controls) revealed activation-induced phenotypic transitions (Figure 8B). Two-way ANOVA showed a significant effect of CD56^dim^ subset (F(1,16)=805.3, p<0.0001, 31.09% of variation), a significant effect of treatment (F(3,16)=23.83, p<0.0001, 2.77%), and a significant subset × treatment interaction (F(3,16)=565.8, p<0.0001, 65.52%, Supplemental Table 24). Under DMSO conditions, CD56^dim^CD16^bright^ frequency decreased by 5.20 ± 0.36% while CD56^dim^CD16^dim^ frequency increased by 3.55 ± 0.15% (between-subset mean diff −8.747, p<0.0001). ADAM17 inhibition reduced these shifts: CD56^dim^CD16^bright^ Δ −1.13 ± 0.35% and CD56^dim^CD16^dim^ Δ 0.30 ± 0.10% (between-subset mean difference −1.430, p<0.0001). Anti-NKG2D blockade and the anti-NKG2D + ADAM17 inhibitor combination effectively abolished the shifts: under anti-NKG2D, CD56^dim^CD16^bright^ Δ 0.13 ± 0.25% and CD56^dim^CD16^dim^ Δ −0.03 ± 0.06% (between-subset mean diff 0.163, p=0.3703); under the combination, CD56^dim^CD16^bright^ Δ 0.07 ± 0.15% and CD56^dim^CD16^dim^ Δ 0.11 ± 0.05% (between-subset mean diff −0.043, p=0.8099, Figure 8B, Supplemental Table 24).

Within CD56^dim^CD16^bright^ cell, the change in subset frequency differed significantly between DMSO and each of the three perturbations: DMSO vs ADAM17 inhibition (mean diff −4.067, p<0.0001), DMSO vs anti-NKG2D (mean diff −5.333, p<0.0001), DMSO vs anti-NKG2D + ADAM17 inhibition (mean diff −5.267, p<0.0001). ADAM17 inhibition also differed from anti-NKG2D (mean diff −1.267, p<0.0001) and from the combination (mean diff −1.200, p<0.0001), with no significant difference between anti-NKG2D and the combination (mean diff 0.067, p=0.9994, Figure 8B, Supplemental Table 24). Within CD56^dim^CD16^dim^ cells, DMSO differed significantly from all three perturbations: vs ADAM17 inhibition (mean diff 3.250, p<0.0001), vs anti-NKG2D (mean diff 3.577, p<0.0001), vs anti-NKG2D + ADAM17 inhibition (mean diff 3.437, p<0.0001), with no significant differences among the three perturbations themselves (Figure 8B, Supplemental Table 24).

### CD56^dim^CD16^dim^ NK cells perform serial ADCC against anti-gp120-coated HIV-infected cells via ADAM17-dependent CD16 turnover

To determine whether CD56^dim^CD16^dim^ NK cells could engage in repeated degranulation against anti-gp120 antibody-coated HIV-infected T-cells, and whether this serial ADCC required ADAM17-mediated CD16 turnover, we employed sequential CD107a labeling at 15-minute intervals over 60 minutes against VRC01-coated autologous HIV-infected T-cells. NK cells were treated with either DMSO vehicle control or an ADAM17 inhibitor before exposure to anti-gp120 antibody-coated targets, and the number of degranulation events per cell within the 60-minute window was quantified.

For CD56^dim^CD16^bright^ NK cells, two-way ANOVA (treatment × number of degranulations) showed a significant effect of the number of degranulations (F(3,16)=6145, p<0.0001, 99.60% of variation), with no significant effect of treatment (F(1,16)=0.13, p=0.7233) and a significant degranulations × treatment interaction (F(3,16)=19.50, p<0.0001, 0.32%, Supplemental Table 25). Under DMSO conditions, 91.06 ± 0.68% of CD56^dim^CD16^bright^ cells failed to degranulate, 5.17 ± 0.20% degranulated once, 3.10 ± 0.36% degranulated twice, and 0.79 ± 0.18% degranulated three times. Under ADAM17 inhibition, the distribution shifted to 84.48 ± 2.34% non-degranulating cells, 6.64 ± 0.07% single events, 4.65 ± 1.54% double events, and 5.12 ± 2.26% triple events. Šidák’s comparisons of DMSO vs ADAM17 inhibition at each degranulation count showed a significant difference at 0 degranulations (mean diff 6.581, p<0.0001) and at 3 degranulations (mean diff −4.326, p=0.0009), with no significant differences at 1 degranulation (mean diff −1.471, p=0.1866) or 2 degranulations (mean diff −1.553, p=0.1645, Figure 9 CD56^dim^CD16^bright^ panel, Supplemental Table 25).

**Figure 9:**
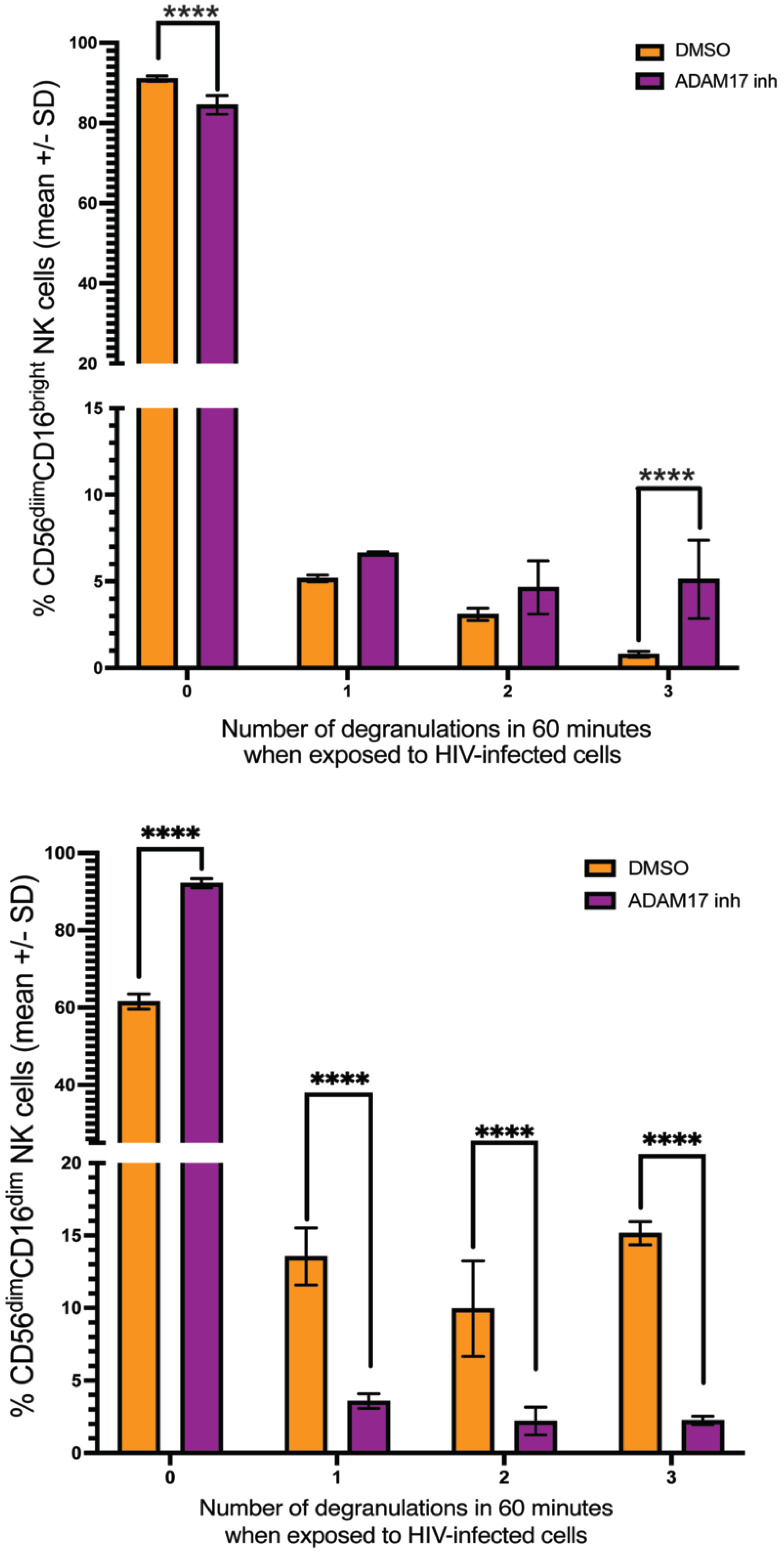
The number of NK cell subsets that degranulated when exposed to anti-gp120 Ab-coated HIV-infected cells over 60 minutes. Purified autologous HIV-1_SHM-1_ productively infected T-cells were coated with 1 µg/mL of the broadly neutralizing anti-gp120 antibody VRC01. NK cells, pretreated with DMSO vehicle or an ADAM17 inhibitor dissolved in DMSO, were exposed to the coated target cells at an effector-to-target cell ratio of 1:1 and treated with a different fluorophore-labeled anti-CD107a antibody every 15 minutes for 60 minutes, with each labeling step blocked by excess unlabeled anti-CD107a and washed between steps. The number of fluorophores accumulated on CD56^dim^CD16^bright^ (upper panel) and CD56d^im^CD16^dim^ (lower panel) NK cells determined how many times each cell degranulated during the 60-minute exposure. SD = standard deviation. Statistical analyses: separate two-way ANOVAs (treatment × number of degranulations) for the CD56^dim^CD16^bright^ panel (with Šidák’s multiple comparisons test) and the CD56^dim^CD16^dim^ panel (with Dunnett’s multiple comparisons test), comparing DMSO and ADAM17 inhibition at each degranulation count. Full ANOVA tables and post-hoc results are in Supplemental Tables 25 and 26 for Figure 9. Significance: * p < 0.05, ** p < 0.01, *** p < 0.001, **** p < 0.0001.

For CD56^dim^CD16^dim^ NK cells, two-way ANOVA showed a significant effect of the number of degranulations (F(3,16)=2619, p<0.0001, 91.74% of variation), no significant effect of treatment (F(1,16)=0.0007, p=0.9788), and a significant degranulations × treatment interaction (F(3,16)=230.6, p<0.0001, 8.08%, Supplemental Table 26). Under DMSO conditions, 61.56 ± 1.94% of CD56^dim^CD16^dim^ cells failed to degranulate, 13.55 ± 1.97% achieved one degranulation, 9.95 ± 3.30% achieved two degranulations, and 15.16 ± 0.80% achieved three degranulations within the 60-minute window. Under ADAM17 inhibition, the population shifted toward non-degranulation: 92.13 ± 1.21% non-degranulating cells, 3.57 ± 0.50% single events, 2.19 ± 0.96% double events, and 2.24 ± 0.30% triple events. Šidák’s comparisons of DMSO vs ADAM17 inhibition at each degranulation count showed significant differences at every level: 0 degranulations (mean diff −30.57, p<0.0001), 1 degranulation (mean diff 9.98, p<0.0001), 2 degranulations (mean diff 7.75, p<0.0001), and 3 degranulations (mean diff 12.92, p<0.0001, Figure 9 CD56^dim^CD16^dim^ panel, Supplemental Table 26).

## Discussion

Cellular immunotherapy has emerged as a promising strategy for HIV functional cure, and its development benefits from identifying which effector cells are most capable of eliminating HIV-infected cells. Knowing which population mediates the dominant response informs cell selection for adoptive transfer. The relevant effector cells for HIV control are likely those that are naturally selected during successful immune containment of the virus. HIV elite controllers maintain undetectable viremia and preserved CD4^positive^ T-cell counts for at least 10 years without antiretroviral therapy, and this control is associated with a reorganization of the CD56^dim^ NK cell compartment, with expansion of the CD56^dim^CD16^dim^ subset and corresponding decrease in the CD56^dim^CD16^bright^ subset compared to people living with HIV on ART and uninfected controls (Rallon et al., 2024). Whether this compartmental reorganization contributes mechanistically to control or simply correlates with it has been unclear. Here, we addressed this question by characterizing the responses of CD56^dim^CD16^dim^ and CD56^dim^CD16^bright^ NK cells to purified autologous productively HIV-infected primary T-cells, a major physiologically relevant target for HIV control. Our findings demonstrate that the minor CD56^dim^CD16^dim^ subset, despite representing less than 6% of circulating NK cells in uninfected subjects, mediates the dominant cytolytic response against HIV-infected cells, whereas the numerically dominant CD56^dim^CD16^bright^ population contributes minimally, even when HIV-infected cells are coated with antibodies that trigger CD16 activation on NK cells. We found that the CD56^dim^CD16^dim^ functional advantage stems from higher NKG2D expression, which is critical for CD56^dim^CD16^dim^ NK cells to mediate both direct killing and ADCC of HIV-infected T-cells, and from a greater capacity for serial killing. The CD56^dim^CD16^dim^ functional advantage is independent of inhibitory KIR repertoire and cytotoxic granule content. These findings provide a mechanistic basis for the compartmental reorganization observed in elite controllers, with expanded CD56^dim^CD16^dim^ cells providing the relevant effector capacity, while the contracted CD56^dim^CD16^bright^ population contributes minimally to HIV-infected cell killing under any condition. Thus, our data support the identification of CD56^dim^CD16^dim^ NK cells as the relevant effector population for cellular therapeutic strategies aimed at achieving functional cure of HIV.

Previous studies (Amand et al., 2017), as well as our own, have shown that the relatively rare CD56^dim^CD16^negative^ cells are more effective against the chronic myeloid leukemia, K562 cell line than other CD56^dim^ subsets (Figure 1-figure supplement 2). Our findings demonstrate that CD56^dim^CD16^dim^ NK cells, comprising ∼3-6% of peripheral blood NK cells and expressing 10-fold lower CD16 levels than CD56^dim^CD16^bright^ cell, are the primary effectors that eliminate HIV-infected T-cells, outperforming both CD56^dim^CD16^bright^ and CD56^dim^CD16^negative^ NK cells (Figure 1).

Despite their low frequency in uninfected subjects, CD56^dim^CD16^dim^ NK cells mediate disproportionate anti-HIV activity. This HIV-specific functional advantage was consistent across multiple donors (Figs. 1C and D), indicating an inherent characteristic rather than donor-specific variation. The killing frequency measurements (Figure 3) confirmed that CD56^dim^CD16^dim^ NK cells achieved significantly higher per-cell killing rates than CD56^dim^CD16^bright^ NK cells, translating enhanced degranulation into effective target elimination. At a 1:4 E:T ratio, one CD56^dim^CD16^dim^ cell could kill one HIV-infected cell, whereas four CD56^dim^CD16^bright^ cells were required for equivalent T-cell killing.

The functional advantage of CD56^dim^CD16^dim^ NK cells against HIV-infected cells emerges in a target system that has not traditionally been a focus of NK cell characterization, where CD56^dim^CD16^dim^ cells have been characterized as transitional or exhausted states (Romee et al., 2013; Yang et al., 2019). Our observations indicate that CD56^dim^CD16^dim^ cells, rather than being transitional or exhausted in the HIV context, drive the optimal NK cell response against HIV-infected cells. At matched effector cell-to-target cell ratios, purified CD56^dim^CD16^dim^ cells also lysed substantially more HIV-infected T-cells than purified CD56^dim^CD16^bright^ cell (Figure 2), confirming through sorted populations what bulk per-cell killing frequency analysis (Figure 3) demonstrated: the cytotoxic advantage is intrinsic to CD56^dim^CD16^dim^ cells rather than a reflection of subset composition. Moreover, CD56d^im^CD16^dim^ cells exhibited lower activation thresholds than CD56^dim^CD16^bright^ cells, evident even in responses to uninfected targets (Figure 1—figure supplement 3C). Notably, CD56^dim^CD16^dim^ cells responding to uninfected targets showed 2.0-fold higher degranulation than CD56^dim^CD16^bright^ cells at a 1:4 E:T ratio when responding to HIV-infected targets, indicating a naturally primed state. Furthermore, the 2.3- to 7.3-fold higher responses of CD56^dim^CD16^dim^ against infected versus uninfected targets indicate pathogen-specific recognition rather than generalized hyperreactivity.

We found in our study that CD56^dim^CD16^dim^ NK cells not only degranulate 4-6 times more frequently than CD56^dim^CD16^bright^ NK cells, but also respond to HIV-infected cells multiple times within one hour. The data presented in Figure 4 reveal two distinct response patterns in CD56^dim^CD16^bright^ cells. First, the majority of CD56^dim^CD16^bright^ cells did not initiate degranulation despite exposure to HIV-infected targets. Second, among the minority of CD56^dim^CD16^bright^ cells that do respond, most achieve only a single degranulation event and fail to continue killing. In contrast, the majority of CD56^dim^CD16^dim^ cells not only respond to HIV-infected targets but also undergo multiple rounds of degranulation, with approximately 15% achieving three degranulation events within one hour. Collectively, approximately 25% of CD56^dim^CD16^dim^ cells achieved two or more degranulations, compared with only approximately 4% of CD56^dim^CD16^bright^ cells, demonstrating that CD56^dim^CD16^dim^ cells are more effective serial killers of HIV-infected cells. The greater capacity of CD56^dim^CD16^dim^ NK cells to respond to HIV-infected cells across multiple rounds of degranulation may explain why they are more effective than CD56^dim^CD16^bright^ cells at lysing HIV-infected cells over a given period.

The HIV-specific cytotoxic advantage of CD56^dim^CD16^dim^ cells is not attributable to higher cytotoxic protein content, as CD56^dim^CD16^bright^ and CD56^dim^CD16^dim^ subsets contain similar levels of perforin, granzyme A, and B (Figure 5-figure supplement 1). These functional differences likely stem from enhanced target recognition, serial killing, and more efficient degranulation. The consistently higher frequency of responding CD56^dim^CD16^dim^ cells across degranulation (Figure 1), IFN-γ production (Figure 1-figure supplement 3A and B), and cytotoxicity (Figure 2) measures indicates that the CD56^dim^CD16^dim^ subset has greater potential for activation against productively HIV-infected cells.

Our profiling of inhibitory receptors revealed an unexpected paradox that argues against differential inhibition by MHC class I molecules as the mechanism underlying the functional advantage of CD56^dim^CD16^dim^ cells. Despite CD56^dim^CD16^bright^ cells expressing the highest frequency and density of HLA-C-restricted KIRs, while CD56^dim^CD16^negative^ cells express the lowest, both subsets exhibit equivalently poor degranulation compared to CD56^dim^CD16^dim^ cells (Figure 5-figure supplement 3A and Figure 5-figure supplement 4A). This functional hierarchy of CD56^dim^CD16^dim^ > CD56^dim^CD16^negative^ and CD56^dim^CD16^bright^ persists even when comparing KIR-negative populations (Figure 5-figure supplement 3C and Figure 5-figure supplement 4C), indicating that inhibitory receptor expression does not account for the observed differences. Further evidence against differential inhibition as the mechanism comes from our analysis of KIR3DL1-mediated education (Kim et al., 2005). When HLA-A and - B molecules containing the HLA-Bw4 serological epitope are downmodulated by HIV Nef (Bonaparte & Barker, 2004; Boudreau et al., 2016; Schwartz et al., 1996), removing KIR3DL1-mediated inhibition, only educated KIR3DL1^positive^ CD56^dim^CD16^dim^ cells showed enhanced degranulation compared to their KIR3DL1^negative^ counterparts (Figure 5-figure supplement 2C), consistent with NK cell licensing (Boudreau et al., 2016; Kim et al., 2005). Remarkably, this education-derived advantage was absent in CD56^dim^CD16^bright^ cells (Figure 5-figure supplement 2A): they remained poorly functional despite greater education and disinhibition. NKG2A expression and HLA-E-mediated inhibition showed only minor differences between CD56^dim^ NK cell subsets that do not account for the observed functional differences (Figure 5-figure supplement 5). This is consistent with our previous demonstration that HIV-infected cells present an HLA-E-bound HIV peptide that fails to engage NKG2A/CD94 productively (Davis et al., 2016), effectively neutralizing this inhibitory pathway. These findings collectively demonstrate that NK cell inhibitory receptor signaling is secondary to intrinsic subset-specific properties in determining NK cell effectiveness against HIV-infected cells. A caveat applies to this interpretation. The stratifications in Figure 5-figure supplement 2C through Figure 5-figure supplement 5C each evaluate degranulation according to a single inhibitory receptor at a time (KIR3DL1, KIR2DL2/3, KIR2DL1, or NKG2A), comparing receptor-positive with receptor-negative cells within each CD56^dim^ subset. Because individual NK cells frequently co-express more than one inhibitory receptor, a cell scored as receptor-negative in any one of these analyses may still carry one or more of the others. These single-receptor stratifications therefore cannot resolve the combined inhibitory input of co-expressed receptors, nor can they formally exclude the possibility that specific combinations of inhibitory receptors influence degranulation in ways not captured when each receptor is analyzed in isolation.

We show in our study that HIV-1 Vpr induces expression of ligands for the NK cell activation receptor NKG2D on infected cells, and CD56^dim^CD16^dim^ cells respond more robustly to these stress ligands than CD56^dim^CD16^bright^ cells (Figure 5). The hierarchy of HIV-specific responses was consistent across NKG2D expression analysis (Figure 6) and blocking studies (Figure 5C), confirming that CD56^dim^CD16^dim^ cells were optimized for NKG2D-mediated recognition. While both CD56^dim^CD16^bright^ and CD56^dim^CD16^dim^ subsets show heavy dependence on NKG2D for HIV-specific responses, with NKG2D blockade reducing degranulation by approximately 65-75% in both subsets, the higher baseline activation of CD56^dim^CD16^dim^ cells (Figure 1-figure supplement 3C) renders them the more effective HIV-specific CD56^dim^ NK cell subset. This functional difference correlates with 1.3-1.5 fold higher NKG2D surface expression on CD56^dim^CD16^dim^ versus CD56^dim^CD16^bright^ cells, while NKp46 expression remains equivalent (Figure 6), suggesting that NKG2D receptor density contributes to HIV-specific functional capacity.

Beyond NKG2D, we examined the expression of other activating receptors implicated in NK cell recognition of infected or stressed cells, namely NKp30, NKp44, NKG2C, DNAM-1, 2B4, and NTB-A, across the CD56^dim^ subsets, and none showed a between-subset difference (data not shown) comparable to that seen with NKG2D. This suggests that the elevated NKG2D on CD56^dim^CD16^dim^ cells distinguishes these subsets among the activating receptors surveyed, although expression analysis alone cannot exclude a functional contribution from these receptors.

In our studies, we observed that, despite CD16 being 10-fold less abundant, CD56^dim^CD16^dim^ NK cells mediated ADCC against productively HIV-infected T-cells 3-4 times more effectively than CD56^dim^CD16^bright^ NK cells (Figure 7A). The ADCC advantage of CD56^dim^CD16^dim^ NK cells was dependent on ADAM17. Inhibition of the enzyme significantly decreased ADCC mediated by CD56^dim^CD16^dim^ NK cells (Figs. 7B, 7C). In contrast, CD56^dim^CD16^bright^ NK cells showed increased ADCC when ADAM17 was inhibited, indicating that ADAM17 dampens the ability of CD56^dim^CD16^bright^ NK cells to function. This asymmetric dependence reframes ADAM17 not as a uniform brake on NK cell function but as a context-dependent regulator that enables serial engagement in CD56^dim^CD16^dim^ cells (Fig 9) while constraining sustained activation in CD56^dim^CD16^bright^ cells.

The inverse relationship between CD16 density and NKG2D expression on these subsets further suggests a compensatory mechanism: the higher NKG2D levels on CD56^dim^CD16^dim^ cells (Figure 6) may offset their lower CD16, enabling sustained recognition of HIV-infected cells through dual receptor engagement (Figure 8). This compensation is likely critical given the low surface density of gp120 on HIV-infected primary T-cells (∼6.4×10^2 molecules per cell (Vasiliver-Shamis et al., 2008)) relative to high-density antigens such as CD20 on Raji cells (∼5×10^4 molecules per cell (Lallemand et al., 2017)). Meeting a specific degranulation threshold against HIV-infected cells may therefore require NKG2D engagement regardless of CD16 expression (Figure 7A). Consistent with this, ADAM17 inhibition reduced CD56^dim^CD16^dim^ degranulation even at 0 µg/mL VRC01 (Figure 7B and C), where no antibody is present to engage CD16, suggesting that NKG2D-driven activation triggers ADAM17 activity independently of CD16 engagement and positioning ADAM17 as an enabler of sustained CD56^dim^CD16^dim^ function downstream of NKG2D signaling rather than solely a regulator of antibody-engaged CD16.

Previous studies have shown that ADAM17-mediated CD16 shedding facilitates NK cell detachment for serial killing (Srpan et al., 2018; Zambarda et al., 2025), an outcome that aligns with serial killing optimization models (Anft et al., 2020) and enables effective ADCC followed by rapid detachment via ADAM17-mediated synapse disassembly (Lorenzen et al., 2016). Our data (Figure 9) extend these findings to HIV-infected targets and suggest that CD56^dim^CD16^dim^ cells depend on efficient CD16 shedding to disengage from one target and engage the next. When shedding is blocked, sustained CD16 expression may prolong antibody-target tethering, which can explain why ADAM17 inhibition enhances otherwise modest CD56^dim^CD16^bright^ responses (Figs. 7B and 8A) but collapses CD56^dim^CD16^dim^ serial degranulation. A receptor-level mechanism for this asymmetry is provided by ADAM17 cleavage of CD16 at Valine 158 (Lajoie et al., 2014; Romee et al., 2013; Valipour et al., 2024), which releases CD2 from the central supramolecular activation cluster (cSMAC) of the immunological synapse, where CD2 is held by a cis interaction with CD16 mediated by Leucine 66 on CD16 (Grier et al., 2012). The release of CD2 from this cis interaction likely permits CD2 to move to the corolla of the immune synapse, where it signals 77x more strongly when it engages CD58 on T-cells (Siokis et al., 2021), thereby contributing to sustained activation of CD56^dim^CD16^dim^ cells during serial engagement (Figure 9).

This study has several significant limitations. Our studies primarily used cells from healthy donors in vitro. Validation in clinical cohorts, particularly elite controllers and patients across disease stages, is essential (Amand et al., 2017; Rallon et al., 2024). Most experiments used HIV-1_SHM-1_, a primary clade B isolate, except for Vpr deletion studies (Figs. 4a-b), which used VSV-G pseudotyped DHIV3 to avoid Vpr strain growth disadvantages (Cohen et al., 1990). Validation across non-clade B isolates is crucial. In vitro-to-in vivo translation uncertainty persists, particularly because peripheral NK cells may not reach HIV reservoir sites, such as lymph nodes or the gastrointestinal tract (Estes et al., 2017). Chronic HIV markedly alters NK cell phenotypes and functions (Jost & Altfeld, 2012), potentially limiting the applicability of healthy donor findings described in this study. Despite limitations, our results identify CD56^dim^CD16^dim^ NK cells as specialized anti-HIV effector cells and provide a mechanistic framework for understanding clinical relevance (Amand et al., 2017; Rallon et al., 2024).

An important consideration in interpreting our results is that NK cells were rested overnight in 100 U/mL IL-2 following isolation, as detailed in the Materials and Methods section. This standard practice maintains NK cell viability while also providing activation signals that could normalize dysfunctional markers and reduce exhaustion phenotypes. Indeed, the dramatic functional differences between CD56^dim^CD16 subsets observed after cytokine rest likely underestimate the disparities present in vivo, where dysfunctional cells would not receive this restorative signal.

Our CD107a degranulation assays were performed without the protein transport inhibitor brefeldin A, to preserve physiological intracellular trafficking conditions (Doms et al., 1989) during NK cell activation. This methodological choice preserves the natural dynamics of both degranulation and CD107a recycling during serial target engagement (Cohnen et al., 2013; Li et al., 2011; Liu et al., 2009). The decline in CD107a surface expression observed at higher E:T ratios in our study has been consistently reported across studies, regardless of inhibitor use, indicating that this reflects authentic NK cell functional kinetics (Vanherberghen et al., 2013). Moreover, since we measure intracellular interferon gamma and surface CD107a at the same time, we found that even in the presence of monensin, CD56^dim^CD16^dim^ NK cells display a higher ability to degranulate than CD56^dim^CD16^bright^ NK cells (Figure 1-figure supplement 3).

An important limitation of the present study is that we did not evaluate trafficking, sustainability, or homing of CD56^dim^CD16^dim^ cells to HIV reservoir sites such as mucosal tissues, which would be required for therapeutic translation. Peripheral blood NK cells naturally lack receptors for gut trafficking such as CCR9 and α4β7 (Lachota et al., 2023), and engineering approaches to add these features and to generate memory-like NK cells capable of sustained response without continuous cytokine treatment (Bednarski et al., 2022; Fehniger et al., 1999; Romee et al., 2012) will be necessary to translate the natural anti-HIV configuration of CD56^dim^CD16^dim^ cells into a durable therapeutic platform.

The findings presented here demonstrate that CD56^dim^CD16^dim^ NK cells, a rare population expanded in HIV elite controllers, are the dominant effectors against HIV-infected primary CD4+ T-cells, and that their effectiveness depends on the integration of NKG2D and CD16-ADAM17 signaling. This combination of dominant per-cell cytotoxicity, NKG2D-dependent threshold activation against low-density gp120, and ADAM17-mediated serial engagement provides a mechanistic basis for the correlation between expansion of this subset and the loss of the dominant CD56^dim^CD16^bright^ population in elite controllers, who maintain long-term viral control without ART. More broadly, the observation that different NK cell subsets dominate against different targets, with CD56^dim^CD16^dim^ cells against HIV-infected primary T-cells and CD56^dim^CD16^negative^ cells against K562 (Figure 1-figure supplement 2), suggests that the dominant effector population is not a fixed property of the NK cell compartment but is matched to the activating and inhibitory signal profile of the target. Primary tumors also show substantial heterogeneity in NK cell ligand and antibody target expression (Shembrey et al., 2019), and antigen heterogeneity is recognized as a central challenge for current CAR-NK cell engineering against solid tumors (Qutub et al., 2026). Whether the dual receptor logic operating in CD56^dim^CD16^dim^ cells against HIV-infected T-cells generalizes to other limiting-antigen and heterogeneous-target contexts, including other intracellular pathogens and antigenically heterogeneous tumors, remains an open question that the framework developed here can address.

## Materials and Methods

### Human subjects

All primary cells, including NK cells and CD4^positive^ T-cells, used in this study were isolated from the peripheral blood of healthy HIV-1-seronegative donors following informed written consent obtained in accordance with the Declaration of Helsinki. The Institutional Review Board at Rush University Medical Center approved the study under protocol 20062305-IRB01 (Chicago, IL, USA).

### Cell isolation and NK cell subset purification

NK cells and CD4^positive^ T-cells were isolated from peripheral blood in separate draws as described by Davis et al. (Davis et al., 2011). Briefly, CD4^positive^ T-cells were isolated from PBMCs using positive selection with anti-CD4 magnetic beads (Miltenyi Biotec) and stimulated for 3 days with anti-CD3 and CD28 Ab beads (Miltenyi Biotec) before HIV infection. NK cells were isolated from PBMCs using the NK cell negative selection kit (Miltenyi Biotec) and cultured in RPMI-1640 medium supplemented with 10% FBS, 2mM glutamine, 100U/ml penicillin, and 100 mg/ml of streptomycin (complete media). We then added 100 IU/ml of recombinant human IL-2 (Peprotech) to the cells and allowed them to rest overnight at 37 °C in a 5% CO_2_ incubator (Fisher). In the cytotoxicity assay, CD16 NK cell subsets were sorted for purity using a Beckman Coulter MoFlo Astrios EQ (flow core facility, University of Illinois at Chicago). The purity of CD4^positive^ and NK cells was greater than 97%, and sorted NK cells (the two subsets evaluated in Figure 2) exceeded 99.9% positivity in our studies.

### HIV infection of primary CD4^positive^ T-cells

Anti-CD3 and anti-CD28 Ab coupled to bead-stimulated primary CD4^positive^ T-cells were infected with either a primary isolate (HIV-1_SHM-1_) or an HIV-1 strain lacking the envelope protein (DHIV3). These envelope-defective viruses were pseudotyped with the vesicular stomatitis virus G protein. For Vpr deletion studies (Figs. 5A and B), VSV-G pseudotyped DHIV3 (wild-type and ΔVpr) were used as target cells in the degranulation assay as described (Davis et al., 2011). All other studies used HIV-1_SHM-1._ HIV infection was carried out by spin-inoculation at an MOI of 0.5, as detailed by O’Doherty et al. (O’Doherty et al., 2000). Following infection, cells were cultured at 1.5 × 10^6^/mL in RPMI complete medium supplemented with 200 IU/mL recombinant IL-2 (Peprotech). Cells were split 1:2 every three days with fresh complete medium containing 200 U/ml of IL-2. Between days 7 and 10 post-infection, the infected cells were processed for target cells as described (Davis et al., 2011). Purified infected cells were verified by flow cytometry using anti-CD4 surface staining and intracellular HIV-1 p24 staining (Supplemental Table 27). Representative dot plots (Figure 1-figure supplement 4) showing CD4 and HIV-1 p24 staining of uninfected (left) and HIV-infected (right) primary T-cells. Across all preparations used in this study, the purity of infected T-cells was greater than 92% p24 antigen-positive.

### Functional assays and flow cytometry

The purified infected cells were added to a U-bottom 96-well plate at a final cell count of 5 × 10^4^ cells per well. In some studies, uninfected CD4^positive^ T-cells, cultured alongside infected cells (Figure 1-figure supplement 3), or K562 (ATCC) cells (Figure 1-figure supplement 2) were used as target cells instead of purified HIV-infected cells. In Ab-dependent cell-mediated cytotoxicity assays, the VRC01 clone of an anti-HIV-1 gp120-specific broadly neutralizing Ab (BEI Resources Repository) was used. VRC01 was added to the infected cells at various concentrations and incubated for 30 minutes at room temperature before the addition of NK cells (Figs. 7, 8, and 9) In some cases the NK cells were either pretreated with 10 μg/ml of anti-NKG2D blocking Ab (R&D Systems; Supplemental Table 27) and/or the ADAM17 inhibitor GW280264X (Aobious) suspended in ^dim^ethyl sulfoxide (DMSO) (Supplemental Table 27) and used at a final concentration of 1.74 mM (Figs. 6 and 7). Controls consisted of adding DMSO to the same final volume as the DMSO used to resuspend the ADAM17 inhibitor (Figs. 7 and 8). Following the incubation period with blocking reagents, NK cells were added to the target cells at NK cell E:T ratios ranging from 1:8 to 10:1, depending on the experiment. All groups were performed in triplicate wells at each E:T ratio, with a final volume of 200 µL. For controls of CD107a surface and intracellular IFNg expression, NK cells were added to wells without target cells. For the cytotoxicity controls, infected target-cells were added without NK cells. The 96-well U-bottom plates containing cells were centrifuged at 20 x g for 2 minutes and incubated at 37 °C in a 5% CO_2_ incubator for 4 hours. For intracellular IFNγ expression by NK cells, we added Monensin (Becton Dickinson) to a final concentration of 2.05 mM, 1 hour into the 4-hour incubation (Supplemental Table 27). At the end of the incubation period, the cells were surface-stained with anti-CD56, CD16, and anti-CD3 antibodies, and then further stained for CD107a to assess NK cell surface expression according to the procedures described by Davis et al. (Davis et al., 2011). All Abs used for flow-based assays are listed in Supplemental Table 27. NK cell intracellular expression of IFN-γ (Figure 1-figure supplement 3A) was determined using the methods described by Cogswell et al. (Cogswell et al., 2022). For the cytotoxicity assessment of HIV-infected cells (Figure 2), we followed the manufacturer’s (Abcam) protocol and control cell treatments for the flow-based cytotoxicity assay (reagents listed in Supplemental Table 27), which used 7-amino-actinomycin D (7AAD) to stain infected cells killed by NK cells.

The flow cytometer used in our study was a Becton Dickinson LSR_Fortessa_, equipped with four lasers set up for Violet (405nm), Blue (488 nm), Yellow (561 nm), and Red (637 nm) emission. All samples collected were analyzed using FlowJo Software, Version 10.9.0. (Becton, Dickinson). Cells were stained with live-dead stain (Supplemental Table 27) followed by cell surface staining with fluorochrome-conjugated Ab against CD3, CD56, and CD16 (Supplemental Table 27). In some studies (Figure 1 and Figure 1-figure supplement 1) fluorochrome-conjugated Ab to NKp80 (Supplemental Table 27) was used. Fluorescence-minus-one (FMO) controls were also set up for all stains and fluorochrome-conjugated antibodies. All cells used in the analysis were viable singlet CD3^negative^ lymphocytes (Figure 1-figure supplement 5) except for cytotoxicity assays, where CD3 was used to identify the infected T-cells (Figure 2). Nine NK cell populations were identified based on differential expression of CD16 and CD56 (Figure 1-figure supplement 1). Fluorescence-minus-one (FMO) controls were used to set negative (neg) gates for CD16 and CD56 (Figure 1-figure supplement 1). Anti-NKp80 staining was used as an NK cell marker to identify CD56^negative^ NK cells (Figure 1-figure supplement 1). CD56^bright^ gates were set based on cells in quadrant 3, and CD16^bright^ gates were set based on cells in quadrant 4 (Figure 1-figure supplement 1).

For functional assays, event acquisition varied with the E:T ratio, reflecting the number of NK cells plated per well. For the CD56^dim^CD16^dim^ subset (3-6% of total NK cells), the average event acquisition ranged from 384 events per well at E:T 1:4 to 3,029 events per well at E:T 4:1 (Figure 1-figure supplement 6). At E:T ratios of 1:2 and 1:4, fewer than 1,000 CD56^dim^CD16^dim^ events were acquired per well. Event numbers for each subset reflected their normal frequencies within the CD56^dim^ population, with CD56^dim^CD16^bright^ (∼80-85% of NK cells) yielding 3,974-63,369 mean events collected per well and CD56^dim^CD16neg (1-3%) yielding 283-5,820 mean events collected per well across all E:T ratios.

### Serial degranulation assay

This flow-based serial degranulation assay is based on the protocol by Niemann et al. (Niemann et al., 2025). Briefly, purified autologous HIV-infected T-cell targets were added to V-bottomed 96-well plates at 5 × 10^4^ cells and pelleted before the addition of 2.5 × 10^4^ NK cells. Under ADCC conditions, 1 μg/ml VRC01 antibody is pre-incubated with the target cells for 30 minutes at 4 °C before NK cells are added to the wells. Control wells consisted of NK cells without target cells. All groups were performed in triplicate wells, with a final volume of 50 μL of RPMI-1640, 10% FBS, 2 mM glutamine, 100 U/ml penicillin, and 100 μg/ml streptomycin (complete medium). The 96-well V-bottom plates were then centrifuged at 500 x g for 10 seconds and incubated at 37 °C in a 5% CO_2_ incubator for 10 minutes. Following the incubation step, the pellets were resuspended in 5 μL of anti-CD107a BV421-conjugated antibody (Table 1) and incubated at 37 °C for 5 minutes. This step fluorescently labels all CD107a present on NK cells at the initial 15-minute exposure. Next, we added 5 μL of non-conjugated anti-

CD107a antibody (Table 1), and incubated for another 5 minutes at 37°C. Blocking allows for only newly externalized CD107a molecules to be stained in the subsequent staining. Following the blocking step, the wells were diluted with 100 μL complete medium, and plates were centrifuged at 500 x g for 5 minutes before being resuspended in a final volume of 50 μL of complete medium. The 96-well V-bottom plates were then centrifuged at 500 x g for 10 seconds and incubated at 37 °C in a 5% CO_2_ incubator for 10 minutes. Following the incubation step, the pellets were resuspended in 5 μL of an anti-CD107a FITC-conjugated antibody (Table 1) and incubated at 37 °C for 5 min. Next, we added 5 μL of non-conjugated anti-CD107a antibody (Table 1), and incubated for another 5 minutes at 37°C. Following the blocking step, the wells were diluted with 100 μL complete medium, and plates were centrifuged at 500 x g for 5 minutes before being resuspended in a final volume of 50 μL of complete medium. The 96-well V-bottom plates were then centrifuged at 500 x g for 10 seconds and incubated at 37 °C in a 5% CO_2_ incubator for 10 minutes. Following the incubation step, the pellets were resuspended in 5 μL of an anti-CD107a PE conjugated antibody (Table 1) and incubated at 37 °C for 5 min. The unconjugated anti-CD107a incubation step is unnecessary during the final stage. At the end of the one-hour incubation, the cells were washed with 100 μL of PBS and centrifuged at 500 × g for 5 minutes. The pellet was then stained for viability and surface markers (anti-CD56, CD16, and CD3) as described above under “**Functional assays and flow cytometry**.”

### Statistical Analyses

Statistical analyses were performed using GraphPad Prism version 10.6.1 (GraphPad Software). Data are presented as mean ± standard deviation (SD) with n = 3 per group for all functional studies. This sample size reflects the practical yield of purified, productively HIV-infected primary CD4+ T-cells, donor-to-donor variability in HIV infection efficiency, and the requirement that p24 purity be verified for every functional study before NK cell exposure. A donor-matched design was used throughout to control for between-donor variation in KIR and HLA polymorphism, recent infections, diet, sleep, and stress, which can influence baseline NK cell phenotype and function.

For Figure 1 and Figure 1-figure supplement 1, comparisons across the three CD56^dim^ NK cell subsets (CD56^dim^CD16^bright^, CD56^dim^CD16^dim^, and CD56^dim^CD16^negative^) were performed using one-way ANOVA with Dunnett’s multiple comparisons test for between-subset contrasts. For Figure 1C and Figure 1D, donor-matched comparisons were performed using repeated-measures one-way ANOVA with donor matching. Based on these data and consistent with Rallon et al. (Rallon et al., 2024), subsequent analyses focused on direct comparisons between the CD56^dim^CD16^bright^ and CD56^dim^CD16^dim^ subsets, which show the most pronounced alterations in HIV elite controllers relative to people living with HIV on antiretroviral therapy and uninfected donors. The CD56^dim^CD16^negative^ subset was retained as a comparator for the inhibitory receptor analyses (KIR3DL1, KIR2DL2/3, KIR2DL1, and NKG2A; Figure 5-figure supplement 2 through Figure 5-figure supplement 5).

Two-way ANOVA with Šidák’s multiple comparisons test was used for the majority of analyses involving two independent factors, including Figures 2, 3B, 3C, 4, 5C, 6A, 6B, 6C, 6D, 7A, 7C, 8B, and 9 (CD56^dim^CD16^bright^ and CD56^dim^CD16^dim^ panels), and figure supplements: Figure 4-figure supplement 1A, Figure 4-figure supplement 1B, Figure 1-figure supplement 3A, Figure 1-figure supplement 3B, Figure 1-figure supplement 3C, Figure 5-figure supplement 2A, Figure 5-figure supplement 2C, Figure 5-figure supplement 3A, Figure 5-figure supplement 3B, Figure 5-figure supplement 3C, Figure 5-figure supplement 4A, Figure 5-figure supplement 4B, Figure 5-figure supplement 4C, Figure 5-figure supplement 5A, Figure 5-figure supplement 5B, and Figure 5-figure supplement 5C. Two-way ANOVA with Tukey’s multiple comparisons test was used where all pairwise contrasts among groups were of interest, specifically Figure 5-figure supplement 2B (KIR3DL1 density) and Figure 8A (NKG2D and ADAM17 combined effects on ADCC). One-way ANOVA with Šidák’s multiple comparisons test for preselected pairwise contrasts was used for Figure 7-figure supplement 1 (anti-CD16 3G8 cross-linking). Three-way ANOVA with Šidák’s multiple comparisons test was used for Figure 7B (NK cell subset × treatment × VRC01 concentration).

For all ANOVA analyses, F-statistics, degrees of freedom, percentage of total variation, and adjusted p-values are reported in the Results section and in the corresponding Supplemental Tables (Supplemental Tables 1 through 26). Statistical significance was defined as p < 0.05, with exact adjusted p-values reported throughout. Post-hoc multiple comparisons p-values are reported as adjusted values.

## Acknowledgments

We thank Kerry Campbell, Ph.D., of Fox Chase Cancer Center, for his helpful insights into experimental design throughout this study. We also thank the Flow Cytometry Cores at Rush University and the University of Illinois at Chicago for technical support.

## Author contributions

**William Howell:** Data curation, Formal analysis, Investigation, Methodology, Validation, Writing – original draft.

**Courtney Branch:** Data curation, Formal analysis, Investigation.

**Jeffrey Ward:** Conceptualization, Data curation, Formal analysis, Investigation.

**Zachary Davis:** Conceptualization, Data curation, Formal analysis, Investigation.

**Emma Geatches:** Data curation, Formal analysis, Investigation.

**Edward Barker:** Formal analysis, Funding acquisition, Investigation, Project administration, Supervision, Writing – original draft, Writing – review & editing.

## Data availability

All data generated and analyzed during this study are included in the manuscript, figure supplements, and supplemental statistical tables. Additional raw donor-level data and source files are available from the corresponding author upon reasonable request. Flow cytometry data files will be deposited in FlowRepository upon acceptance.

