## Supplemental Table 1 for "CD56^dim^CD16^dim^ NK cells are the dominant effector cells against HIV-infected primary T-cells"

**Supplemental Table 1 for the statistical analysis of Figure 1—figure supplement 2**

*Degranulation of CD56^dim^ NK cell subsets (i.e., CD16^bright^, CD16^dim^, CD16^negative^) in response to K562 target cells in two HIV-uninfected donors*

Design: One-way ANOVA per donor (three subsets: CD56^dim^CD16^bright^, CD56^dim^CD16^dim^, CD56^dim^CD16^negative^). Post-hoc: Tukey's multiple comparisons test. n=3 technical replicates per subset per donor; 9 values per donor.

**One-way ANOVA omnibus per donor**

| **Donor** | **F (DFn, DFd)** | **P value** | **R²** | **Brown-Forsythe (variance homogeneity)** | **Significance** |
| --- | --- | --- | --- | --- | --- |
| Donor 1 | F (2, 6) = 570.0 | <0.0001 | 0.9948 | p = 0.6814 (ns) | **** |
| Donor 2 | F (2, 6) = 3139 | <0.0001 | 0.9990 | p = 0.7055 (ns) | **** |

**Tukey's multiple comparisons**

| **Comparison** | **Mean diff.** | **95% CI of diff.** | **q** | **DF** | **Adj. P value** |
| --- | --- | --- | --- | --- | --- |
| ***Donor 1 — subset comparisons*** | | | | | |
| CD56^dim^CD16^bright^ vs CD56^dim^CD16^dim^ | −50.82 | −56.92 to −44.73 | 36.19 | 6 | <0.0001 **** |
| CD56^dim^CD16^bright^ vs CD56^dim^CD16^negative^ | −63.29 | −69.38 to −57.19 | 45.07 | 6 | <0.0001 **** |
| CD56^dim^CD16^dim^ vs CD56^dim^CD16^negative^ | −12.46 | −18.56 to −6.37 | 8.88 | 6 | 0.0018 ** |
| ***Donor 2 — subset comparisons*** | | | | | |
| CD56^dim^CD16^bright^ vs CD56^dim^CD16^dim^ | −36.14 | −38.03 to −34.25 | 82.90 | 6 | <0.0001 **** |
| CD56^dim^CD16^bright^ vs CD56^dim^CD16^negative^ | −46.54 | −48.43 to −44.65 | 106.7 | 6 | <0.0001 **** |
| CD56^dim^CD16^dim^ vs CD56^dim^CD16^negative^ | −10.40 | −12.29 to −8.51 | 23.85 | 6 | <0.0001 **** |

*Significance: ** P < 0.01; **** P < 0.0001.*
