## Supplemental Table 2 for "CD56^dim^CD16^dim^ NK cells are the dominant effector cells against HIV-infected primary T-cells"

**Supplemental Table 2 for the statistical analysis of Figure 1B**

*Statistical analysis of CD107a degranulation responses among NK cell subsets defined by differential CD56 and CD16 expression, against purified autologous HIV-1_SHM-1_ productively infected T-cells at a 1:1 effector cell-to-target cell ratio.*

**Table 1. One-way ANOVA summary.**

| **Source of Variation** | **SS** | **DF** | **MS** | **F (DFn, DFd)** | **P value** |
| --- | --- | --- | --- | --- | --- |
| Between subsets (treatment) | 1397 | 6 | 232.9 | F(6, 14) = 85.96 | < 0.0001 |
| Within subsets (residual) | 37.93 | 14 | 2.709 | — | — |
| Total | 1435 | 20 | — | — | — |

*One-way ANOVA was performed in GraphPad Prism v10.6.1 across seven NK cell subsets. n = 21 values (7 subsets × 3 technical replicates). The Brown-Forsythe test confirmed equality of variances across groups (F(6, 14) = 1.284, p = 0.3260). R² = 0.974.*

**Table 2. Dunnett's multiple comparisons test (each subset vs. CD56^dim^CD16^dim^).**

| **Comparison (control vs.)** | **Mean diff.** | **95% CI of diff.** | **Adjusted P value** | **Summary** | **Significant?** |
| --- | --- | --- | --- | --- | --- |
| CD56^dim^CD16^dim^ vs. CD56^bright^CD16^dim^ | 19.91 | 16.00 to 23.83 | < 0.0001 | **** | Yes |
| CD56^dim^CD16^dim^ vs. CD56^bright^CD16^negative^ | 22.58 | 18.67 to 26.49 | < 0.0001 | **** | Yes |
| CD56^dim^CD16^dim^ vs. CD56^dim^CD16^bright^ | 23.31 | 19.39 to 27.22 | < 0.0001 | **** | Yes |
| CD56^dim^CD16^dim^ vs. CD56^dim^CD16^negative^ | 12.28 | 8.37 to 16.19 | < 0.0001 | **** | Yes |
| CD56^dim^CD16^dim^ vs. CD56^negative^CD16^bright^ | 21.16 | 17.25 to 25.08 | < 0.0001 | **** | Yes |
| CD56^dim^CD16^dim^ vs. CD56^negative^CD16^negative^ | 24.84 | 20.93 to 28.75 | < 0.0001 | **** | Yes |

*Significance summary: * p < 0.05, ** p < 0.01, *** p < 0.001, **** p < 0.0001. CD56^dim^CD16^dim^ is the designated group for all comparisons.*
