## Supplemental Table 3 for "CD56^dim^CD16^dim^ NK cells are the dominant effector cells against HIV-infected primary T-cells"

**Supplemental Table 3 for the statistical analysis of Figure 1C**

*Statistical analysis of CD107a degranulation responses among NK cell subsets defined by differential CD56 and CD16 expression, across six independent donors, against purified autologous productively HIV-1_SHM-1_ infected T-cells at a 1:1 effector cell-to-target cell (E:T) ratio. The CD56^negative^CD16^dim^ subset was excluded from this analysis because its very low frequency in peripheral blood (<0.2% of NK cells; Supplemental Fig. 1B) resulted in insufficient event counts for reliable measurement.*

**Table 1. Repeated-measures one-way ANOVA summary.**

| **Source of Variation** | **SS** | **DF** | **MS** | **F (DFn, DFd)** | **P value** |
| --- | --- | --- | --- | --- | --- |
| Between subsets (treatment) | 430.2 | 7 | 61.46 | F(1.941, 9.705) = 17.88 | 0.0006 |
| Between donors (matching) | 88.40 | 5 | 17.68 | F(5, 35) = 5.143 | 0.0012 |
| Residual (random) | 120.3 | 35 | 3.438 | — | — |
| Total | 638.9 | 47 | — | — | — |

*Repeated-measures one-way ANOVA was performed in GraphPad Prism v10.6.1 across seven NK cell subsets, with donor as a matching factor. Geisser-Greenhouse correction was applied (ε = 0.2773). Donor matching was statistically effective (F(5, 35) = 5.143, p = 0.0012), justifying the repeated-measures design. n = 48 values (8 subsets [including CD56^dim^CD16^dim^] × 6 donors). R² = 0.781.*

**Table 2. Dunnett's multiple comparisons test (each subset vs. CD56^dim^CD16^dim^ control).**

| **Comparison (control vs.)** | **Mean diff.** | **95% CI of diff.** | **Adjusted P value** | **Summary** | **Significant?** |
| --- | --- | --- | --- | --- | --- |
| CD56^dim^CD16^dim^ vs. CD56^bright^CD16^bright^ | 8.613 | 2.69 to 14.53 | 0.0110 | * | Yes |
| CD56^dim^CD16^dim^ vs. CD56^bright^CD16^dim^ | 7.848 | 1.64 to 14.06 | 0.0200 | * | Yes |
| CD56^dim^CD16^dim^ vs. CD56^bright^CD16^negative^ | 8.441 | 1.83 to 15.06 | 0.0192 | * | Yes |
| CD56^dim^CD16^dim^ vs. CD56^dim^CD16^bright^ | 8.673 | 3.86 to 13.48 | 0.0043 | ** | Yes |
| CD56^dim^CD16^dim^ vs. CD56^dim^CD16^negative^ | 8.103 | 3.66 to 12.54 | 0.0040 | ** | Yes |
| CD56^dim^CD16^dim^ vs. CD56^negative^CD16^bright^ | 10.18 | 3.27 to 17.08 | 0.0104 | * | Yes |
| CD56^dim^CD16^dim^ vs. CD56^negative^CD16^negative^ | 9.640 | 2.57 to 16.71 | 0.0145 | * | Yes |

*Significance summary: ns, not significant; * p < 0.05, ** p < 0.01, *** p < 0.001, **** p < 0.0001. CD56^dim^CD16^dim^ is the designated control group for all comparisons.*
