## Supplemental Table 4 for "CD56^dim^CD16^dim^ NK cells are the dominant effector cells against HIV-infected primary T-cells"

**Supplemental Table 4 for the statistical analysis of Figure 2**

*Statistical analysis of percent specific lysis of purified autologous HIV-1_SHM-1_ productively infected T-cells by purified CD56^dim^CD16^bright^ and CD56^dim^CD16^dim^ NK cells across four effector-to-target (E:T) cell ratios (1:8, 1:4, 1:2, and 1:1).*

**Table 1. Two-way ANOVA summary.**

| **Source of Variation** | **% of total variation** | **SS** | **DF** | **MS** | **F (DFn, DFd)** | **P value** |
| --- | --- | --- | --- | --- | --- | --- |
| Interaction (E:T × subset) | 7.78 | 969.7 | 3 | 323.2 | F(3, 16) = 18.42 | < 0.0001 |
| E:T ratio | 62.53 | 7796 | 3 | 2599 | F(3, 16) = 148.1 | < 0.0001 |
| NK cell subset | 27.44 | 3421 | 1 | 3421 | F(1, 16) = 195.0 | < 0.0001 |
| Residual | — | 280.7 | 16 | 17.54 | — | — |
| Total | — | 12467 | 23 | — | — | — |

*Two-way ANOVA was performed in GraphPad Prism v10.6.1 with E:T ratio and NK cell subset as factors. n = 24 values (2 subsets × 4 E:T ratios × 3 replicates per condition per ratio).*

**Table 2. Šidák's multiple comparisons test (within each E:T ratio, compare subsets).**

| **E:T ratio** | **Comparison** | **Mean diff.** | **95% CI of diff.** | **Adjusted P value** | **Summary** |
| --- | --- | --- | --- | --- | --- |
| 1:8 | CD16^bright^ vs. CD16^dim^ | −6.99 | −16.58 to 2.60 | 0.2121 | ns |
| 1:4 | CD16^bright^ vs. CD16^dim^ | −16.48 | −26.07 to −6.89 | 0.0008 | *** |
| 1:2 | CD16^bright^ vs. CD16^dim^ | −33.65 | −43.24 to −24.06 | < 0.0001 | **** |
| 1:1 | CD16^bright^ vs. CD16^dim^ | −38.40 | −47.99 to −28.81 | < 0.0001 | **** |

*Significance summary: ns, not significant; * p < 0.05, ** p < 0.01, *** p < 0.001, **** p < 0.0001.*
