## Supplemental Table 5 for "CD56^dim^CD16^dim^ NK cells are the dominant effector cells against HIV-infected primary T-cells"

**Supplemental Table 5 for the statistical analysis of Figure 3B**

*Statistical analysis of CD107a expression on CD56^dim^CD16^bright^ and CD56^dim^CD16^dim^ NK cells against purified autologous HIV-1_SHM-1_ productively infected T cells across five effector cell-to-target cell (E:T) ratios (1:4, 1:2, 1:1, 3:1, and 10:1).*

**Table 1. Two-way ANOVA summary.**

| **Source of Variation** | **% of total variation** | **SS (Type III)** | **DF** | **MS** | **F (DFn, DFd)** | **P value** |
| --- | --- | --- | --- | --- | --- | --- |
| Interaction (E:T × subset) | 20.28 | 43.44 | 4 | 10.86 | F(4, 19) = 23.94 | < 0.0001 |
| E:T ratio | 34.88 | 74.72 | 4 | 18.68 | F(4, 19) = 41.18 | < 0.0001 |
| NK cell subset | 53.79 | 115.2 | 1 | 115.2 | F(1, 19) = 254.1 | < 0.0001 |
| Residual | — | 8.618 | 19 | 0.4536 | — | — |
| Total | — | 214.2 | 28 | — | — | — |

*Two-way ANOVA was performed in GraphPad Prism v10.6.1 with E:T ratio and NK cell subset as factors. n = 29 values across the design (one technical replicate excluded at the 1:4 ratio for CD16^dim^).*

**Table 2. Šidák's multiple comparisons test (within each E:T ratio, compare subsets).**

| **E:T ratio** | **Comparison** | **Mean diff. (LS)** | **95% CI of diff.** | **Adjusted P value** | **Summary** |
| --- | --- | --- | --- | --- | --- |
| 1:4 | CD16^bright^ vs. CD16^dim^ | −8.434 | −10.19 to −6.68 | < 0.0001 | **** |
| 1:2 | CD16^bright^ vs. CD16^dim^ | −5.263 | −6.83 to −3.70 | < 0.0001 | **** |
| 1:1 | CD16^bright^ vs. CD16^dim^ | −2.995 | −4.56 to −1.43 | 0.0001 | *** |
| 3:1 | CD16^bright^ vs. CD16^dim^ | −1.952 | −3.52 to −0.384 | 0.0107 | * |
| 10:1 | CD16^bright^ vs. CD16^dim^ | −1.438 | −3.01 to 0.130 | 0.0824 | ns |

*Significance summary: ns, not significant; * p < 0.05, ** p < 0.01, *** p < 0.001, **** p < 0.0001.*
