## Supplemental Table 6 for "CD56^dim^CD16^dim^ NK cells are the dominant effector cells against HIV-infected primary T-cells"

**Supplemental Table 6 for the statistical analysis of Figure 3C**

*Killing frequency of CD56^dim^CD16^bright^ and CD56^dim^CD16^dim^ NK cells against purified autologous productively HIV-1_SHM-1_-infected primary T-cells across effector-to-target (E:T) cell ratios*

*Design: Two-way ANOVA (E:T ratio × CD56^dim^ NK cell subset). Subsets: CD56^dim^CD16^bright^, CD56^dim^CD16^dim^. E:T ratios: 1:4, 1:2, 1:1, 3:1, 10:1. Post-hoc: Šidák's multiple comparisons test. n=3 per group; 30 values total.*

**Two-way ANOVA summary**

| **Source of variation** | **% of variation** | **F (DFn, DFd)** | **P value** | **Significance** |
| --- | --- | --- | --- | --- |
| CD56dim subset | 22.20 | F (1, 20) = 79.47 | <0.0001 | **** |
| E:T ratio | 53.89 | F (4, 20) = 48.23 | <0.0001 | **** |
| E:T × subset interaction | 18.32 | F (4, 20) = 16.39 | <0.0001 | **** |

**Šidák's multiple comparisons**

| **Comparison** | **Mean diff.** | **95% CI of diff.** | **t** | **DF** | **Adj. P value** |
| --- | --- | --- | --- | --- | --- |
| *Between-subset comparisons at each E:T ratio* | | | | | |
| 1:4 — CD16^bright^ vs CD16^dim^ | -0.7215 | -0.8721 to -0.5708 | 9.990 | 20 | <0.0001 **** |
| 1:2 — CD16^bright^ vs CD16^dim^ | -0.4411 | -0.5918 to -0.2905 | 6.108 | 20 | <0.0001 **** |
| 1:1 — CD16^bright^ vs CD16^dim^ | -0.1928 | -0.3434 to -0.04213 | 2.669 | 20 | 0.0147 * |
| 3:1 — CD16^bright^ vs CD16^dim^ | -0.06046 | -0.2111 to 0.09019 | 0.8371 | 20 | 0.4124 ns |
| 10:1 — CD16^bright^ vs CD16^dim^ | -0.02375 | -0.1744 to 0.1269 | 0.3288 | 20 | 0.7457 ns |
| *Within-subset comparisons across E:T ratios — CD56^dim^CD16^bright^* | | | | | |
| CD16^bright^ — 1:4 vs 1:2 | 0.1302 | -0.09681 to 0.3572 | 1.803 | 20 | 0.5954 ns |
| CD16^bright^ — 1:4 vs 1:1 | 0.1809 | -0.04610 to 0.4079 | 2.505 | 20 | 0.1913 ns |
| CD16^bright^ — 1:4 vs 3:1 | 0.2463 | 0.01934 to 0.4734 | 3.411 | 20 | 0.0274 * |
| CD16^bright^ — 1:4 vs 10:1 | 0.2594 | 0.03242 to 0.4864 | 3.592 | 20 | 0.0181 * |
| CD16^bright^ — 1:2 vs 1:1 | 0.05071 | -0.1763 to 0.2777 | 0.7022 | 20 | 0.9988 ns |
| CD16^bright^ — 1:2 vs 3:1 | 0.1162 | -0.1109 to 0.3432 | 1.608 | 20 | 0.7322 ns |
| CD16^bright^ — 1:2 vs 10:1 | 0.1292 | -0.09777 to 0.3562 | 1.789 | 20 | 0.6050 ns |
| CD16^bright^ — 1:1 vs 3:1 | 0.06544 | -0.1616 to 0.2925 | 0.9061 | 20 | 0.9910 ns |
| CD16^bright^ — 1:1 vs 10:1 | 0.07853 | -0.1485 to 0.3055 | 1.087 | 20 | 0.9674 ns |
| CD16^bright^ — 3:1 vs 10:1 | 0.01309 | -0.2139 to 0.2401 | 0.1812 | 20 | >0.9999 ns |
| *Within-subset comparisons across E:T ratios — CD56^dim^CD16^dim^* | | | | | |
| CD16^dim^ — 1:4 vs 1:2 | 0.4105 | 0.1835 to 0.6375 | 5.684 | 20 | 0.0001 *** |
| CD16^dim^ — 1:4 vs 1:1 | 0.7096 | 0.4826 to 0.9366 | 9.825 | 20 | <0.0001 **** |
| CD16^dim^ — 1:4 vs 3:1 | 0.9073 | 0.6803 to 1.134 | 12.56 | 20 | <0.0001 **** |
| CD16^dim^ — 1:4 vs 10:1 | 0.9571 | 0.7301 to 1.184 | 13.25 | 20 | <0.0001 **** |
| CD16^dim^ — 1:2 vs 1:1 | 0.2991 | 0.07207 to 0.5261 | 4.141 | 20 | 0.0050 ** |
| CD16^dim^ — 1:2 vs 3:1 | 0.4968 | 0.2698 to 0.7239 | 6.880 | 20 | <0.0001 **** |
| CD16^dim^ — 1:2 vs 10:1 | 0.5466 | 0.3196 to 0.7736 | 7.569 | 20 | <0.0001 **** |
| CD16^dim^ — 1:1 vs 3:1 | 0.1978 | -0.02924 to 0.4248 | 2.738 | 20 | 0.1197 ns |
| CD16^dim^ — 1:1 vs 10:1 | 0.2476 | 0.02055 to 0.4746 | 3.428 | 20 | 0.0263 * |
| CD16^dim^ — 3:1 vs 10:1 | 0.04979 | -0.1772 to 0.2768 | 0.6895 | 20 | 0.9990 ns |

*Significance: ns, not significant; * P < 0.05; ** P < 0.01; *** P < 0.001; **** P < 0.0001.*
