## Supplemental Table 7 for "CD56^dim^CD16^dim^ NK cells are the dominant effector cells against HIV-infected primary T-cells"

**Supplemental Table 7 for the statistical analysis of Figure 4—supplement 1A**

*Time course of CD107a degranulation by CD56^dim^CD16^bright^ and CD56^dim^CD16^dim^ NK cells against purified autologous productively HIV-1_SHM-1_ infected primary T cells*

**Panel A — 4-hour time course**

Design: Two-way ANOVA (subset × time). Subset: CD56^dim^CD16^bright^ vs CD56^dim^CD16^dim^. Time points: 0, 30 min, 1 hr, 2 hr, and 4 hr. Post-hoc: Šidák's multiple comparisons test, between-subset comparisons at each time point. n = 3 per group; 30 values total.

**Two-way ANOVA summary**

| **Source of variation** | **% of variation** | **F (DFn, DFd)** | **P value** | **Significance** |
| --- | --- | --- | --- | --- |
| Subset (CD16^bright^ vs CD16^dim^) | 50.62 | F (1, 20) = 2664 | <0.0001 | **** |
| Time (0, 30 min, 1 hr, 2 hr, 4 hr) | 35.22 | F (4, 20) = 463.3 | <0.0001 | **** |
| Subset × time interaction | 13.78 | F (4, 20) = 181.3 | <0.0001 | **** |

**CD56^dim^CD16^bright^ vs CD56^dim^CD16^dim^ at each time point**

| **Time** | **Mean diff. (CD16^bright^ − CD16^dim^)** | **95% CI of diff.** | **t** | **DF** | **Adj. P value** |
| --- | --- | --- | --- | --- | --- |
| 0 | 0.00 | −2.80 to 2.80 | 0.00 | 20 | >0.9999 ns |
| 30 min | −25.17 | −27.97 to −22.36 | 25.45 | 20 | <0.0001 **** |
| 1 hr | −34.84 | −37.64 to −32.03 | 35.24 | 20 | <0.0001 **** |
| 2 hr | −28.14 | −30.94 to −25.33 | 28.46 | 20 | <0.0001 **** |
| 4 hr | −25.97 | −28.77 to −23.17 | 26.27 | 20 | <0.0001 **** |

**Panel B — 60-minute early kinetics**

Design: Two-way ANOVA (subset × time). Subset: CD56^dim^CD16^bright^ vs CD56^dim^CD16^dim^. Time points: 0, 15, 30, 45, 60 minutes. Post-hoc: Šidák's multiple comparisons test, between-subset comparisons at each time point. n = 3 per group; 30 values total.

**Two-way ANOVA summary**

| **Source of variation** | **% of variation** | **F (DFn, DFd)** | **P value** | **Significance** |
| --- | --- | --- | --- | --- |
| Subset (CD16^bright^ vs CD16^dim^) | 24.98 | F (1, 20) = 438.3 | <0.0001 | **** |
| Time (0, 15, 30, 45, 60 min) | 46.37 | F (4, 20) = 203.4 | <0.0001 | **** |
| Subset × time interaction | 27.52 | F (4, 20) = 120.7 | <0.0001 | **** |

**CD56^dim^CD16^bright^ vs CD56^dim^CD16^dim^ at each time point**

| **Time** | **Mean diff. (CD16^bright^ − CD16^dim^)** | **95% CI of diff.** | **t** | **DF** | **Adj. P value** |
| --- | --- | --- | --- | --- | --- |
| 0 min | 0.00 | −1.43 to 1.43 | 0.00 | 20 | >0.9999 ns |
| 15 min | −0.53 | −1.95 to 0.90 | 1.04 | 20 | 0.8423 ns |
| 30 min | −3.12 | −4.55 to −1.69 | 6.20 | 20 | <0.0001 **** |
| 45 min | −6.42 | −7.84 to −4.99 | 12.76 | 20 | <0.0001 **** |
| 60 min | −13.49 | −14.91 to −12.06 | 26.81 | 20 | <0.0001 **** |

*Significance: ns, not significant; **** P < 0.0001.*
