## Supplemental Table 9 for "CD56^dim^CD16^dim^ NK cells are the dominant effector cells against HIV-infected primary T-cells"

**Supplemental Table 9 for the statistical analysis of Figure 4**

*Statistical analysis of serial degranulation by CD56^dim^ CD16^bright^ and CD56^dim^ CD16^dim^ NK cells against purified autologous HIV-1_SHM-1_ productively infected T cells over a 60-minute period, stratified by number of degranulation events (0, 1, 2, or 3) per NK cell.*

**Table 1. Two-way ANOVA summary.**

| **Source of Variation** | **% of total variation** | **SS** | **DF** | **MS** | **F (DFn, DFd)** | **P value** |
| --- | --- | --- | --- | --- | --- | --- |
| Interaction (subset × events) | 30.96 | 3742 | 3 | 1247 | F(3, 16) = 1478 | < 0.0001 |
| Number of degranulation events | 68.84 | 8320 | 3 | 2773 | F(3, 16) = 3287 | < 0.0001 |
| NK cell subset | 0.089 | 10.80 | 1 | 10.80 | F(1, 16) = 12.80 | 0.0025 |
| Residual | — | 13.50 | 16 | 0.8436 | — | — |
| Total | — | 12086 | 23 | — | — | — |

*Two-way ANOVA was performed in GraphPad Prism v10.6.1 with the number of degranulation events and NK cell subset as factors. n = 24 values (2 subsets × 4 event categories × 3 replicates).*

**Table 2. Šidák's multiple comparisons test (within each event category, compare subsets).**

| **Number of events** | **Comparison** | **Mean diff.** | **95% CI of diff.** | **Adjusted P value** | **Summary** |
| --- | --- | --- | --- | --- | --- |
| 0 | CD16^bright^ vs. CD16^dim^ | 40.61 | 38.50 to 42.71 | < 0.0001 | **** |
| 1 | CD16^bright^ vs. CD16^dim^ | −9.784 | −11.89 to −7.68 | < 0.0001 | **** |
| 2 | CD16^bright^ vs. CD16^dim^ | −10.94 | −13.05 to −8.84 | < 0.0001 | **** |
| 3 | CD16^bright^ vs. CD16^dim^ | −25.25 | −27.35 to −23.14 | < 0.0001 | **** |

*Significance summary: * p < 0.05, ** p < 0.01, *** p < 0.001, **** p < 0.0001.*
