## Supplemental Table 10 for "CD56^dim^CD16^dim^ NK cells are the dominant effector cells against HIV-infected primary T-cells"

**Supplemental Table 10 for the statistical analysis of** **Figure 1—figure supplement 3A**

*IFN-γ expression by CD56^dim^CD16^bright^ and CD56^dim^CD16^dim^ NK cells against purified autologous productively HIV-1_SHM-1_ infected primary T-cells at various effector cell-to-target (E:T) cell ratios*

**Panel A — IFN-γ production**

Design: Two-way ANOVA (CD16 subset × E:T ratio). Subset: CD56^dim^CD16^bright^ vs CD56^dim^CD16^dim^. E:T ratios: 1:4, 1:2, 1:1, 2:1. Post-hoc: Šidák's multiple comparisons test. n = 3 per group; 24 values total.

**Two-way ANOVA summary**

| **Source of variation** | **% of variation** | **F (DFn, DFd)** | **P value** | **Significance** |
| --- | --- | --- | --- | --- |
| CD16 subset | 59.44 | F (1, 16) = 251.4 | <0.0001 | **** |
| E:T ratio | 21.33 | F (3, 16) = 30.08 | <0.0001 | **** |
| Subset × E:T interaction | 15.44 | F (3, 16) = 21.77 | <0.0001 | **** |

**Šidák's multiple comparisons**

| **Comparison** | **Mean diff.** | **95% CI of diff.** | **t** | **DF** | **Adj. P value** |
| --- | --- | --- | --- | --- | --- |
| ***Between-subset comparisons at each E:T ratio*** | | | | | |
| 1:4 — CD16^bright^ vs CD16^dim^ | −4.152 | −4.801 to −3.503 | 13.55 | 16 | <0.0001 **** |
| 1:2 — CD16^bright^ vs CD16^dim^ | −2.819 | −3.468 to −2.169 | 9.20 | 16 | <0.0001 **** |
| 1:1 — CD16^bright^ vs CD16^dim^ | −1.995 | −2.644 to −1.345 | 6.51 | 16 | <0.0001 **** |
| 2:1 — CD16^bright^ vs CD16^dim^ | −0.750 | −1.399 to −0.100 | 2.45 | 16 | 0.0263 * |
| ***Within-subset comparisons across E:T ratios*** | | | | | |
| CD16^bright^ — 1:4 vs 1:2 | 0.068 | −0.851 to 0.986 | 0.22 | 16 | >0.9999 ns |
| CD16^bright^ — 1:4 vs 1:1 | 0.279 | −0.639 to 1.198 | 0.91 | 16 | 0.9406 ns |
| CD16^bright^ — 1:4 vs 2:1 | 0.257 | −0.661 to 1.176 | 0.84 | 16 | 0.9593 ns |
| CD16^bright^ — 1:2 vs 1:1 | 0.212 | −0.707 to 1.130 | 0.69 | 16 | 0.9842 ns |
| CD16^bright^ — 1:2 vs 2:1 | 0.190 | −0.729 to 1.108 | 0.62 | 16 | 0.9911 ns |
| CD16^bright^ — 1:1 vs 2:1 | −0.022 | −0.941 to 0.896 | 0.07 | 16 | >0.9999 ns |
| CD16^dim^ — 1:4 vs 1:2 | 1.401 | 0.483 to 2.319 | 4.57 | 16 | 0.0019 ** |
| CD16^dim^ — 1:4 vs 1:1 | 2.437 | 1.518 to 3.355 | 7.95 | 16 | <0.0001 **** |
| CD16^dim^ — 1:4 vs 2:1 | 3.660 | 2.741 to 4.578 | 11.95 | 16 | <0.0001 **** |
| CD16^dim^ — 1:2 vs 1:1 | 1.036 | 0.117 to 1.954 | 3.38 | 16 | 0.0226 * |
| CD16^dim^ — 1:2 vs 2:1 | 2.259 | 1.340 to 3.177 | 7.37 | 16 | <0.0001 **** |
| CD16^dim^ — 1:1 vs 2:1 | 1.223 | 0.305 to 2.141 | 3.99 | 16 | 0.0063 ** |
