## Supplemental Table 11 for "CD56^dim^CD16^dim^ NK cells are the dominant effector cells against HIV-infected primary T-cells"

**Supplemental Table 11 for the statistical analysis of Figure 1—figure supplemental 1B**

*Degranulation of CD56^dim^CD16^bright^ and CD56^dim^CD16^dim^ NK cells (from the CD56^dim^ subsets used in the experiment shown in sFig 4A) against purified autologous productively HIV-1_SHM-1_ infected primary T-cells at various effector cell-to-target (E:T) cell ratios.*

Design: Two-way ANOVA (CD16 subset × E:T ratio). Subset: CD56^dim^CD16^bright^ vs CD56^dim^CD16^dim^. E:T ratios: 1:4, 1:2, 1:1, 2:1. Post-hoc: Šidák's multiple comparisons test. n = 3 per group; 24 values total.

**Two-way ANOVA summary**

| **Source of variation** | **% of variation** | **F (DFn, DFd)** | **P value** | **Significance** |
| --- | --- | --- | --- | --- |
| CD16 subset | 56.01 | F (1, 16) = 649.1 | <0.0001 | **** |
| E:T ratio | 29.54 | F (3, 16) = 114.1 | <0.0001 | **** |
| Subset × E:T interaction | 13.07 | F (3, 16) = 50.49 | <0.0001 | **** |

**Šidák's multiple comparisons (between-subset at each E:T ratio)**

| **Comparison** | **Mean diff.** | **95% CI of diff.** | **t** | **DF** | **Adj. P value** |
| --- | --- | --- | --- | --- | --- |
| ***Between-subset comparisons at each E:T ratio*** | | | | | |
| 1:4 — CD16^bright^ vs CD16^dim^ | −12.13 | −13.81 to −10.45 | 20.26 | 16 | <0.0001 **** |
| 1:2 — CD16^bright^ vs CD16^dim^ | −9.511 | −11.19 to −7.832 | 15.88 | 16 | <0.0001 **** |
| 1:1 — CD16^bright^ vs CD16^dim^ | −6.687 | −8.367 to −5.008 | 11.16 | 16 | <0.0001 **** |
| 2:1 — CD16^bright^ vs CD16^dim^ | −2.187 | −3.866 to −0.508 | 3.65 | 16 | 0.0086 ** |
