## Supplemental Table 12 for "CD56^dim^CD16^dim^ NK cells are the dominant effector cells against HIV-infected primary T-cells"

**Supplemental Table 12 for the statistical analysis of Figure 1—figure supplement 3C**

*Statistical analysis of CD56^dim^CD16^bright^ and CD56^dim^CD16^dim^ NK cell degranulation against autologous uninfected and purified HIV-1_SHM-1_, productively infected T-cells across five effector-to-target (E:T) cell ratios (0.25:1, 0.5:1, 1:1, 2:1, and 4:1).*

**Table 1. Two-way ANOVA summary.**

| **Source of Variation** | **% of total variation** | **SS** | **DF** | **MS** | **F (DFn, DFd)** | **P value** |
| --- | --- | --- | --- | --- | --- | --- |
| Interaction (E:T × condition) | 13.37 | 246.4 | 12 | 20.53 | F(12, 40) = 18.94 | < 0.0001 |
| E:T ratio | 20.16 | 371.6 | 4 | 92.90 | F(4, 40) = 85.69 | < 0.0001 |
| Condition (subset × target type) | 64.11 | 1182 | 3 | 393.9 | F(3, 40) = 363.3 | < 0.0001 |
| Residual | — | 43.37 | 40 | 1.084 | — | — |
| Total | — | 1843 | 59 | — | — | — |

*Two-way ANOVA was performed in GraphPad Prism v10.6.1 with E:T ratio and condition (NK cell subset × target type) as factors. n = 60 values (4 conditions × 5 E:T ratios × 3 replicates per condition per ratio).*

**Table 2. Šidák's multiple comparisons test (main column effect).**

| **Comparison** | **Mean diff.** | **95% CI of diff.** | **Adjusted P value** | **Summary** | **Significant?** |
| --- | --- | --- | --- | --- | --- |
| CD16^bright^ uninfected vs. CD16^dim^ uninfected | −2.848 | −3.901 to −1.796 | < 0.0001 | **** | Yes |
| CD16^bright^ uninfected vs. CD16^bright^ HIV-infected | −1.793 | −2.845 to −0.7408 | 0.0002 | *** | Yes |
| CD16^bright^ uninfected vs. CD16^dim^ HIV-infected | −11.52 | −12.57 to −10.47 | < 0.0001 | **** | Yes |
| CD16^dim^ uninfected vs. CD16^bright^ HIV-infected | 1.055 | 0.003 to 2.108 | 0.0490 | * | Yes |
| CD16^dim^ uninfected vs. CD16^dim^ HIV-infected | −8.674 | −9.726 to −7.621 | < 0.0001 | **** | Yes |
| CD16^bright^ HIV-infected vs. CD16^dim^ HIV-infected | −9.729 | −10.78 to −8.677 | < 0.0001 | **** | Yes |

*All six pairwise comparisons among the four conditions were significant after correction for multiple comparisons. Significance summary: * p < 0.05, ** p < 0.01, *** p < 0.001, **** p < 0.0001.*
