## Supplemental Table 13 for "CD56^dim^CD16^dim^ NK cells are the dominant effector cells against HIV-infected primary T-cells"

**Supplemental Table 13 for the statistical analysis of Figure 5—figure supplement 2**

*KIR3DL1 frequency and density on CD56^dim^ NK cell subsets and KIR3DL1-stratified degranulation responses to purified autologous productively HIV-1_SHM-1_-infected primary T cells*

**Panel 1 — KIR3DL1 frequency**

Design: Two-way ANOVA (CD16 subset × E:T ratio). Subsets: CD56^dim^CD16^bright^, CD56^dim^CD16^dim^, and CD56^dim^CD16^negative^. E:T ratios: no targets, 3:1, 1:1. Post-hoc: Šidák's multiple comparisons test. n = 3 per group; 27 values total.

**Two-way ANOVA summary**

| **Source of variation** | **% of variation** | **F (DFn, DFd)** | **P value** | **Significance** |
| --- | --- | --- | --- | --- |
| CD16 subset | 77.50 | F (2, 18) = 54.34 | <0.0001 | **** |
| E:T ratio | 2.83 | F (2, 18) = 1.99 | 0.1661 | ns |
| Subset × E:T interaction | 6.83 | F (4, 18) = 2.39 | 0.0888 | ns |

**Šidák's multiple comparisons**

| **Comparison** | **Mean diff.** | **95% CI of diff.** | **stat** | **DF** | **Adj. P value** |
| --- | --- | --- | --- | --- | --- |
| ***Between-subset comparisons at each E:T ratio*** | | | | | |
| No targets CD16^bright^ vs CD16^dim^ | 2.400 | 0.996 to 3.804 | t=4.50 | 18 | 0.0008 *** |
| No targets CD16^brigh^t vs CD16^neg^ | −1.533 | −2.938 to −0.129 | t=2.87 | 18 | 0.0301 * |
| No targets — CD16^dim^ vs CD16^neg^ | −3.933 | −5.338 to −2.529 | t=7.37 | 18 | <0.0001 **** |
| 3:1 — CD16^bright^ vs CD16^dim^ | 0.467 | −0.938 to 1.871 | t=0.87 | 18 | 0.7769 ns |
| 3:1 — CD16^bright^ vs CD16^neg^ | −1.700 | −3.104 to −0.296 | t=3.19 | 18 | 0.0153 * |
| 3:1 — CD16^dim^ vs CD16^negative^ | −2.167 | −3.571 to −0.762 | t=4.06 | 18 | 0.0022 ** |
| 1:1 — CD16^brigh^t vs CD16^dim^ | 2.233 | 0.829 to 3.638 | t=4.18 | 18 | 0.0017 ** |
| 1:1 — CD16^bright^ vs CD16^negative^ | −1.300 | −2.704 to 0.104 | t=2.43 | 18 | 0.0746 ns |
| 1:1 — CD16^dim^ vs CD16^negative^ | −3.533 | −4.938 to −2.129 | t=6.62 | 18 | <0.0001 **** |
| ***Within-subset comparisons across E:T ratios*** | | | | | |
| CD16^bright^ — No targets vs 3:1 | 0.133 | −1.271 to 1.538 | t=0.25 | 18 | 0.9927 ns |
| CD16^bright^ — No targets vs 1:1 | −0.100 | −1.504 to 1.304 | t=0.19 | 18 | 0.9969 ns |
| CD16^bright^ — 3:1 vs 1:1 | −0.233 | −1.638 to 1.171 | t=0.44 | 18 | 0.9631 ns |
| CD16^dim^ — No targets vs 3:1 | −1.800 | −3.204 to −0.396 | t=3.37 | 18 | 0.0102 * |
| CD16^dim^ — No targets vs 1:1 | −0.267 | −1.671 to 1.138 | t=0.50 | 18 | 0.9466 ns |
| CD16^dim^ — 3:1 vs 1:1 | 1.533 | 0.129 to 2.938 | t=2.87 | 18 | 0.0301 * |
| CD16^negative^ — No targets vs 3:1 | −0.033 | −1.438 to 1.371 | t=0.06 | 18 | 0.9999 ns |
| CD16^negative^ — No targets vs 1:1 | 0.133 | −1.271 to 1.538 | t=0.25 | 18 | 0.9927 ns |
| CD16^negative^ — 3:1 vs 1:1 | 0.167 | −1.238 to 1.571 | t=0.31 | 18 | 0.9859 ns |

**Panel 2 — KIR3DL1 density (gMFI)**

Design: Two-way ANOVA (CD16 subset × E:T ratio). Subsets: CD56dimCD16bright, CD56dimCD16dim, CD56dimCD16neg. E:T ratios: no targets, 3:1, 1:1. Post-hoc: Tukey's multiple comparisons test. n = 3 per group; 27 values total.

**Two-way ANOVA summary**

| **Source of variation** | **% of variation** | **F (DFn, DFd)** | **P value** | **Significance** |
| --- | --- | --- | --- | --- |
| CD16 subset | 98.68 | F (2, 18) = 3970 | <0.0001 | **** |
| E:T ratio | 0.27 | F (2, 18) = 10.70 | 0.0009 | *** |
| Subset × E:T interaction | 0.83 | F (4, 18) = 16.78 | <0.0001 | **** |

**Tukey's multiple comparisons**

| **Comparison** | **Mean diff.** | **95% CI of diff.** | **stat** | **DF** | **Adj. P value** |
| --- | --- | --- | --- | --- | --- |
| ***Between-subset comparisons at each E:T ratio*** | | | | | |
| No targets — CD16^bright^ vs CD16^dim^ | 5025 | 4625 to 5425 | q=45.4 | 18 | <0.0001 **** |
| No targets — CD16^bright^ vs CD16^negative^ | 8146 | 7747 to 8546 | q=73.6 | 18 | <0.0001 **** |
| No targets — CD16^dim^ vs CD16^negative^ | 3121 | 2722 to 3521 | q=28.2 | 18 | <0.0001 **** |
| 3:1 — CD16^bright^ vs CD16^dim^ | 3666 | 3266 to 4066 | q=33.1 | 18 | <0.0001 **** |
| 3:1 — CD16^bright^ vs CD16^negative^ | 8004 | 7604 to 8404 | q=72.3 | 18 | <0.0001 **** |
| 3:1 — CD16^dim^ vs CD16^negative^ | 4338 | 3938 to 4738 | q=39.2 | 18 | <0.0001 **** |
| 1:1 — CD16^bright^ vs CD16^dim^ | 3547 | 3147 to 3946 | q=32.0 | 18 | <0.0001 **** |
| 1:1 — CD16^bright^ vs CD16^negative^ | 8018 | 7619 to 8418 | q=72.4 | 18 | <0.0001 **** |
| 1:1 — CD16^dim^ vs CD16^negative^ | 4472 | 4072 to 4871 | q=40.4 | 18 | <0.0001 **** |
| ***Within-subset comparisons across E:T ratios*** | | | | | |
| CD16^bright^ — No targets vs 3:1 | 157.3 | −242.3 to 557.0 | q=1.42 | 18 | 0.5834 ns |
| CD16^bright^ — No targets vs 1:1 | 156.7 | −243.0 to 556.3 | q=1.42 | 18 | 0.5860 ns |
| CD16^bright^ — 3:1 vs 1:1 | −0.667 | −400.3 to 399.0 | q=0.01 | 18 | >0.9999 ns |
| CD16^dim^ — No targets vs 3:1 | −1202 | −1601 to −802.0 | q=10.85 | 18 | <0.0001 **** |
| CD16^dim^ — No targets vs 1:1 | −1322 | −1721 to −922.0 | q=11.94 | 18 | <0.0001 **** |
| CD16^dim^ — 3:1 vs 1:1 | −120.0 | −519.7 to 279.7 | q=1.08 | 18 | 0.7279 ns |
| CD16^negative^ — No targets vs 3:1 | 15.00 | −384.7 to 414.7 | q=0.14 | 18 | 0.9950 ns |
| CD16^negative^ — No targets vs 1:1 | 28.67 | −371.0 to 428.3 | q=0.26 | 18 | 0.9817 ns |
| CD16^negative^ — 3:1 vs 1:1 | 13.67 | −386.0 to 413.3 | q=0.12 | 18 | 0.9958 ns |

**Panel 3 — Degranulation stratified by KIR3DL1 expression**

Design: Two-way ANOVA (CD16 subset × KIR3DL1 expression). Subsets: CD56^dim^CD16^bright^, CD56^dim^CD16^dim^, CD56^dim^CD16^negative^. KIR3DL1 status: KIR3DL1^positive^, KIR3DL1^negative^. Post-hoc: Šidák's multiple comparisons test. n = 3 per group; 18 values total.

**Two-way ANOVA summary**

| **Source of variation** | **% of variation** | **F (DFn, DFd)** | **P value** | **Significance** |
| --- | --- | --- | --- | --- |
| CD16 subset | 78.25 | F (2, 12) = 80.34 | <0.0001 | **** |
| KIR3DL1 expression (+ vs −) | 6.28 | F (1, 12) = 12.90 | 0.0037 | ** |
| Subset × KIR3DL1 interaction | 9.62 | F (2, 12) = 9.88 | 0.0029 | ** |

**Šidák's multiple comparisons**

| **Comparison** | **Mean diff.** | **95% CI of diff.** | **stat** | **DF** | **Adj. P value** |
| --- | --- | --- | --- | --- | --- |
| ***Between-subset comparisons within each KIR3DL1 status*** | | | | | |
| KIR3DL1+ — CD16^bright^ vs CD16^dim^ | −6.783 | −8.646 to −4.920 | t=10.09 | 12 | <0.0001 **** |
| KIR3DL1+ — CD16^bright^ vs CD16^negative^ | 0.426 | −1.437 to 2.288 | t=0.63 | 12 | 0.9019 ns |
| KIR3DL1+ — CD16^dim^ vs CD16^negative^ | 7.208 | 5.346 to 9.071 | t=10.72 | 12 | <0.0001 **** |
| KIR3DL1− — CD16^bright^ vs CD16^dim^ | −2.870 | −4.732 to −1.007 | t=4.27 | 12 | 0.0033 ** |
| KIR3DL1− — CD16^bright^ vs CD16^negative^ | 1.000 | −0.863 to 2.862 | t=1.49 | 12 | 0.4135 ns |
| KIR3DL1− — CD16^dim^ vs CD16^negative^ | 3.869 | 2.006 to 5.732 | t=5.75 | 12 | 0.0003 *** |
| ***Within-subset comparisons: KIR3DL1+ vs KIR3DL1−*** | | | | | |
| CD16^bright^ — KIR3DL1+ vs KIR3DL1− | −0.102 | −1.567 to 1.363 | t=0.15 | 12 | 0.8822 ns |
| CD16^dim^ — KIR3DL1+ vs KIR3DL1− | 3.812 | 2.347 to 5.277 | t=5.67 | 12 | 0.0001 *** |
| CD16^negative^ — KIR3DL1+ vs KIR3DL1− | 0.472 | −0.993 to 1.937 | t=0.70 | 12 | 0.4958 ns |

*Significance: ns, not significant; * P < 0.05; ** P < 0.01; *** P < 0.001; **** P < 0.0001.*
