## Supplemental Table 14 for "CD56^dim^CD16^dim^ NK cells are the dominant effector cells against HIV-infected primary T-cells"

**Supplemental Table 14 for the statistical analysis of Figure 5—figure supplement 3**

*KIR2DL2/3 frequency and density on CD56^dim^ NK cell subsets and KIR2DL2/3-stratified degranulation responses to purified autologous productively HIV-1_SHM-1_ infected primary T-cells at an effector cell-to-target cell (E:T) ratio of 3:1 and 1:1. As a control, we also looked at CD56^dim^ NK cell subsets not exposed to target cells.*

**Panel 1 — KIR2DL2/3 frequency**

Design: Two-way ANOVA (CD16 subset × E:T ratio). Subsets: CD56^dim^CD16^bright^, CD56^dim^CD16^dim^, CD56^dim^CD16^negative^. E:T ratios: no targets, 3:1, 1:1. Post-hoc: Šidák's multiple comparisons test. n = 3 per group; 27 values total.

**Two-way ANOVA summary**

| **Source of variation** | **% of variation** | **F (DFn, DFd)** | **P value** | **Significance** |
| --- | --- | --- | --- | --- |
| CD16 subset | 81.90 | F (2, 18) = 54.61 | <0.0001 | **** |
| E:T ratio | 3.27 | F (2, 18) = 2.18 | 0.1417 | ns |
| Subset × E:T interaction | 1.33 | F (4, 18) = 0.44 | 0.7757 | ns |

**Šidák's multiple comparisons**

| **Comparison** | **Mean diff.** | **95% CI of diff.** | **t** | **DF** | **Adj. P value** |
| --- | --- | --- | --- | --- | --- |
| ***Between-subset comparisons at each E:T ratio*** | | | | | |
| No targets — CD16^bright^ vs CD16^dim^ | 7.500 | 4.037 to 10.96 | 5.70 | 18 | <0.0001 **** |
| No targets — CD16^bright^ vs CD16^negative^ | 8.133 | 4.670 to 11.60 | 6.18 | 18 | <0.0001 **** |
| No targets — CD16^dim^ vs CD16^negative^ | 0.633 | −2.830 to 4.097 | 0.48 | 18 | 0.9519 ns |
| 3:1 — CD16^bright^ vs CD16^dim^ | 6.700 | 3.237 to 10.16 | 5.09 | 18 | 0.0002 *** |
| 3:1 — CD16^bright^ vs CD16^neg^ | 7.400 | 3.937 to 10.86 | 5.62 | 18 | <0.0001 **** |
| 3:1 — CD16^dim^ vs CD16^negative^ | 0.700 | −2.763 to 4.163 | 0.53 | 18 | 0.9367 ns |
| 1:1 — CD16^bright^ vs CD16^dim^ | 5.533 | 2.070 to 8.997 | 4.20 | 18 | 0.0016 ** |
| 1:1 — CD16^bright^ vs CD16^negative^ | 5.900 | 2.437 to 9.363 | 4.48 | 18 | 0.0009 *** |
| 1:1 — CD16^dim^ vs CD16^negative^ | 0.367 | −3.097 to 3.830 | 0.28 | 18 | 0.9899 ns |
| ***Within-subset comparisons across E:T ratios*** | | | | | |
| CD16^bright^ — No targets vs 3:1 | −1.000 | −4.463 to 2.463 | 0.76 | 18 | 0.8402 ns |
| CD16^bright^ — No targets vs 1:1 | 1.067 | −2.397 to 4.530 | 0.81 | 18 | 0.8132 ns |
| CD16^bright^ — 3:1 vs 1:1 | 2.067 | −1.397 to 5.530 | 1.57 | 18 | 0.3502 ns |
| CD16^dim^ — No targets vs 3:1 | −1.800 | −5.263 to 1.663 | 1.37 | 18 | 0.4653 ns |
| CD16^dim^ — No targets vs 1:1 | −0.900 | −4.363 to 2.563 | 0.68 | 18 | 0.8771 ns |
| CD16^dim^ — 3:1 vs 1:1 | 0.900 | −2.563 to 4.363 | 0.68 | 18 | 0.8771 ns |
| CD16^negative^ — No targets vs 3:1 | −1.733 | −5.197 to 1.730 | 1.32 | 18 | 0.4965 ns |
| CD16^negative^ — No targets vs 1:1 | −1.167 | −4.630 to 2.297 | 0.89 | 18 | 0.7698 ns |
| CD16^negative^ — 3:1 vs 1:1 | 0.567 | −2.897 to 4.030 | 0.43 | 18 | 0.9647 ns |

**Panel 2 — KIR2DL2/3 density (gMFI)**

Design: Two-way ANOVA (CD16 subset × E:T ratio). Subsets: CD56^dim^CD16^bright^, CD56^dim^CD16^dim^, CD56^dim^CD16^negative^. E:T ratios: no targets, 3:1, 1:1. Post-hoc: Šidák's multiple comparisons test. n = 3 per group; 27 values total.

**Two-way ANOVA summary**

| **Source of variation** | **% of variation** | **F (DFn, DFd)** | **P value** | **Significance** |
| --- | --- | --- | --- | --- |
| CD16 subset | 84.76 | F (2, 18) = 58.68 | <0.0001 | **** |
| E:T ratio | 0.93 | F (2, 18) = 0.65 | 0.5363 | ns |
| Subset × E:T interaction | 1.31 | F (4, 18) = 0.45 | 0.7685 | ns |

**Šidák's multiple comparisons**

| **Comparison** | **Mean diff.** | **95% CI of diff.** | **t** | **DF** | **Adj. P value** |
| --- | --- | --- | --- | --- | --- |
| ***Between-subset comparisons at each E:T ratio*** | | | | | |
| No targets — CD16^bright^ vs CD16^dim^ | 402.0 | 137.5 to 666.5 | 4.00 | 18 | 0.0025 ** |
| No targets — CD16^bright^ vs CD16^negative^ | 714.0 | 449.5 to 978.5 | 7.10 | 18 | <0.0001 **** |
| No targets — CD16^dim^ vs CD16^negative^ | 312.0 | 47.53 to 576.5 | 3.10 | 18 | 0.0183 * |
| 3:1 — CD16^bright^ vs CD16^dim^ | 241.0 | −23.47 to 505.5 | 2.40 | 18 | 0.0804 ns |
| 3:1 — CD16^bright^ vs CD16^negative^ | 560.7 | 296.2 to 825.1 | 5.58 | 18 | <0.0001 **** |
| 3:1 — CD16^dim^ vs CD16^negative^ | 319.7 | 55.20 to 584.1 | 3.18 | 18 | 0.0155 * |
| 1:1 — CD16^bright^ vs CD16^dim^ | 269.0 | 4.533 to 533.5 | 2.68 | 18 | 0.0455 * |
| 1:1 — CD16^bright^ vs CD16^negative^ | 611.0 | 346.5 to 875.5 | 6.08 | 18 | <0.0001 **** |
| 1:1 — CD16^dim^ vs CD16^negative^ | 342.0 | 77.53 to 606.5 | 3.40 | 18 | 0.0095 ** |
| ***Within-subset comparisons across E:T ratios*** | | | | | |
| CD16^bright^ — No targets vs 3:1 | 169.0 | −95.47 to 433.5 | 1.68 | 18 | 0.2950 ns |
| CD16^bright^ — No targets vs 1:1 | 123.7 | −140.8 to 388.1 | 1.23 | 18 | 0.5513 ns |
| CD16^bright^ — 3:1 vs 1:1 | −45.33 | −309.8 to 219.1 | 0.45 | 18 | 0.9598 ns |
| CD16^dim^ — No targets vs 3:1 | 8.00 | −256.5 to 272.5 | 0.08 | 18 | 0.9998 ns |
| CD16^dim^ — No targets vs 1:1 | −9.33 | −273.8 to 255.1 | 0.09 | 18 | 0.9996 ns |
| CD16^dim^ — 3:1 vs 1:1 | −17.33 | −281.8 to 247.1 | 0.17 | 18 | 0.9975 ns |
| CD16^negative^ — No targets vs 3:1 | 15.67 | −248.8 to 280.1 | 0.16 | 18 | 0.9982 ns |
| CD16^negative^ — No targets vs 1:1 | 20.67 | −243.8 to 285.1 | 0.21 | 18 | 0.9959 ns |
| CD16^negative^ — 3:1 vs 1:1 | 5.00 | −259.5 to 269.5 | 0.05 | 18 | >0.9999 ns |

**Panel 3 — Degranulation stratified by KIR2DL2/3 expression**

Design: Two-way ANOVA (CD16 subset × KIR2DL2/3 expression). Subsets: CD56^dim^CD16^bright^, CD56^dim^CD16^dim^, CD56^dim^CD16^negative^. KIR2DL2/3 status: KIR2DL2/3+, KIR2DL2/3−. Post-hoc: Šidák's multiple comparisons test. n = 3 per group; 18 values total.

**Two-way ANOVA summary**

| **Source of variation** | **% of variation** | **F (DFn, DFd)** | **P value** | **Significance** |
| --- | --- | --- | --- | --- |
| CD16 subset | 83.31 | F (2, 12) = 67.44 | <0.0001 | **** |
| KIR2DL2/3 expression (+ vs −) | 4.29 | F (1, 12) = 6.95 | 0.0217 | * |
| Subset × KIR2DL2/3 interaction | 4.98 | F (2, 12) = 4.03 | 0.0457 | * |

**Šidák's multiple comparisons**

| **Comparison** | **Mean diff.** | **95% CI of diff.** | **t** | **DF** | **Adj. P value** |
| --- | --- | --- | --- | --- | --- |
| ***Between-subset comparisons within each KIR2DL2/3 status*** | | | | | |
| KIR2DL2/3+ — CD16^bright^ vs CD16^dim^ | −7.516 | −10.06 to −4.973 | 8.19 | 12 | <0.0001 **** |
| KIR2DL2/3+ — CD16^bright^ vs CD16^negative^ | 1.087 | −1.455 to 3.630 | 1.19 | 12 | 0.5933 ns |
| KIR2DL2/3+ — CD16^dim^ vs CD16^negative^ | 8.603 | 6.061 to 11.15 | 9.37 | 12 | <0.0001 **** |
| KIR2DL2/3− — CD16^bright^ vs CD16^dim^ | −4.359 | −6.901 to −1.816 | 4.75 | 12 | 0.0014 ** |
| KIR2DL2/3− — CD16^bright^ vs CD16^negative^ | 1.018 | −1.525 to 3.560 | 1.11 | 12 | 0.6410 ns |
| KIR2DL2/3− — CD16^dim^ vs CD16^negative^ | 5.376 | 2.834 to 7.919 | 5.86 | 12 | 0.0002 *** |
| ***Within-subset comparisons: KIR2DL2/3+ vs KIR2DL2/3−*** | | | | | |
| CD16^bright^ — KIR2DL2/3+ vs KIR2DL2/3− | 0.368 | −1.632 to 2.367 | 0.40 | 12 | 0.6958 ns |
| CD16^dim^ — KIR2DL2/3+ vs KIR2DL2/3− | 3.525 | 1.525 to 5.525 | 3.84 | 12 | 0.0023 ** |
| CD16^negative^ — KIR2DL2/3+ vs KIR2DL2/3− | 0.298 | −1.702 to 2.297 | 0.32 | 12 | 0.7512 ns |

*Significance: ns, not significant; * P < 0.05; ** P < 0.01; *** P < 0.001; **** P < 0.0001.*
