## Supplemental Table 15 for "CD56^dim^CD16^dim^ NK cells are the dominant effector cells against HIV-infected primary T-cells"

**Supplemental Table 15 for the statistical analysis of Figure 5—figure supplement 4**

*KIR2DL1 frequency and density on CD56^dim^ NK cell subsets and KIR2DL1-stratified degranulation responses to purified autologous productively HIV-1_SHM-1_-infected primary T-cells at an effector cell-to-target cell (E:T) ratio of 3:1 and 1:1. As a control, we also looked at CD56^dim^ NK cell subsets not exposed to target cells.*

**Panel 1 — KIR2DL1 frequency**

Design: Two-way ANOVA (CD16 subset × E:T ratio). Subsets: CD56^dim^CD16^bright^, CD56^dim^CD16^dim^, CD56^dim^CD16^negative^. E:T ratios: no targets, 3:1, 1:1. Post-hoc: Šidák's multiple comparisons test. n = 3 per group; 27 values total.

**Two-way ANOVA summary**

| **Source of variation** | **% of variation** | **F (DFn, DFd)** | **P value** | **Significance** |
| --- | --- | --- | --- | --- |
| CD16 subset | 97.02 | F (2, 18) = 376.8 | <0.0001 | **** |
| E:T ratio | 0.36 | F (2, 18) = 1.40 | 0.2732 | ns |
| Subset × E:T interaction | 0.30 | F (4, 18) = 0.59 | 0.6753 | ns |

**Šidák's multiple comparisons**

| **Comparison** | **Mean diff.** | **95% CI of diff.** | **t** | **DF** | **Adj. P value** |
| --- | --- | --- | --- | --- | --- |
| ***Between-subset comparisons at each E:T ratio*** | | | | | |
| No targets — CD16^bright^ vs CD16^dim^ | 2.993 | 2.295 to 3.691 | 11.29 | 18 | <0.0001 **** |
| No targets — CD16^bright^ vs CD16^negative^ | 4.090 | 3.392 to 4.788 | 15.42 | 18 | <0.0001 **** |
| No targets — CD16^dim^ vs CD16^negative^ | 1.097 | 0.399 to 1.795 | 4.14 | 18 | 0.0019 ** |
| 3:1 — CD16^bright^ vs CD16^dim^ | 3.123 | 2.425 to 3.821 | 11.78 | 18 | <0.0001 **** |
| 3:1 — CD16^bright^ vs CD16^negative^ | 4.153 | 3.455 to 4.851 | 15.66 | 18 | <0.0001 **** |
| 3:1 — CD16^dim^ vs CD16n^negative^ | 1.030 | 0.332 to 1.728 | 3.88 | 18 | 0.0033 ** |
| 1:1 — CD16^bright^ vs CD16dim | 2.593 | 1.895 to 3.291 | 9.78 | 18 | <0.0001 **** |
| 1:1 — CD16^bright^ vs CD16^negative^ | 4.010 | 3.312 to 4.708 | 15.12 | 18 | <0.0001 **** |
| 1:1 — CD16^dim^ vs CD16^negative^ | 1.417 | 0.719 to 2.115 | 5.34 | 18 | 0.0001 *** |
| ***Within-subset comparisons across E:T ratios*** | | | | | |
| CD16bright — No targets vs 3:1 | 0.093 | −0.605 to 0.791 | 0.35 | 18 | 0.9801 ns |
| CD16bright — No targets vs 1:1 | 0.413 | −0.285 to 1.111 | 1.56 | 18 | 0.3563 ns |
| CD16bright — 3:1 vs 1:1 | 0.320 | −0.378 to 1.018 | 1.21 | 18 | 0.5667 ns |
| CD16dim — No targets vs 3:1 | 0.223 | −0.475 to 0.921 | 0.84 | 18 | 0.7955 ns |
| CD16dim — No targets vs 1:1 | 0.013 | −0.685 to 0.711 | 0.05 | 18 | >0.9999 ns |
| CD16dim — 3:1 vs 1:1 | −0.210 | −0.908 to 0.488 | 0.79 | 18 | 0.8233 ns |
| CD16neg — No targets vs 3:1 | 0.157 | −0.541 to 0.855 | 0.59 | 18 | 0.9160 ns |
| CD16neg — No targets vs 1:1 | 0.333 | −0.365 to 1.031 | 1.26 | 18 | 0.5344 ns |
| CD16neg — 3:1 vs 1:1 | 0.177 | −0.521 to 0.875 | 0.67 | 18 | 0.8851 ns |

**Panel 2 — KIR2DL1 density (gMFI)**

Design: Two-way ANOVA (CD16 subset × E:T ratio). Subsets: CD56dimCD16bright, CD56dimCD16dim, CD56dimCD16neg. E:T ratios: no targets, 3:1, 1:1. Post-hoc: Šidák's multiple comparisons test. n = 3 per group; 27 values total.

**Two-way ANOVA summary**

| **Source of variation** | **% of variation** | **F (DFn, DFd)** | **P value** | **Significance** |
| --- | --- | --- | --- | --- |
| CD16 subset | 87.06 | F (2, 18) = 151.1 | <0.0001 | **** |
| E:T ratio | 3.35 | F (2, 18) = 5.82 | 0.0112 | * |
| Subset × E:T interaction | 4.40 | F (4, 18) = 3.82 | 0.0204 | * |

**Šidák's multiple comparisons**

| **Comparison** | **Mean diff.** | **95% CI of diff.** | **t** | **DF** | **Adj. P value** |
| --- | --- | --- | --- | --- | --- |
| ***Between-subset comparisons at each E:T ratio*** | | | | | |
| No targets — CD16^bright^ vs CD16^dim^ | 75.33 | −20.52 to 171.2 | 2.07 | 18 | 0.1517 ns |
| No targets — CD16^bright^ vs CD16^negative^ | 419.3 | 323.5 to 515.2 | 11.51 | 18 | <0.0001 **** |
| No targets — CD16^dim^ vs CD16^negative^ | 344.0 | 248.1 to 439.9 | 9.44 | 18 | <0.0001 **** |
| 3:1 — CD16^bright^ vs CD16^dim^ | −3.667 | −99.52 to 92.19 | 0.10 | 18 | 0.9995 ns |
| 3:1 — CD16^bright^ vs CD16^negative^ | 335.0 | 239.1 to 430.9 | 9.20 | 18 | <0.0001 **** |
| 3:1 — CD16^dim^ vs CD16^negative^ | 338.7 | 242.8 to 434.5 | 9.30 | 18 | <0.0001 **** |
| 1:1 — CD16^bright^ vs CD16^dim^ | −0.667 | −96.52 to 95.19 | 0.02 | 18 | >0.9999 ns |
| 1:1 — CD16^bright^ vs CD16^negative^ | 229.3 | 133.5 to 325.2 | 6.30 | 18 | <0.0001 **** |
| 1:1 — CD16^dim^ vs CD16^negative^ | 230.0 | 134.1 to 325.9 | 6.31 | 18 | <0.0001 **** |
| ***Within-subset comparisons across E:T ratios*** | | | | | |
| CD16^bright^ — No targets vs 3:1 | −14.67 | −110.5 to 81.19 | 0.40 | 18 | 0.9708 ns |
| CD16^bright^ — No targets vs 1:1 | 37.33 | −58.52 to 133.2 | 1.03 | 18 | 0.6843 ns |
| CD16^bright^ — 3:1 vs 1:1 | 52.00 | −43.86 to 147.9 | 1.43 | 18 | 0.4295 ns |
| CD16^dim^ — No targets vs 3:1 | −93.67 | −189.5 to 2.191 | 2.57 | 18 | 0.0566 ns |
| CD16^dim^ — No targets vs 1:1 | −38.67 | −134.5 to 57.19 | 1.06 | 18 | 0.6608 ns |
| CD16^dim^ — 3:1 vs 1:1 | 55.00 | −40.86 to 150.9 | 1.51 | 18 | 0.3826 ns |
| CD16^negative^ — No targets vs 3:1 | −99.00 | −194.9 to −3.143 | 2.72 | 18 | 0.0418 * |
| CD16^negative^ — No targets vs 1:1 | −152.7 | −248.5 to −56.81 | 4.19 | 18 | 0.0016 ** |
| CD16^negative^ — 3:1 vs 1:1 | −53.67 | −149.5 to 42.19 | 1.47 | 18 | 0.4031 ns |

**Panel 3 — Degranulation stratified by KIR2DL1 expression**

Design: Two-way ANOVA (CD16 subset × KIR2DL1 expression). Subsets: CD56dimCD16bright, CD56dimCD16dim, CD56dimCD16neg. KIR2DL1 status: KIR2DL1+, KIR2DL1−. Post-hoc: Šidák's multiple comparisons test.

**Šidák's multiple comparisons**

| **Comparison** | **Mean diff.** | **95% CI of diff.** | **t** | **DF** | **Adj. P value** |
| --- | --- | --- | --- | --- | --- |
| ***Between-subset comparisons within each KIR2DL1 status*** | | | | | |
| KIR2DL1+ — CD16^bright^ vs CD16^dim^ | −3.270 | −5.013 to −1.527 | 5.20 | 12 | 0.0007 *** |
| KIR2DL1+ — CD16^bright^ vs CD16^negative^ | −0.782 | −2.526 to 0.961 | 1.24 | 12 | 0.5567 ns |
| KIR2DL1+ — CD16^dim^ vs CD16^negative^ | 2.488 | 0.744 to 4.231 | 3.95 | 12 | 0.0057 ** |
| KIR2DL1− — CD16^bright^ vs CD16^dim^ | −3.920 | −5.664 to −2.177 | 6.23 | 12 | 0.0001 *** |
| KIR2DL1− — CD16^bright^ vs CD16^negative^ | 1.201 | −0.543 to 2.944 | 1.91 | 12 | 0.2229 ns |
| KIR2DL1− — CD16^dim^ vs CD16^negative^ | 5.121 | 3.378 to 6.864 | 8.14 | 12 | <0.0001 **** |
| ***Within-subset comparisons: KIR2DL1+ vs KIR2DL1−*** | | | | | |
| CD16^bright^ — KIR2DL1+ vs KIR2DL1− | −0.076 | −1.447 to 1.296 | 0.12 | 12 | 0.9063 ns |
| CD16^dim^ — KIR2DL1+ vs KIR2DL1− | −0.726 | −2.097 to 0.645 | 1.15 | 12 | 0.2711 ns |
| CD16^negative^ — KIR2DL1+ vs KIR2DL1− | 1.907 | 0.536 to 3.279 | 3.03 | 12 | 0.0105 * |

*Significance: ns, not significant; * P < 0.05; ** P < 0.01; *** P < 0.001; **** P < 0.0001.*
