## Supplemental Table 16 for "CD56^dim^CD16^dim^ NK cells are the dominant effector cells against HIV-infected primary T-cells"

**Supplemental Table 16 for the statistical analysis of Figure 5—figure supplement 5**

*NKG2A frequency and density on CD56dim NK cell subsets and NKG2A-stratified degranulation responses to purified autologous productively HIV-1_SHM-1_ infected primary T cells*

**Panel 1 — NKG2A frequency**

Design: Two-way ANOVA (CD16 subset × E:T ratio). Subsets: CD56dimCD16bright, CD56dimCD16dim, CD56dimCD16neg. E:T ratios: no targets, 3:1, 1:1. Post-hoc: Šidák's multiple comparisons test. n = 3 per group; 27 values total. NKG2A frequency is at ceiling (>99% across all subsets and conditions); the two flagged significant differences reflect 0.2–0.3% point differences that are not biologically meaningful.

**Šidák's multiple comparisons**

| **Comparison** | **Mean diff.** | **95% CI of diff.** | **t** | **DF** | **Adj. P value** |
| --- | --- | --- | --- | --- | --- |
| ***Between-subset comparisons at each E:T ratio*** | | | | | |
| No targets — CD16bright vs CD16dim | −0.167 | −0.482 to 0.148 | 1.39 | 18 | 0.4500 ns |
| No targets — CD16bright vs CD16neg | −0.367 | −0.682 to −0.052 | 3.06 | 18 | 0.0199 * |
| No targets — CD16dim vs CD16neg | −0.200 | −0.515 to 0.115 | 1.67 | 18 | 0.2997 ns |
| 3:1 — CD16bright vs CD16dim | −0.167 | −0.482 to 0.148 | 1.39 | 18 | 0.4500 ns |
| 3:1 — CD16bright vs CD16neg | −0.200 | −0.515 to 0.115 | 1.67 | 18 | 0.2997 ns |
| 3:1 — CD16dim vs CD16neg | −0.033 | −0.348 to 0.282 | 0.28 | 18 | 0.9899 ns |
| 1:1 — CD16bright vs CD16dim | 0.133 | −0.182 to 0.448 | 1.11 | 18 | 0.6265 ns |
| 1:1 — CD16bright vs CD16neg | −0.200 | −0.515 to 0.115 | 1.67 | 18 | 0.2997 ns |
| 1:1 — CD16dim vs CD16neg | −0.333 | −0.648 to −0.018 | 2.79 | 18 | 0.0362 * |
| ***Within-subset comparisons across E:T ratios*** | | | | | |
| CD16bright — No targets vs 3:1 | −0.100 | −0.415 to 0.215 | 0.84 | 18 | 0.7991 ns |
| CD16bright — No targets vs 1:1 | −0.133 | −0.448 to 0.182 | 1.11 | 18 | 0.6265 ns |
| CD16bright — 3:1 vs 1:1 | −0.033 | −0.348 to 0.282 | 0.28 | 18 | 0.9899 ns |
| CD16dim — No targets vs 3:1 | −0.100 | −0.415 to 0.215 | 0.84 | 18 | 0.7991 ns |
| CD16dim — No targets vs 1:1 | 0.167 | −0.148 to 0.482 | 1.39 | 18 | 0.4500 ns |
| CD16dim — 3:1 vs 1:1 | 0.267 | −0.048 to 0.582 | 2.23 | 18 | 0.1121 ns |
| CD16neg — No targets vs 3:1 | 0.067 | −0.248 to 0.382 | 0.56 | 18 | 0.9282 ns |
| CD16neg — No targets vs 1:1 | 0.033 | −0.282 to 0.348 | 0.28 | 18 | 0.9899 ns |
| CD16neg — 3:1 vs 1:1 | −0.033 | −0.348 to 0.282 | 0.28 | 18 | 0.9899 ns |

**Panel 2 — NKG2A density (gMFI)**

Design: Two-way ANOVA (CD16 subset × E:T ratio). Subsets: CD56dimCD16bright, CD56dimCD16dim, CD56dimCD16neg. E:T ratios: no targets, 3:1, 1:1. Post-hoc: Šidák's multiple comparisons test. n = 3 per group; 27 values total.

**Two-way ANOVA summary**

| **Source of variation** | **% of variation** | **F (DFn, DFd)** | **P value** | **Significance** |
| --- | --- | --- | --- | --- |
| CD16 subset | 16.13 | F (2, 18) = 1.98 | 0.1676 | ns |
| E:T ratio | 8.75 | F (2, 18) = 1.07 | 0.3630 | ns |
| Subset × E:T interaction | 1.67 | F (4, 18) = 0.10 | 0.9804 | ns |

**Šidák's multiple comparisons**

| **Comparison** | **Mean diff.** | **95% CI of diff.** | **t** | **DF** | **Adj. P value** |
| --- | --- | --- | --- | --- | --- |
| ***Between-subset comparisons at each E:T ratio*** | | | | | |
| No targets — CD16bright vs CD16dim | −462.7 | −1301 to 375.5 | 1.45 | 18 | 0.4149 ns |
| No targets — CD16bright vs CD16neg | −214.3 | −1053 to 623.9 | 0.67 | 18 | 0.8821 ns |
| No targets — CD16dim vs CD16neg | 248.3 | −589.9 to 1087 | 0.78 | 18 | 0.8298 ns |
| 3:1 — CD16bright vs CD16dim | −220.3 | −1059 to 617.9 | 0.69 | 18 | 0.8735 ns |
| 3:1 — CD16bright vs CD16neg | 29.67 | −808.5 to 867.9 | 0.09 | 18 | 0.9996 ns |
| 3:1 — CD16dim vs CD16neg | 250.0 | −588.2 to 1088 | 0.78 | 18 | 0.8270 ns |
| 1:1 — CD16bright vs CD16dim | −387.0 | −1225 to 451.2 | 1.22 | 18 | 0.5613 ns |
| 1:1 — CD16bright vs CD16neg | −141.0 | −979.2 to 697.2 | 0.44 | 18 | 0.9618 ns |
| 1:1 — CD16dim vs CD16neg | 246.0 | −592.2 to 1084 | 0.77 | 18 | 0.8336 ns |
| ***Within-subset comparisons across E:T ratios*** | | | | | |
| CD16bright — No targets vs 3:1 | −430.3 | −1269 to 407.9 | 1.35 | 18 | 0.4754 ns |
| CD16bright — No targets vs 1:1 | −205.3 | −1044 to 632.9 | 0.64 | 18 | 0.8944 ns |
| CD16bright — 3:1 vs 1:1 | 225.0 | −613.2 to 1063 | 0.71 | 18 | 0.8666 ns |
| CD16dim — No targets vs 3:1 | −188.0 | −1026 to 650.2 | 0.59 | 18 | 0.9162 ns |
| CD16dim — No targets vs 1:1 | −129.7 | −967.9 to 708.5 | 0.41 | 18 | 0.9699 ns |
| CD16dim — 3:1 vs 1:1 | 58.33 | −779.9 to 896.5 | 0.18 | 18 | 0.9971 ns |
| CD16neg — No targets vs 3:1 | −186.3 | −1025 to 651.9 | 0.58 | 18 | 0.9182 ns |
| CD16neg — No targets vs 1:1 | −132.0 | −970.2 to 706.2 | 0.41 | 18 | 0.9683 ns |
| CD16neg — 3:1 vs 1:1 | 54.33 | −783.9 to 892.5 | 0.17 | 18 | 0.9976 ns |

**Panel 3 — Degranulation stratified by NKG2A expression**

Design: Two-way ANOVA (CD16 subset × NKG2A expression). Subsets: CD56dimCD16bright, CD56dimCD16dim, CD56dimCD16neg. NKG2A status: NKG2A+, NKG2A−. Post-hoc: Šidák's multiple comparisons test. n = 3 per group; 18 values total.

**Two-way ANOVA summary**

| **Source of variation** | **% of variation** | **F (DFn, DFd)** | **P value** | **Significance** |
| --- | --- | --- | --- | --- |
| CD16 subset | 78.52 | F (2, 12) = 25.34 | <0.0001 | **** |
| NKG2A expression (+ vs −) | 0.17 | F (1, 12) = 0.11 | 0.7470 | ns |
| Subset × NKG2A interaction | 2.72 | F (2, 12) = 0.88 | 0.4410 | ns |

**Šidák's multiple comparisons**

| **Comparison** | **Mean diff.** | **95% CI of diff.** | **t** | **DF** | **Adj. P value** |
| --- | --- | --- | --- | --- | --- |
| ***Between-subset comparisons within each NKG2A status*** | | | | | |
| NKG2A+ — CD16bright vs CD16dim | −6.157 | −9.760 to −2.554 | 4.73 | 12 | 0.0015 ** |
| NKG2A+ — CD16bright vs CD16neg | 1.015 | −2.588 to 4.618 | 0.78 | 12 | 0.8339 ns |
| NKG2A+ — CD16dim vs CD16neg | 7.172 | 3.569 to 10.77 | 5.52 | 12 | 0.0004 *** |
| NKG2A− — CD16bright vs CD16dim | −4.316 | −7.919 to −0.714 | 3.32 | 12 | 0.0182 * |
| NKG2A− — CD16bright vs CD16neg | 0.553 | −3.049 to 4.156 | 0.43 | 12 | 0.9666 ns |
| NKG2A− — CD16dim vs CD16neg | 4.870 | 1.267 to 8.472 | 3.75 | 12 | 0.0084 ** |
| ***Within-subset comparisons: NKG2A+ vs NKG2A−*** | | | | | |
| CD16bright — NKG2A+ vs NKG2A− | −0.708 | −3.541 to 2.126 | 0.54 | 12 | 0.5963 ns |
| CD16dim — NKG2A+ vs NKG2A− | 1.133 | −1.700 to 3.966 | 0.87 | 12 | 0.4007 ns |
| CD16neg — NKG2A+ vs NKG2A− | −1.169 | −4.003 to 1.664 | 0.90 | 12 | 0.3863 ns |

*Significance: ns, not significant; * P < 0.05; ** P < 0.01; *** P < 0.001; **** P < 0.0001.*
