## Supplemental Table 17 for "CD56^dim^CD16^dim^ NK cells are the dominant effector cells against HIV-infected primary T-cells"

**Supplemental Table 17 for the statistical analysis of Figure 5C**

*Statistical analysis of NKG2D-blocking effects on CD107a degranulation responses of CD56^dim^CD16^bright^ and CD56^dim^CD16^dim^ NK cells against purified autologous productively HIV-1_SHM-1_ infected T-cells across four effector cell-to-target cell (E:T) ratios (1:4, 1:2, 1:1, and 2:1). Each subset was analyzed by an independent two-way ANOVA.*

**CD56^dim^CD16^bright^ panel**

**Table 1A. Two-way ANOVA summary (CD16^bright^).**

| **Source of Variation** | **% of total variation** | **SS** | **DF** | **MS** | **F (DFn, DFd)** | **P value** |
| --- | --- | --- | --- | --- | --- | --- |
| Interaction (E:T × treatment) | 7.99 | 4.480 | 3 | 1.493 | F(3, 16) = 10.06 | 0.0006 |
| E:T ratio | 17.13 | 9.593 | 3 | 3.198 | F(3, 16) = 21.54 | < 0.0001 |
| Treatment (isotype vs. anti-NKG2D) | 70.63 | 39.56 | 1 | 39.56 | F(1, 16) = 266.4 | < 0.0001 |
| Residual | — | 2.376 | 16 | 0.1485 | — | — |
| Total | — | 56.01 | 23 | — | — | — |

**Table 1B. Šidák's multiple comparisons test (CD16^bright^).**

| **E:T ratio** | **Comparison** | **Mean diff.** | **95% CI of diff.** | **Adjusted P value** | **Summary** |
| --- | --- | --- | --- | --- | --- |
| 1:4 | Isotype vs. anti-NKG2D Ab | 3.349 | 2.47 to 4.23 | < 0.0001 | **** |
| 1:2 | Isotype vs. anti-NKG2D Ab | 3.436 | 2.55 to 4.32 | < 0.0001 | **** |
| 1:1 | Isotype vs. anti-NKG2D Ab | 2.106 | 1.22 to 2.99 | < 0.0001 | **** |
| 2:1 | Isotype vs. anti-NKG2D Ab | 1.380 | 0.50 to 2.26 | 0.0018 | ** |

**CD56^dim^CD16^dim^ panel**

**Table 2A. Two-way ANOVA summary (CD16^dim^).**

| **Source of Variation** | **% of total variation** | **SS** | **DF** | **MS** | **F (DFn, DFd)** | **P value** |
| --- | --- | --- | --- | --- | --- | --- |
| Interaction (E:T × treatment) | 5.59 | 51.33 | 3 | 17.11 | F(3, 16) = 13.18 | 0.0001 |
| E:T ratio | 15.39 | 141.4 | 3 | 47.12 | F(3, 16) = 36.30 | < 0.0001 |
| Treatment (isotype vs. anti-NKG2D) | 76.75 | 704.8 | 1 | 704.8 | F(1, 16) = 543.0 | < 0.0001 |
| Residual | — | 20.77 | 16 | 1.298 | — | — |
| Total | — | 918.2 | 23 | — | — | — |

**Table 2B. Šidák's multiple comparisons test (CD16^dim^).**

| **E:T ratio** | **Comparison** | **Mean diff.** | **95% CI of diff.** | **Adjusted P value** | **Summary** |
| --- | --- | --- | --- | --- | --- |
| 1:4 | Isotype vs. anti-NKG2D Ab | 14.61 | 12.01 to 17.22 | < 0.0001 | **** |
| 1:2 | Isotype vs. anti-NKG2D Ab | 12.20 | 9.59 to 14.81 | < 0.0001 | **** |
| 1:1 | Isotype vs. anti-NKG2D Ab | 9.839 | 7.23 to 12.45 | < 0.0001 | **** |
| 2:1 | Isotype vs. anti-NKG2D Ab | 6.701 | 4.09 to 9.31 | < 0.0001 | **** |

*Two-way ANOVAs were performed in GraphPad Prism v10.6.1 with E:T ratio and treatment (isotype control vs. anti-NKG2D blocking antibody) as factors, separately for each NK cell subset. n = 24 values per subset (2 treatments × 4 E:T ratios × 3 replicates). Significance summary: * p < 0.05, ** p < 0.01, *** p < 0.001, **** p < 0.0001.*
