## Supplemental Table 18 for "CD56^dim^CD16^dim^ NK cells are the dominant effector cells against HIV-infected primary T-cells"

**Supplemental Table for the statistical analysis of Figure 6**

*Statistical analyses of NKG2D and NKp46 surface expression (gMFI) and CD107a degranulation responses of CD56^dim^CD16^bright^ and CD56^dim^CD16^dim^ NK cells to purified autologous productively HIV-1_SHM-1_ infected T-cells. All analyses were performed in GraphPad Prism v10.6.1 using two-way ANOVA with Šidák's multiple comparisons test.*

**Panel 6A: NKG2D gMFI**

**Table 1A. Two-way ANOVA summary.**

| **Source of Variation** | **% of total variation** | **SS** | **DF** | **MS** | **F (DFn, DFd)** | **P value** |
| --- | --- | --- | --- | --- | --- | --- |
| Interaction (E:T × subset) | 7.48 | 443760 | 4 | 110940 | F(4, 20) = 49.98 | < 0.0001 |
| E:T ratio | 20.80 | 1233770 | 4 | 308442 | F(4, 20) = 139.0 | < 0.0001 |
| NK cell subset | 70.97 | 4210504 | 1 | 4210504 | F(1, 20) = 1897 | < 0.0001 |
| Residual | — | 44396 | 20 | 2220 | — | — |

**Table 1B. Šidák's multiple comparisons test.**

| **Condition** | **Comparison** | **Mean diff.** | **95% CI of diff.** | **Adjusted P value** | **Summary** |
| --- | --- | --- | --- | --- | --- |
| No targets | CD16^bright^ vs. CD16^dim^ | −1182 | −1291 to −1073 | < 0.0001 | **** |
| 1:2 | CD16^bright^ vs. CD16^dim^ | −501.0 | −610 to −392 | < 0.0001 | **** |
| 1:1 | CD16^bright^ vs. CD16^dim^ | −558.3 | −667 to −449 | < 0.0001 | **** |
| 2:1 | CD16^bright^ vs. CD16^dim^ | −679.3 | −788 to −570 | < 0.0001 | **** |
| 4:1 | CD16^bright^ vs. CD16^dim^ | −826.0 | −935 to −717 | < 0.0001 | **** |

**Panel 6B: NKp46 gMFI on NKp46+ cells**

**Table 2A. Two-way ANOVA summary.**

| **Source of Variation** | **% of total variation** | **SS** | **DF** | **MS** | **F (DFn, DFd)** | **P value** |
| --- | --- | --- | --- | --- | --- | --- |
| Interaction (E:T × subset) | 5.91 | 16171 | 4 | 4043 | F(4, 20) = 9.432 | 0.0002 |
| E:T ratio | 40.66 | 111260 | 4 | 27815 | F(4, 20) = 64.89 | < 0.0001 |
| NK cell subset | 50.30 | 137634 | 1 | 137634 | F(1, 20) = 321.1 | < 0.0001 |
| Residual | — | 8573 | 20 | 428.6 | — | — |

**Table 2B. Šidák's multiple comparisons test.**

| **Condition** | **Comparison** | **Mean diff.** | **95% CI of diff.** | **Adjusted P value** | **Summary** |
| --- | --- | --- | --- | --- | --- |
| No targets | CD16^bright^ vs. CD16^dim^ | 45.67 | −2.28 to 93.61 | 0.0668 | ns |
| 1:2 | CD16^bright^ vs. CD16^dim^ | 178.3 | 130 to 226 | < 0.0001 | **** |
| 1:1 | CD16^bright^ vs. CD16^dim^ | 160.7 | 113 to 209 | < 0.0001 | **** |
| 2:1 | CD16^bright^ vs. CD16^dim^ | 148.3 | 100 to 196 | < 0.0001 | **** |
| 4:1 | CD16^bright^ vs. CD16^dim^ | 144.3 | 96.4 to 192 | < 0.0001 | **** |

**Panel 6C: % CD107a+ CD56^dim^ NK cells at various effector cell to target cell (E:T) ratios**

**Table 3A. Two-way ANOVA summary.**

| **Source of Variation** | **% of total variation** | **SS** | **DF** | **MS** | **F (DFn, DFd)** | **P value** |
| --- | --- | --- | --- | --- | --- | --- |
| Interaction (E:T × subset) | 9.50 | 161.9 | 3 | 53.97 | F(3, 16) = 69.57 | < 0.0001 |
| E:T ratio | 24.53 | 418.2 | 3 | 139.4 | F(3, 16) = 179.7 | < 0.0001 |
| NK cell subset | 65.24 | 1112 | 1 | 1112 | F(1, 16) = 1434 | < 0.0001 |
| Residual | — | 12.41 | 16 | 0.776 | — | — |

**Table 3B. Šidák's multiple comparisons test.**

| **E:T ratio** | **Comparison** | **Mean diff.** | **95% CI of diff.** | **Adjusted P value** | **Summary** |
| --- | --- | --- | --- | --- | --- |
| 1:2 | CD16^bright^ vs. CD16^dim^ | −20.42 | −22.43 to −18.40 | < 0.0001 | **** |
| 1:1 | CD16^bright^ vs. CD16^dim^ | −16.54 | −18.56 to −14.52 | < 0.0001 | **** |
| 2:1 | CD16^bright^ vs. CD16^dim^ | −10.45 | −12.47 to −8.44 | < 0.0001 | **** |
| 4:1 | CD16^bright^ vs. CD1^6dim^ | −7.047 | −9.06 to −5.03 | < 0.0001 | **** |

**Panel 6D: % CD107a+ stratified by NKp46 status at 1:1 E:T ratio**

**Table 4A. Two-way ANOVA summary.**

| **Source of Variation** | **% of total variation** | **SS** | **DF** | **MS** | **F (DFn, DFd)** | **P value** |
| --- | --- | --- | --- | --- | --- | --- |
| Interaction (subset × NKp46 status) | 0.21 | 0.639 | 1 | 0.639 | F(1, 8) = 0.388 | 0.5507 (ns) |
| NK cell subset | 94.96 | 290.0 | 1 | 290.0 | F(1, 8) = 176.2 | < 0.0001 |
| NKp46 expression status | 0.52 | 1.585 | 1 | 1.585 | F(1, 8) = 0.963 | 0.3552 (ns) |
| Residual | — | 13.17 | 8 | 1.646 | — | — |

**Table 4B. Šidák's multiple comparisons test.**

| **Subset** | **Comparison** | **Mean diff.** | **95% CI of diff.** | **Adjusted P value** | **Summary** |
| --- | --- | --- | --- | --- | --- |
| CD56^dim^CD16^bright^ | NKp46+ vs. NKp46- | 1.188 | −1.69 to 4.06 | 0.4952 | ns |
| CD56^dim^CD16^dim^ | NKp46+ vs. NKp46- | 0.265 | −2.61 to 3.14 | 0.9625 | ns |

*Significance summary: ns, not significant; * p < 0.05, ** p < 0.01, *** p < 0.001, **** p < 0.0001.*
