## Supplemental Table 19 for "CD56^dim^CD16^dim^ NK cells are the dominant effector cells against HIV-infected primary T-cells"

**Supplemental Table 19 for the statistical analysis of Figure 7A**

*Statistical analysis of CD107a degranulation responses of CD56^dim^CD16^bright^ and CD56^dim^CD16^dim^ NK cells against anti-gp120 antibody (Ab) (VRC01)-coated purified autologous productively HIV-1_SHM-1_ infected T-cells at a 1:1 effector cell-to-target cell (E:T) ratio of 1:1, across four VRC01 concentrations (0, 0.5, 1, and 2 μg/mL).*

**Table 1. Two-way ANOVA summary.**

| **Source of Variation** | **% of total variation** | **SS** | **DF** | **MS** | **F (DFn, DFd)** | **P value** |
| --- | --- | --- | --- | --- | --- | --- |
| Interaction (subset × VRC01 concentration) | 0.68 | 4.745 | 3 | 1.582 | F(3, 16) = 1.112 | 0.3732 (ns) |
| VRC01 concentration | 10.18 | 71.42 | 3 | 23.81 | F(3, 16) = 16.74 | < 0.0001 |
| NK cell subset | 85.91 | 602.9 | 1 | 602.9 | F(1, 16) = 424.0 | < 0.0001 |
| Residual | — | 22.75 | 16 | 1.422 | — | — |
| Total | — | 701.8 | 23 | — | — | — |

*Two-way ANOVA was performed in GraphPad Prism v10.6.1 with NK cell subset and VRC01 concentration as factors. n = 24 values (2 subsets × 4 VRC01 concentrations × 3 replicates). The non-significant interaction term (p = 0.3732) indicates that the subset effect is consistent across all VRC01 concentrations tested, with CD56^dim^CD16^dim^ NK cells showing higher ADCC responses than CD56^dim^CD16^bright^ cells at every concentration. NK cell subset accounted for 86% of the total variation, while VRC01 concentration accounted for 10%.*

**Table 2. Šidák's multiple comparisons test.**

| **VRC01 (μg/mL)** | **Comparison** | **Mean diff.** | **95% CI of diff.** | **Adjusted P value** | **Summary** |
| --- | --- | --- | --- | --- | --- |
| 0 | CD16^bright^ vs. CD16^dim^ | −9.268 | −12.00 to −6.54 | < 0.0001 | **** |
| 0.5 | CD16^bright^ vs. CD16^dim^ | −9.174 | −11.90 to −6.44 | < 0.0001 | **** |
| 1 | CD16^bright^ vs. CD16^dim^ | −10.29 | −13.02 to −7.56 | < 0.0001 | **** |
| 2 | CD16^bright^ vs. CD16^dim^ | −11.37 | −14.10 to −8.64 | < 0.0001 | **** |

*Significance summary: ns, not significant; * p < 0.05, ** p < 0.01, *** p < 0.001, **** p < 0.0001.*
