## Supplemental Table 20 for "CD56^dim^CD16^dim^ NK cells are the dominant effector cells against HIV-infected primary T-cells"

**Supplemental Table 20 for the statistical analysis of Figure 7—figure supplement 1**

*Degranulation of CD56^dim^CD16^bright^ and CD56^dim^CD16^dim^ NK cells in response to 1 µg/mL anti-CD16 antibody (clone 3G8) or isotype control at an effector cell-to-target cell ratio of 1:1*

Design: One-way ANOVA across four treatment conditions (CD56^dim^CD16^bright^ isotype, CD56^dim^CD16^bright^ 3G8, CD56^dim^CD16^dim^ isotype, CD56^dim^CD16^dim^ 3G8). Post-hoc: Šidák's multiple comparisons test for preselected pairwise contrasts. n = 3 per group; 12 values total*.*

**One-way ANOVA summary**

| **Source of variation** | **F (DFn, DFd)** | **P value** | **Significance** |
| --- | --- | --- | --- |
| Treatment (between groups) | F (3, 8) = 1887 | <0.0001 | **** |

*R² = 0.9986. Brown-Forsythe test for equality of variances: F(3,8)=1.015, p=0.4351 (ns).*

**Šidák's multiple comparisons**

| **Comparison** | **Mean diff.** | **95% CI of diff.** | **t** | **DF** | **Adj. P value** |
| --- | --- | --- | --- | --- | --- |
| CD16^bright^: isotype vs 3G8 | -40.89 | -43.79 to -37.98 | 42.27 | 8 | <0.0001 **** |
| CD16^dim^: isotype vs 3G8 | -57.84 | -60.75 to -54.93 | 59.80 | 8 | <0.0001 **** |
| CD16^bright^ 3G8 vs CD16^dim^ 3G8 | -20.27 | -23.17 to -17.36 | 20.95 | 8 | <0.0001 **** |

*Significance: ns, not significant; **** P < 0.0001.*
