## Supplemental Table 21 for "CD56^dim^CD16^dim^ NK cells are the dominant effector cells against HIV-infected primary T-cells"

**Supplemental Table 21 for the statistical analysis of Figure 7B**

*Statistical analysis of CD107a degranulation responses of CD56^dim^CD16^bright^ and CD56^dim^ CD16^dim^ NK cells against anti-gp120 (VRC01)-coated purified autologous productively*

*HIV-1_SHM-1_ infected T cells in the presence or absence of ADAM17 inhibition, across four VRC01 concentrations (0, 0.5, 1, and 2 μg/mL).*

**Table 1. Three-way ANOVA summary.**

| **Source of Variation** | **% of total variation** | **SS** | **DF** | **MS** | **F (DFn, DFd)** | **P value** |
| --- | --- | --- | --- | --- | --- | --- |
| VRC01 concentration | 37.38 | 174.4 | 3 | 58.14 | F(3, 16) = 309.0 | < 0.0001 |
| Treatment (DMSO vs. ADAM17 inhibitor) | 7.18 | 33.51 | 1 | 33.51 | F(1, 16) = 178.1 | < 0.0001 |
| NK cell subset | 26.25 | 122.5 | 1 | 122.5 | F(1, 16) = 557.9 | < 0.0001 |
| VRC01 × Treatment | 1.41 | 6.592 | 3 | 2.197 | F(3, 16) = 11.68 | 0.0003 |
| VRC01 × Subset | 1.27 | 5.903 | 3 | 1.968 | F(3, 16) = 8.962 | 0.0010 |
| Treatment × Subset | 23.18 | 108.2 | 1 | 108.2 | F(1, 16) = 492.6 | < 0.0001 |
| VRC01 × Treatment × Subset | 1.94 | 9.039 | 3 | 3.013 | F(3, 16) = 13.72 | 0.0001 |
| Residual | — | 3.513 | 16 | 0.2196 | — | — |

*Three-way ANOVA was performed in GraphPad Prism v10.6.1 with NK cell subset, treatment, and VRC01 concentration as factors. n = 48 values (2 subsets × 2 treatments × 4 VRC01 concentrations × 3 replicates). The Treatment × Subset interaction (F(1, 16) = 492.6, p < 0.0001) accounts for 23% of the total variation, indicating that ADAM17 inhibition has fundamentally different effects on the two NK cell subsets.*

**Table 2. Šidák's multiple comparisons test for pairwise comparisons.**

| **VRC01 (μg/mL)** | **Comparison** | **Mean diff.** | **95% CI of diff.** | **Adjusted P value** | **Summary** |
| --- | --- | --- | --- | --- | --- |
| (CD56^dim^CD16^bright^) |  |  |  |  |  |
| 0 | DMSO vs. ADAM-17 inhibitor | −1.001 | −2.45 to 0.447 | 0.7204 | ns |
| 0.5 | DMSO vs. ADAM-17 inhibitor | −0.313 | −1.76 to 1.14 | > 0.9999 | ns |
| 1 | DMSO vs. ADAM-17 inhibitor | −1.817 | −3.27 to −0.369 | 0.0029 | ** |
| 2 | DMSO vs. ADAM-17 inhibitor | −2.194 | −3.64 to −0.745 | 0.0002 | *** |
| (CD56^dim^CD16^dim^) |  |  |  |  |  |
| 0 | DMSO vs. ADAM-17 inhibitor | 2.202 | 0.754 to 3.65 | 0.0001 | *** |
| 0.5 | DMSO vs. ADAM-17 inhibitor | 5.704 | 4.26 to 7.15 | < 0.0001 | **** |
| 1 | DMSO vs. ADAM-17 inhibitor | 5.182 | 3.73 to 6.63 | < 0.0001 | **** |
| 2 | DMSO vs. ADAM-17 inhibitor | 5.605 | 4.16 to 7.05 | < 0.0001 | **** |

*Pairwise comparisons (Šidák-corrected) shown are those used to generate the per-sub-panel figure asterisks. Significance summary: ns, not significant; * p < 0.05, ** p < 0.01, *** p < 0.001, **** p < 0.0001.*
