## Supplemental Table 22 for "CD56^dim^CD16^dim^ NK cells are the dominant effector cells against HIV-infected primary T-cells"

**Supplemental Table 22 for the statistical analysis of Figure 7C**

Statistical analysis of data from Figure 7C. Change in CD56^dim^ NK cell subset frequency (CD56^dim^CD16^bright^ and CD56^dim^CD16^dim^) following 4-hour exposure to anti-gp120 (VRC01)-coated autologous HIV-1_SHM-1_ productively infected T-cells across VRC01 concentrations, under DMSO vehicle or ADAM17 inhibition.

*Design: separate two-way ANOVA per subset (treatment × VRC01 concentration). Post-hoc: Šidák's multiple comparisons test, DMSO vs ADAM17 inhibitor at each VRC01 concentration. n = 3 per condition.*

**CD56dimCD16bright (upper sub-panel)**

*Two-way ANOVA (ordinary)*

| **Source of variation** | **% of total variation** | **P value** | **F (DFn, DFd)** | **Significance** |
| --- | --- | --- | --- | --- |
| Interaction | 4.51 | 0.0459 | F (3, 16) = 3.338 | * |
| VRC01 concentration | 7.35 | 0.0090 | F (3, 16) = 5.437 | ** |
| Treatment | 80.93 | <0.0001 | F (1, 16) = 179.7 | **** |

Residual: DF = 16, MS = 7.565. Total SS = 1679, DF = 23.

*Šidák's multiple comparisons (DMSO vs ADAM17 inhibitor)*

| **VRC01 (µg/mL)** | **Mean diff. (DMSO − ADAM17 inh.)** | **95% CI of diff.** | **Adj. P** | **Summary** |
| --- | --- | --- | --- | --- |
| 0 | −14.77 | −19.53 to −10.01 | <0.0001 | **** |
| 0.5 | −10.23 | −14.99 to −5.473 | 0.0003 | *** |
| 1 | −14.93 | −19.69 to −10.17 | <0.0001 | **** |
| 2 | −20.27 | −25.03 to −15.51 | <0.0001 | **** |

Group means ± SD (n = 3): DMSO −8.03 ± 1.18 (0), −1.07 ± 2.21 (0.5), −2.13 ± 4.31 (1), −8.20 ± 3.76 (2); ADAM17 inhibitor 6.73 ± 3.76 (0), 9.17 ± 0.17 (0.5), 12.80 ± 2.10 (1), 12.07 ± 1.73 (2).

**CD56dimCD16dim (lower sub-panel)**

*Two-way ANOVA (ordinary)*

| **Source of variation** | **% of total variation** | **P value** | **F (DFn, DFd)** | **Significance** |
| --- | --- | --- | --- | --- |
| Interaction | 2.98 | 0.1751 | F (3, 16) = 1.871 | ns |
| VRC01 concentration | 5.57 | 0.0402 | F (3, 16) = 3.495 | * |
| Treatment | 82.96 | <0.0001 | F (1, 16) = 156.3 | **** |

Residual: DF = 16, MS = 4.103. Total SS = 772.8, DF = 23.

*Šidák's multiple comparisons (DMSO vs ADAM17 inhibitor)*

| **VRC01 (µg/mL)** | **Mean diff. (DMSO − ADAM17 inh.)** | **95% CI of diff.** | **Adj. P** | **Summary** |
| --- | --- | --- | --- | --- |
| 0 | 7.733 | 4.227 to 11.24 | 0.0003 | *** |
| 0.5 | 9.483 | 5.977 to 12.99 | <0.0001 | **** |
| 1 | 11.10 | 7.591 to 14.60 | <0.0001 | **** |
| 2 | 13.03 | 9.527 to 16.54 | <0.0001 | **** |

Group means ± SD (n = 3): DMSO 8.27 ± 2.68 (0), 6.20 ± 1.85 (0.5), 8.17 ± 2.85 (1), 11.27 ± 2.61 (2); ADAM17 inhibitor 0.53 ± 2.11 (0), −3.28 ± 0.43 (0.5), −2.93 ± 1.49 (1), −1.77 ± 0.66 (2).

Significance: * P < 0.05; ** P < 0.01; *** P < 0.001; **** P < 0.0001.
