## Supplemental Table 23 for "CD56^dim^CD16^dim^ NK cells are the dominant effector cells against HIV-infected primary T-cells"

**Supplemental Table 23 for the statistical analysis of Figure 8A**

*Degranulation of CD56^dim^CD16^bright^ and CD56^dim^CD16^dim^ NK cells against anti-gp120-coated purified autologous productively HIV-1_SHM-1_ infected T-cells under NKG2D blockade and/or ADAM17 inhibition*

Design: Two-way ANOVA (subset × treatment). Subset: CD56^dim^CD16^bright^ vs CD56^dim^CD16^dim^. Treatment: DMSO, anti-NKG2D antibody, ADAM17 inhibitor, ADAM17 inhibitor + anti-NKG2D. Post-hoc: Tukey's multiple comparisons test. n=3 per group; 24 values total.

**Two-way ANOVA summary**

| **Source of variation** | **% of variation** | **F (DFn, DFd)** | **P value** | **Significance** |
| --- | --- | --- | --- | --- |
| Subset (CD16^bright^ vs CD16^dim^) | 32.59 | F (1, 16) = 245.2 | <0.0001 | **** |
| Treatment | 38.81 | F (3, 16) = 97.33 | <0.0001 | **** |
| Subset × treatment interaction | 26.47 | F (3, 16) = 66.39 | <0.0001 | **** |

**Tukey's multiple comparisons**

| **Comparison** | **Mean diff.** | **95% CI of diff.** | **q** | **DF** | **Adj. P value** |
| --- | --- | --- | --- | --- | --- |
| ***Treatment comparisons within CD56^dim^CD16^bright^*** | | | | | |
| DMSO vs anti-NKG2D | 1.666 | 0.172 to 3.159 | 4.51 | 16 | 0.0263 * |
| DMSO vs ADAM17 inhibitor | −1.410 | −2.903 to 0.084 | 3.82 | 16 | 0.0679 ns |
| DMSO vs ADAM17 inhibitor+ anti-NKG2D | 1.509 | 0.015 to 3.003 | 4.09 | 16 | 0.0472 * |
| anti-NKG2D vs ADAM17 inhibitor | −3.075 | −4.569 to −1.582 | 8.33 | 16 | 0.0001 *** |
| anti-NKG2D vs ADAM17 inhibitor + anti-NKG2D | −0.157 | −1.650 to 1.337 | 0.42 | 16 | 0.9903 ns |
| ADAM17 inhibitor vs ADAM17 inhibitor + anti-NKG2D | 2.919 | 1.425 to 4.412 | 7.91 | 16 | 0.0002 *** |
| ***Treatment comparisons within CD56^dim^CD16^dim^*** | | | | | |
| DMSO vs anti-NKG2D | 7.363 | 5.869 to 8.857 | 19.94 | 16 | <0.0001 **** |
| DMSO vs ADAM17 inhibitor | 7.556 | 6.062 to 9.050 | 20.47 | 16 | <0.0001 **** |
| DMSO vs ADAM17 inhibitor + anti-NKG2D | 10.58 | 9.084 to 12.07 | 28.65 | 16 | <0.0001 **** |
| anti-NKG2D vs ADAM17 inhibitor | 0.193 | −1.301 to 1.687 | 0.52 | 16 | 0.9821 ns |
| anti-NKG2D vs ADAM17 inhibitor + anti-NKG2D | 3.215 | 1.721 to 4.709 | 8.71 | 16 | <0.0001 **** |
| ADAM17 inhibitor vs ADAM17 inhibitor + anti-NKG2D | 3.022 | 1.528 to 4.516 | 8.19 | 16 | 0.0001 *** |
| ***Subset comparison within each treatment (CD16^bright^ vs CD16^dim^)*** | | | | | |
| DMSO | −10.02 | −11.13 to −8.914 | 27.14 | 16 | <0.0001 **** |
| anti-NKG2D | −4.323 | −5.430 to −3.217 | 11.71 | 16 | <0.0001 **** |
| ADAM17 inhibitor | −1.055 | −2.162 to 0.052 | 2.86 | 16 | 0.0604 ns |
| ADAM17 inhibitor + anti-NKG2D | −0.952 | −2.059 to 0.155 | 2.58 | 16 | 0.0871 ns |

*Significance: ns, not significant; * P < 0.05; *** P < 0.001; **** P < 0.0001.*
