## Supplemental Table 24 for "CD56^dim^CD16^dim^ NK cells are the dominant effector cells against HIV-infected primary T-cells"

**Supplemental Table 24 for the statistical analysis of Figure 8B**

*Change in CD56^dim^ NK cell subset frequency (with anti-gp120-coated purified autologous productively HIV-1_SHM-1_ infected T-cells minus no targets) across treatments*

Design: Two-way ANOVA (subset × treatment). Subset: CD56^dim^CD16^bright^ Δ vs CD56^dim^CD16^dim^ Δ. Treatment: DMSO, ADAM17 inhibitor, anti-NKG2D antibody (Ab), anti-NKG2D Ab + ADAM17 inhibitor. Post-hoc: Šidák's multiple comparisons test. n=3 per group; 24 values total.

**Two-way ANOVA summary**

| **Source of variation** | **% of variation** | **F (DFn, DFd)** | **P value** | **Significance** |
| --- | --- | --- | --- | --- |
| Subset (CD16^bright^ Δ vs CD16^dim^ Δ) | 31.09 | F (1, 16) = 805.3 | <0.0001 | **** |
| Treatment | 2.77 | F (3, 16) = 23.93 | <0.0001 | **** |
| Subset × treatment interaction | 65.52 | F (3, 16) = 565.8 | <0.0001 | **** |

**Šidák's multiple comparisons**

| **Comparison** | **Mean diff.** | **95% CI of diff.** | **t** | **DF** | **Adj. P value** |
| --- | --- | --- | --- | --- | --- |
| ***Subset comparison within each treatment (CD16^bright^ Δ vs CD16^dim^ Δ)*** | | | | | |
| DMSO | −8.747 | −9.122 to −8.371 | 49.36 | 16 | <0.0001 **** |
| ADAM17 inhibitor | −1.430 | −1.806 to −1.054 | 8.07 | 16 | <0.0001 **** |
| anti-NKG2D Ab | 0.163 | −0.212 to 0.539 | 0.92 | 16 | 0.3703 ns |
| anti-NKG2D + ADAM17 inhibitor | −0.043 | −0.419 to 0.332 | 0.24 | 16 | 0.8099 ns |
| ***Treatment comparisons within CD56^dim^CD16^bright^ Δ*** | | | | | |
| DMSO vs ADAM17 inhibitor | −4.067 | −4.598 to −3.535 | 22.95 | 16 | <0.0001 **** |
| DMSO vs anti-NKG2D Ab | −5.333 | −5.865 to −4.802 | 30.10 | 16 | <0.0001 **** |
| DMSO vs anti-NKG2D Ab + ADAM17 inhibitor | −5.267 | −5.798 to −4.735 | 29.72 | 16 | <0.0001 **** |
| ADAM17 inhibitor vs anti-NKG2D Ab | −1.267 | −1.798 to −0.735 | 7.15 | 16 | <0.0001 **** |
| ADAM-17 inhibitor vs anti-NKG2D Ab + ADAM17 inhibitor | −1.200 | −1.731 to −0.669 | 6.77 | 16 | <0.0001 **** |
| anti-NKG2D Ab vs anti-NKG2D + ADAM17 inhibitor | 0.067 | −0.465 to 0.598 | 0.38 | 16 | 0.9994 ns |
| ***Treatment comparisons within CD56^dim^CD16^dim^ Δ*** | | | | | |
| DMSO vs ADAM17 inhibitor | 3.250 | 2.719 to 3.781 | 18.34 | 16 | <0.0001 **** |
| DMSO vs anti-NKG2D Ab | 3.577 | 3.045 to 4.108 | 20.19 | 16 | <0.0001 **** |
| DMSO vs anti-NKG2D Ab + ADAM17 inhibitor | 3.437 | 2.905 to 3.968 | 19.40 | 16 | <0.0001 **** |
| ADAM17 inhibitor vs anti-NKG2D Ab | 0.327 | −0.205 to 0.858 | 1.84 | 16 | 0.4087 ns |
| ADAM17 i inhibitor vs anti-NKG2D + ADAM17 inhibitor | 0.187 | −0.345 to 0.718 | 1.05 | 16 | 0.8900 ns |
| anti-NKG2D Ab vs anti-NKG2D + ADAM17 inhibitor | −0.140 | −0.671 to 0.391 | 0.79 | 16 | 0.9695 ns |

*Significance: ns, not significant; **** P < 0.0001.*
