## Supplemental Table 25 for "CD56^dim^CD16^dim^ NK cells are the dominant effector cells against HIV-infected primary T-cells"

**Supplemental Table 25 for the statistical analysis of Figure 9— CD56^dim^CD16^bright^ panel**

*Serial degranulation of CD56^dim^CD16^bright^ NK cells against anti-gp120-coated purified autologous productively HIV-1_SHM-1_ infected T-cells under DMSO vehicle or ADAM17 inhibition, scored as the number of degranulation events per cell within 60 minutes*

Design: Two-way ANOVA (treatment × number of degranulations). Treatment: DMSO, ADAM17 inhibitor. Number of degranulations per cell over 60 minutes: 0, 1, 2, 3. Post-hoc: Šidák's multiple comparisons, DMSO vs ADAM17 inhibitor at each degranulation count. n=3 per group; 24 values total.

**Two-way ANOVA summary**

| **Source of variation** | **% of variation** | **F (DFn, DFd)** | **P value** | **Significance** |
| --- | --- | --- | --- | --- |
| Number of degranulations (0, 1, 2, 3) | 99.60 | F (3, 16) = 6145 | <0.0001 | **** |
| Treatment (DMSO vs ADAM17 inhibitor) | <0.001 | F (1, 16) = 0.13 | 0.7233 | ns |
| Degranulations × treatment interaction | 0.32 | F (3, 16) = 19.50 | <0.0001 | **** |

*The treatment main effect tests whether the overall mean across all four degranulation categories differs between DMSO and ADAM17 inhibition. Because each cell falls into exactly one of the 0, 1, 2, or 3 categories and the four percentages within each treatment sum to ~100, the two treatment means are mathematically constrained to be nearly identical (DMSO mean = 25.03; ADAM17 inhibitor mean = 25.23), and the corresponding p value is uninformative. The biologically meaningful effects are captured by the degranulation main effect, the degranulation × treatment interaction, and the post hoc comparisons of DMSO versus ADAM17 inhibition at each degranulation count.*

**DMSO vs ADAM-17 inhibition at each degranulation count**

| **Degranulations** | **Mean diff. (DMSO − ADAM17 inhibito**r**)** | **95% CI of diff.** | **t** | **DF** | **Adj. P value** |
| --- | --- | --- | --- | --- | --- |
| 0 | 6.581 | 4.322 to 8.841 | 6.17 | 16 | <0.0001 **** |
| 1 | −1.471 | −3.731 to 0.789 | 1.38 | 16 | 0.1866 ns |
| 2 | −1.553 | −3.813 to 0.707 | 1.46 | 16 | 0.1645 ns |
| 3 | −4.326 | −6.585 to −2.066 | 4.06 | 16 | 0.0009 *** |

**Group means (% of CD56^dim^CD16^bright^ NK cells)**

| **Treatment** | **0 degranulations (mean %)** | **1 degranulation (mean %)** | **2 degranulations (mean %)** | **3 degranulations (mean %)** | **n per cell** | **SE of diff.** |
| --- | --- | --- | --- | --- | --- | --- |
| DMSO | 91.06 | 5.17 | 3.10 | 0.79 | 3 | 1.066 |
| ADAM-17 inhibitor | 84.48 | 6.64 | 4.65 | 5.12 | 3 | 1.066 |

*Significance: ns, not significant; *** P < 0.001; **** P < 0.0001.*
