## Supplemental Table 26 for "CD56^dim^CD16^dim^ NK cells are the dominant effector cells against HIV-infected primary T-cells"

**Supplemental Table 26 for the statistical analysis of** **Figure 9— CD56^dim^CD16^dim^ panel**

*Serial degranulation of CD56^dim^CD16^dim^ NK cells against anti-gp120-coated purified autologous productively HIV-1_SHM-1_ infected T-cells under DMSO vehicle or ADAM17 inhibition, scored as the number of degranulation events per cell within 60 minutes.*

Design: Two-way ANOVA (treatment × number of degranulations). Treatment: DMSO, ADAM17 inhibitor. Number of degranulations per cell over 60 minutes: 0, 1, 2, 3. Post-hoc: DMSO vs ADAM-17 inhibitor at each degranulation count. n=3 per group; 24 values total.

**Two-way ANOVA summary**

| **Source of variation** | **% of variation** | **F (DFn, DFd)** | **P value** | **Significance** |
| --- | --- | --- | --- | --- |
| Number of degranulations (0, 1, 2, 3) | 91.74 | F (3, 16) = 2619 | <0.0001 | **** |
| Treatment (DMSO vs ADAM17 inhibitor) | <0.001 | F (1, 16) = 0.0007 | 0.9788 | ns |
| Degranulations × treatment interaction | 8.08 | F (3, 16) = 230.6 | <0.0001 | **** |

*The treatment main effect tests whether the overall mean across all four degranulation categories differs between DMSO and ADAM17 inhibition. Because each cell falls into exactly one of the 0, 1, 2, or 3 categories and the four percentages within each treatment sum to ~100, the two treatment means are mathematically constrained to be nearly identical (DMSO mean = 25.05; ADAM17* i*nhibitor mean = 25.04), and the corresponding p value is uninformative. The biologically meaningful effects are captured by the degranulation main effect, the degranulation × treatment interaction, and the post hoc comparisons of DMSO versus ADAM17 inhibition at each degranulation count.*

**DMSO vs ADAM-17 inhibition at each degranulation count**

| **Degranulations** | **Mean diff. (DMSO − ADAM17 inhibito**r**)** | **95% CI of diff.** | **q** | **DF** | **Adj. P value** |
| --- | --- | --- | --- | --- | --- |
| 0 | −30.57 | −33.44 to −27.71 | 22.64 | 16 | <0.0001 **** |
| 1 | 9.98 | 7.12 to 12.84 | 7.39 | 16 | <0.0001 **** |
| 2 | 7.75 | 4.89 to 10.62 | 5.74 | 16 | <0.0001 **** |
| 3 | 12.92 | 10.05 to 15.78 | 9.57 | 16 | <0.0001 **** |

**Group means (% of CD56^dim^CD16^dim^ NK cells)**

| **Treatment** | **0 degranulations (mean %)** | **1 degranulation (mean %)** | **2 degranulations (mean %)** | **3 degranulations (mean %)** | **n per cell** | **SE of diff.** |
| --- | --- | --- | --- | --- | --- | --- |
| DMSO | 61.56 | 13.55 | 9.95 | 15.16 | 3 | 1.350 |
| ADAM17 inhibitor | 92.13 | 3.57 | 2.19 | 2.24 | 3 | 1.350 |

*Significance: ns, not significant; **** P < 0.0001.*
