## Supplemental Table 27 for "CD56^dim^CD16^dim^ NK cells are the dominant effector cells against HIV-infected primary T-cells"

**Supplemental Table 27: Antibodies and Reagents Used in the Study**

| **For Flow Cytometry** | | | | | | |
| --- | --- | --- | --- | --- | --- | --- |
| **Marker/Antigen** | | **Fluorochrome** | **Clone** | **Vendor** | | **Catalog #** |
| human CD56 | | APC | NCAM16.2 | BD Bioscience | | 341025 |
| human CD56 | | BV786 | NCAM16.2 | BD Bioscience | | 564058 |
| human CD16 | | BV605 | B73.1 | Biolegend | | 360728 |
| human CD3 | | AF700 | UCHT1 | Biolegend | | 300424 |
| human CD3 | | PE Dazzle | UCHT1 | Biolegend | | 300449 |
| human CD107a | | PE | H4A3 | Biolegend | | 328608 |
| human CD107a | | BV421 | H4A3 | Biolegend | | 328625 |
| human CD107a | | FITC | H4A3 | Biolegend | | 328606 |
| human CD107a | | unlabeled | H4A3 | Biolegend | | 328601 |
| human NKG2D | | FITC | 1D11 | Biolegend | | 320820 |
| human NKp80 | | PE-Vio770 | REA845 | Miltenyi Biotec | | 130-112-781 |
| human IFNγ | | FITC | 4S.B3 | Biolegend | | 502505 |
| human GZMA | | PE/Cyanine7 | CB9 | Biolegend | | 507221 |
| human GZMB | | AF647 | GB11 | Biolegend | | 515405 |
| human PFN | | PF488P | Pf-344 | MabTech | | 3465-71-100T |
| human CD4 | | FITC | OKT4 | Biolegend | | 317408 |
| HIV-1 p24 | | RD1 | KC57 | Beckman Coulter | | 6604667 |
| human NKG2D-Fc chimera | | unlabeled |  | R&D Systems | | 1299-NK-050 |
| human IgG Fc | | APC |  | Jackson ImmunoResearch | | 109-136-190 |
| human CD158a | | APC-Vio770 | REA284 | Miltenyi Biotec | | 130-118-345 |
| human CD158b | | PE-Vio615 | REA1006 | Miltenyi Biotec | | 130-116-954 |
| human CD158e | | APC | DX9 | Biolegend | | 312715 |
| human NKG2A | | FITC | S19004C | Biolegend | | 375127 |
| **For Cell Sorting** | | | | | | |
| human CD56 | | FITC | NCAM | Biolegend | | 318303 |
| human CD16 Fab Fragments | | unlabeled | 3G8 | Mybioscource.com | | MBS666071 |
| Affinity Pure Fab Fragment Goat anti-mouse IgG (H+L) | | AF647 |  | Jackson ImmunoResearch | | 115-607-003 |
| human CD3 | | AF700 | UCHT1 | Biolegend | | 300424 |
| **For Mean Fluorescent Intensity Studies** | | | | | | |
| human NKG2D | | APC | 1D11 | Biolegend | | 320808 |
| human NKp46 | | PE-Cy7 | 9E2 | Biolegend | | 331915 |
| **Additional Reagents** | | | | | | |
| **Reagent** | **Final Concentration Used** | | **Vendor** | | **Catalog #** | |
| Aqua Fluorescence Reactive Dye | Aqua | | Invitrogen | | L34965 | |
| Cytotoxicity Assay Kit |  | | AbCam | | ab270780 | |
| GW280264X | 1.74 µM | | Aobious | | AOB3632 | |
| Monensin | 2.05 µM | | Becton Dickinson | | BDB554724 | |
| NKG2D blocking antibody | 10 mg/mL | | R&D Systems | | MAB139-100 | |
| Mouse IgG_1_ Isotype Control | 10 mg/mL | | R&D Systems | | MAB002 | |
| VRC01 | 0.5-2 mg/mL | | BEI Resources | | ARP-12033 | |
