## Supplementary figures and images for "CD56^dim^CD16^dim^ NK cells are the dominant effector cells against HIV-infected primary T-cells"

### Figure 1-figure supplement 1

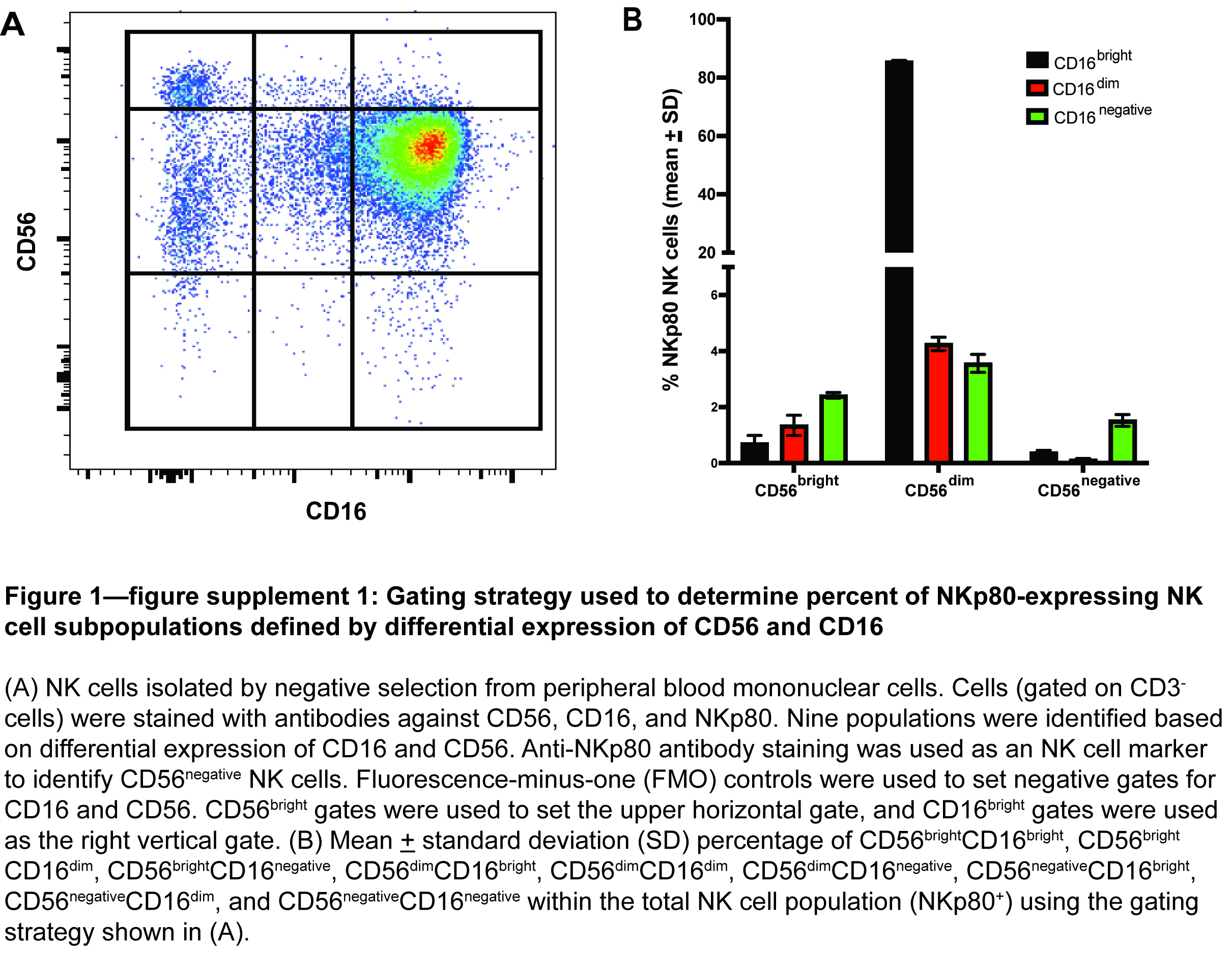

### Figure 1-figure supplement 2

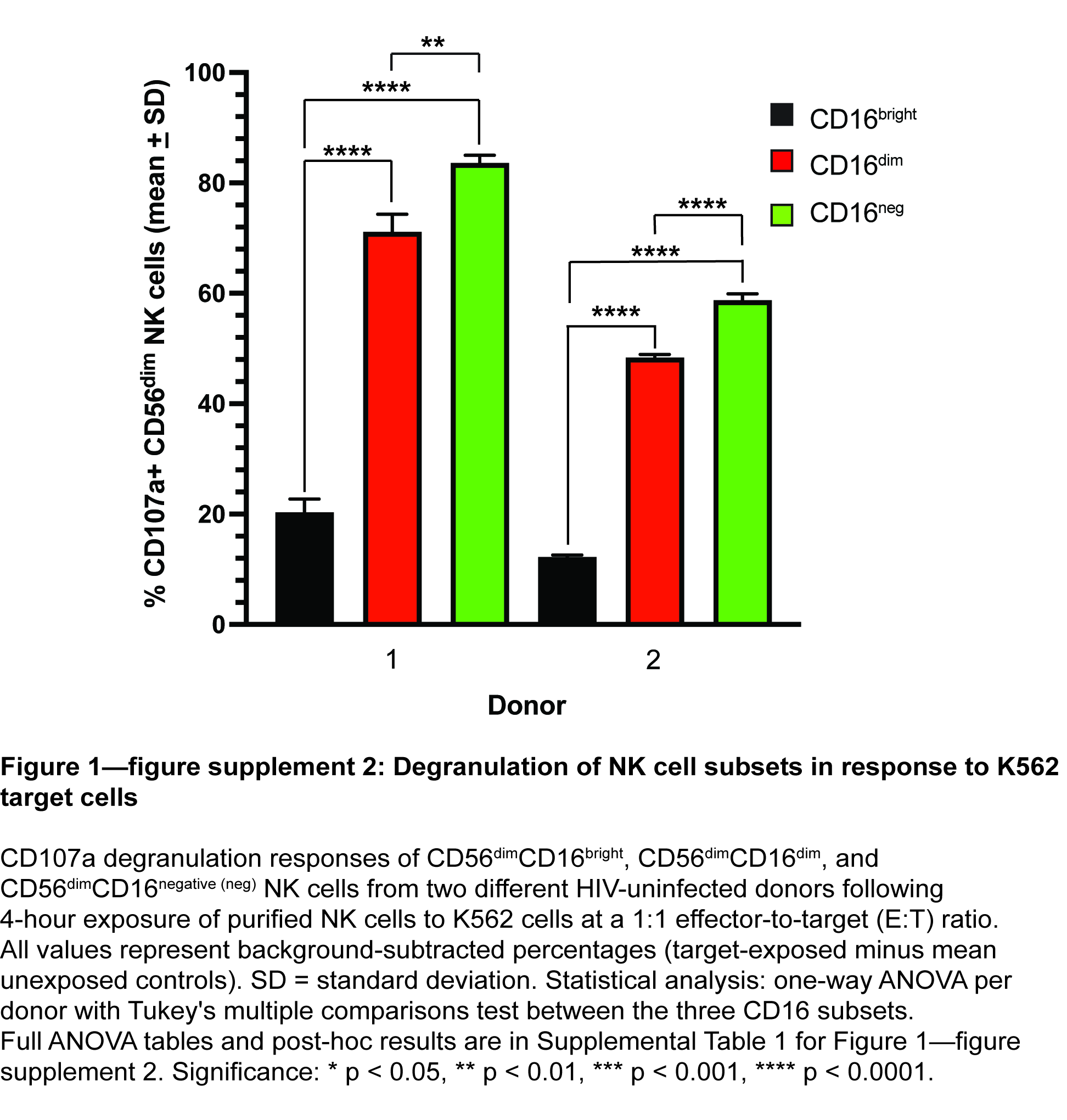

### Figure 1-figure supplement 3

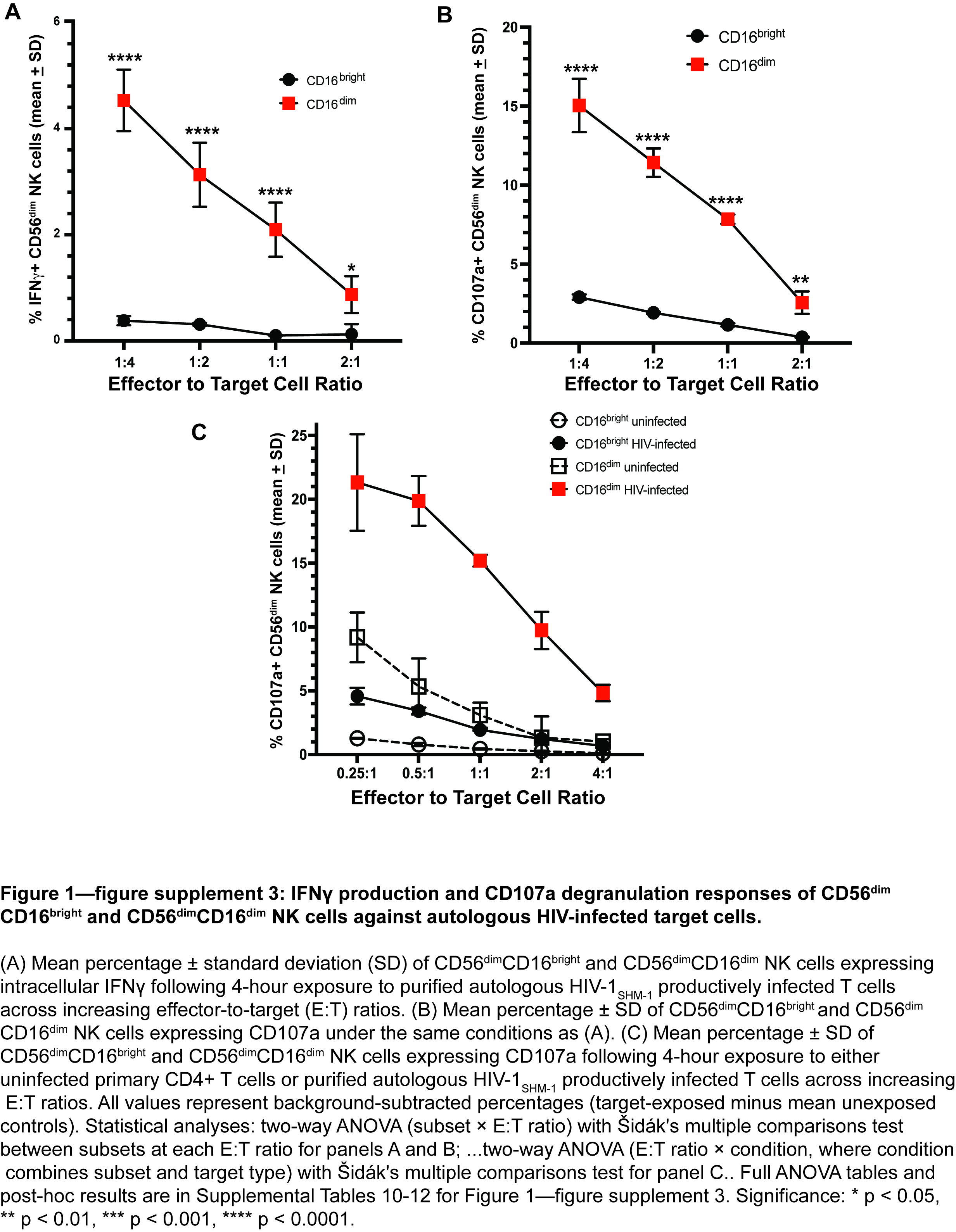

### Figure 1-figure supplement 4

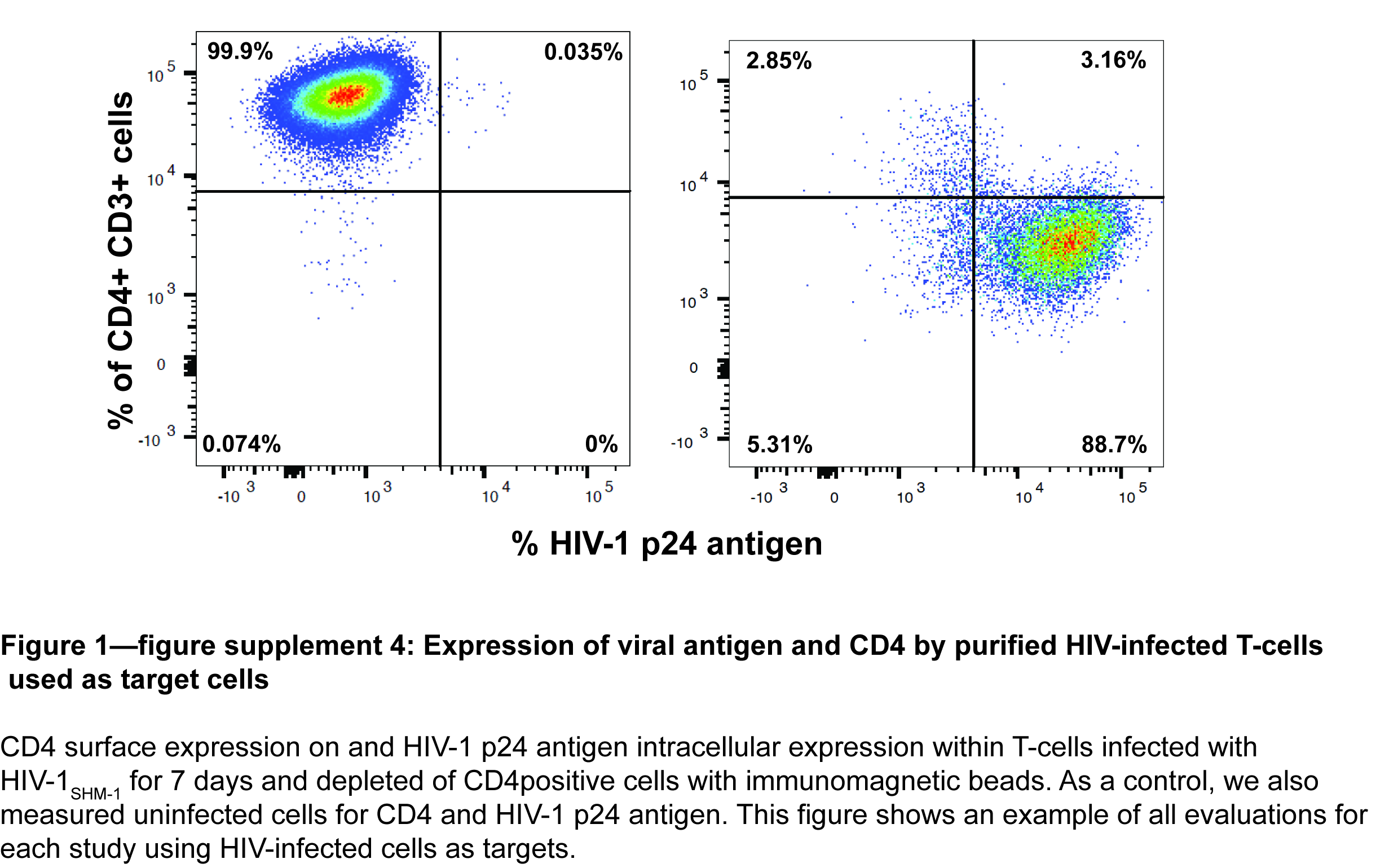

### Figure 1-figure supplement 5

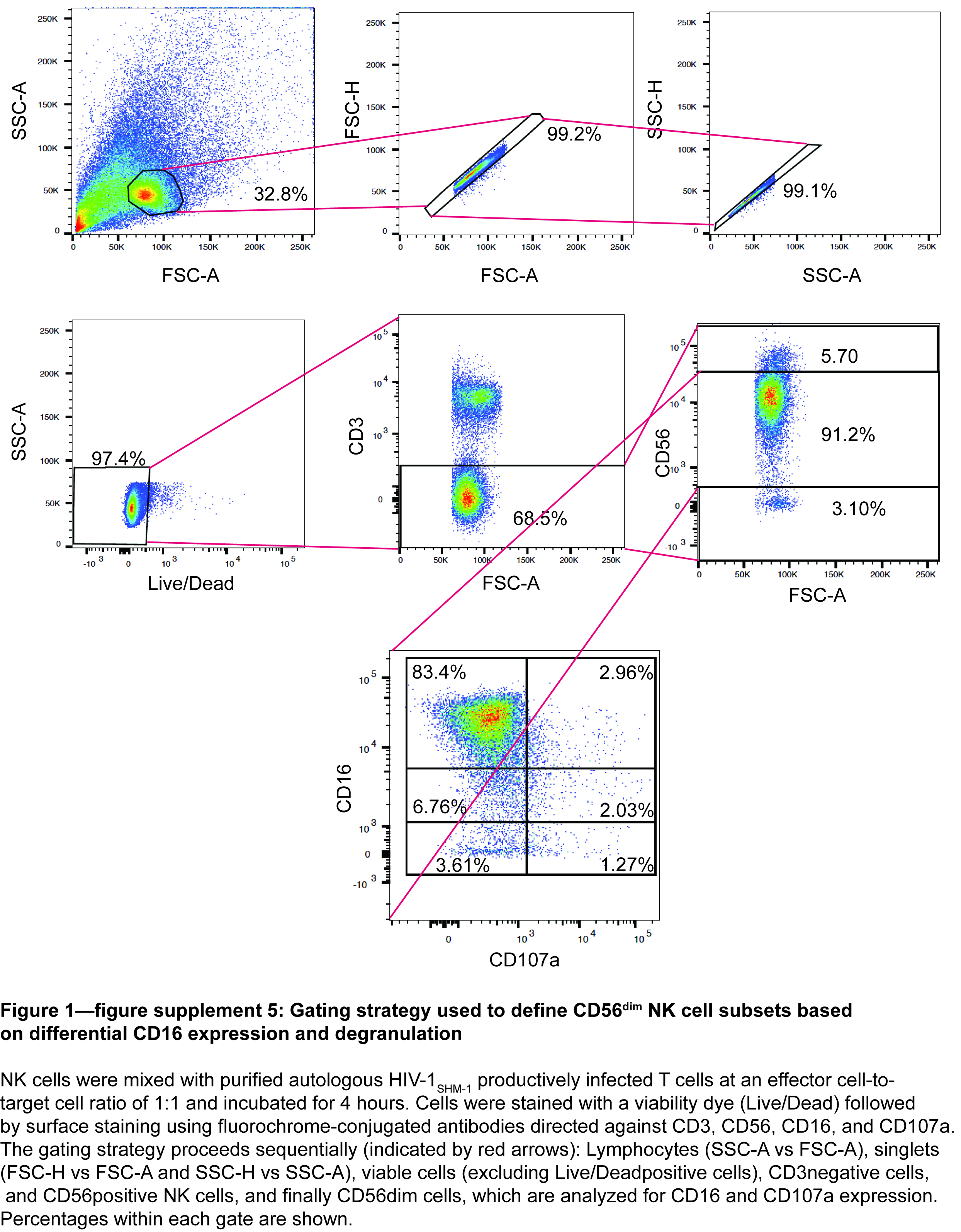

### Figure 1-figure supplement 6

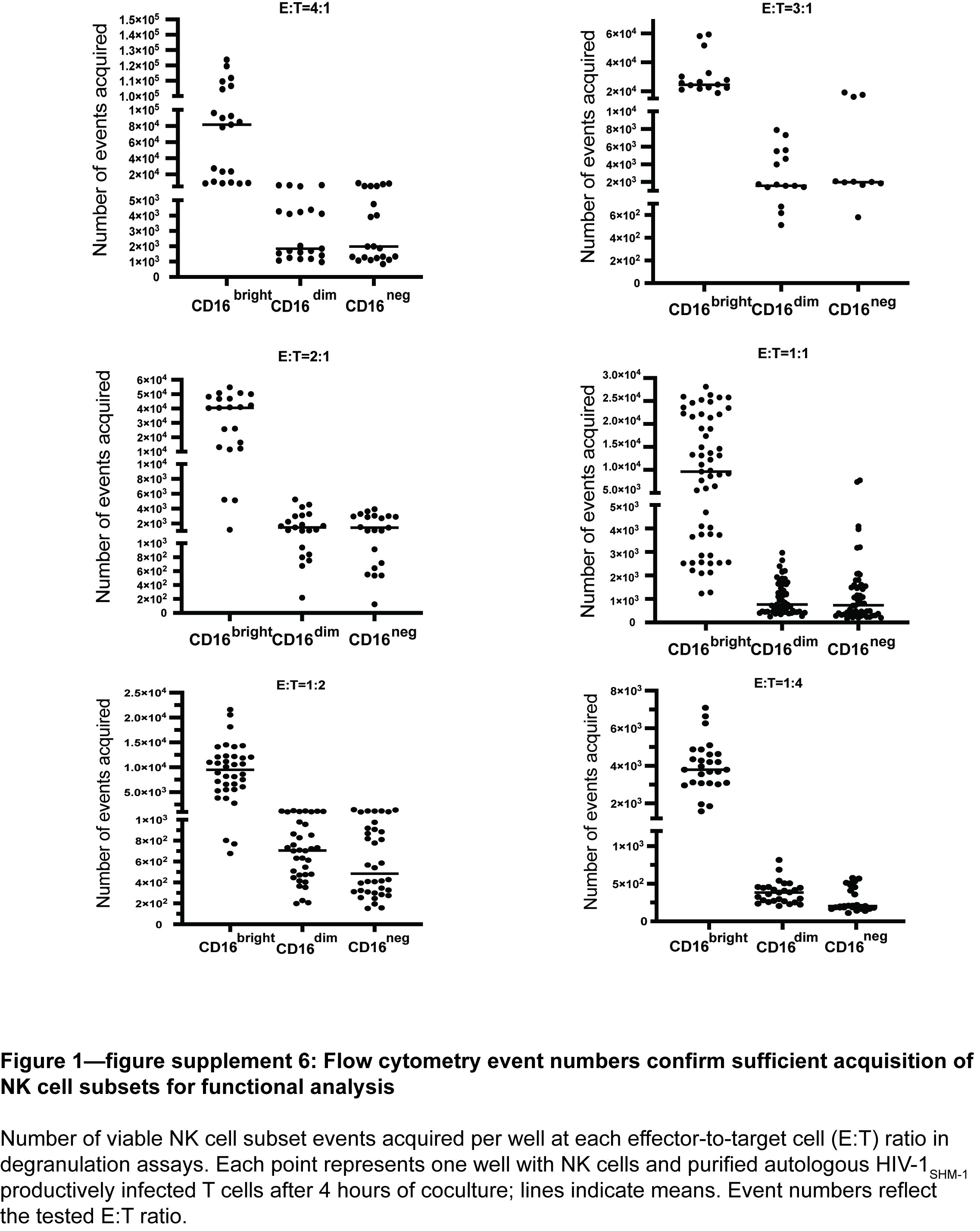

### Figure 4-figure supplement 1

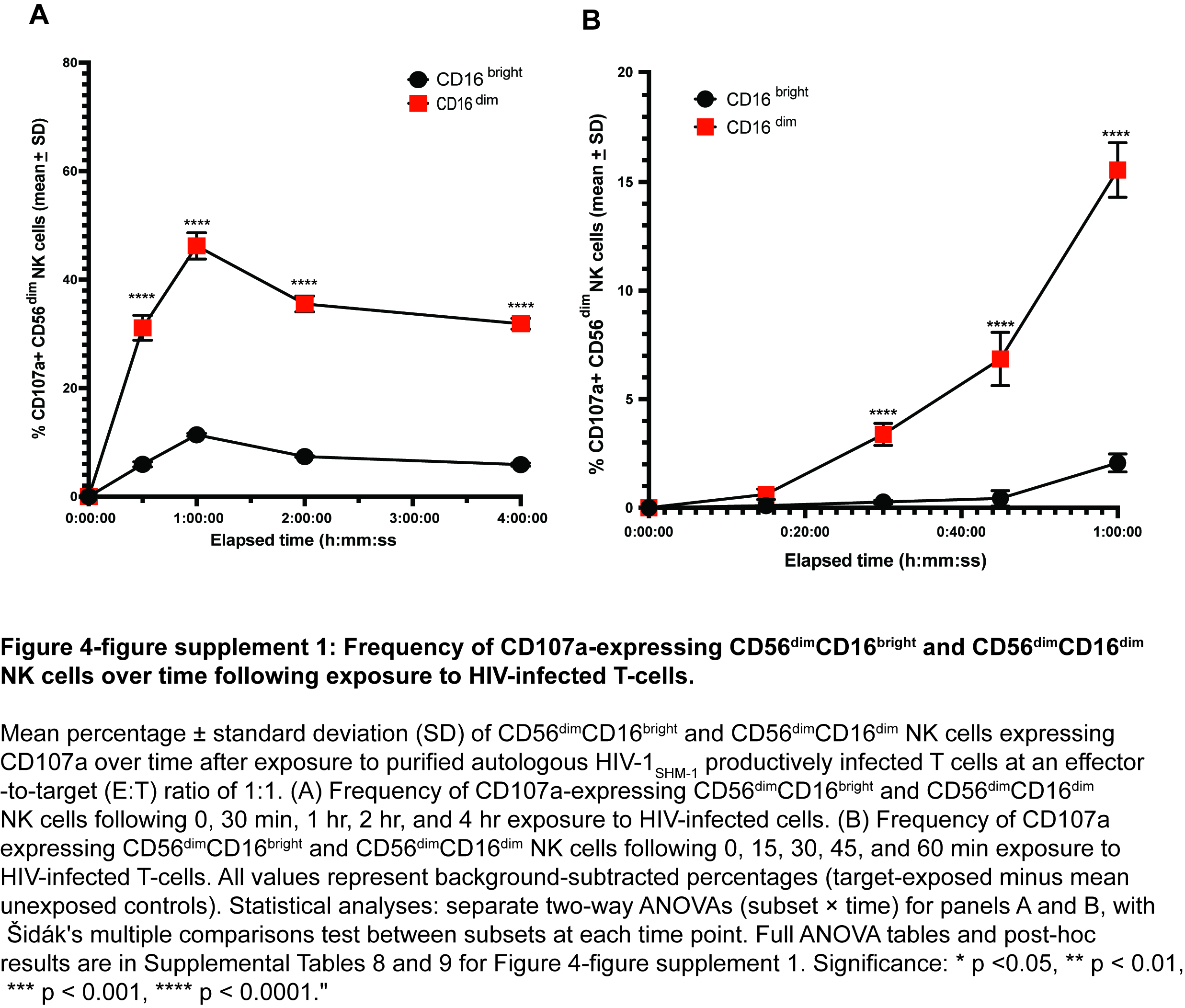

### Figure 5-figure supplement 1

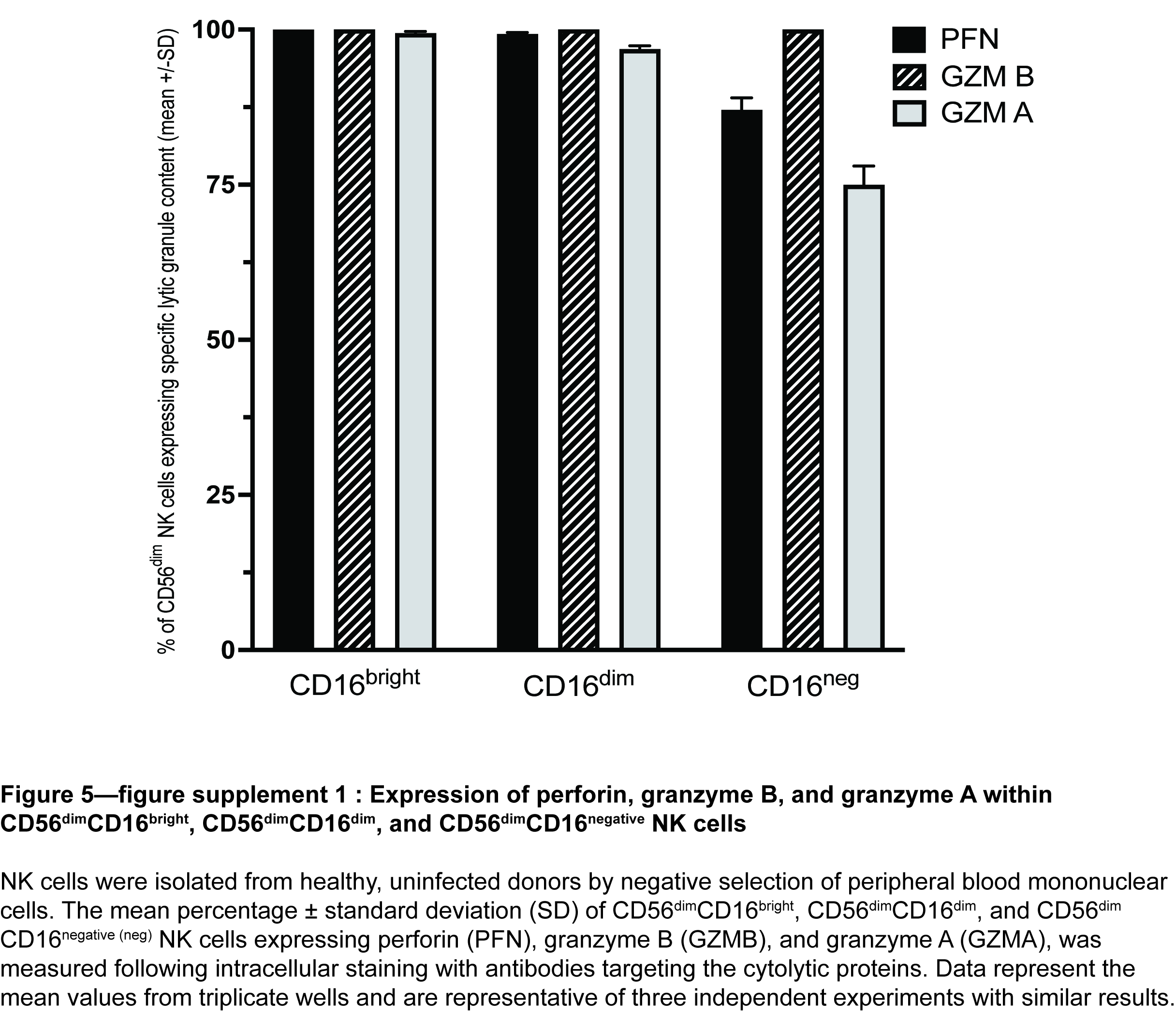

### Figure 5-figure supplement 2

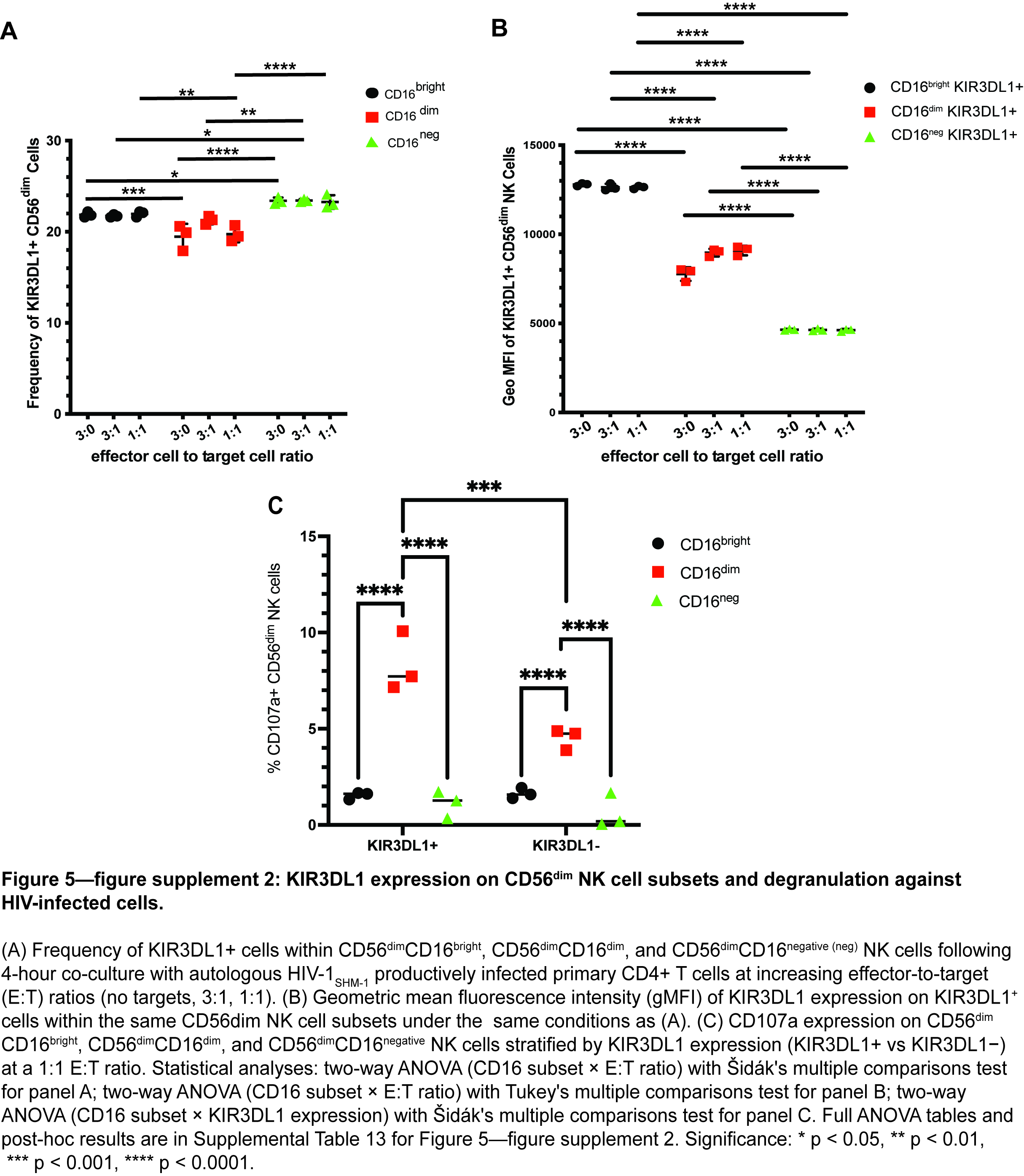

### Figure 5-figure supplement 3

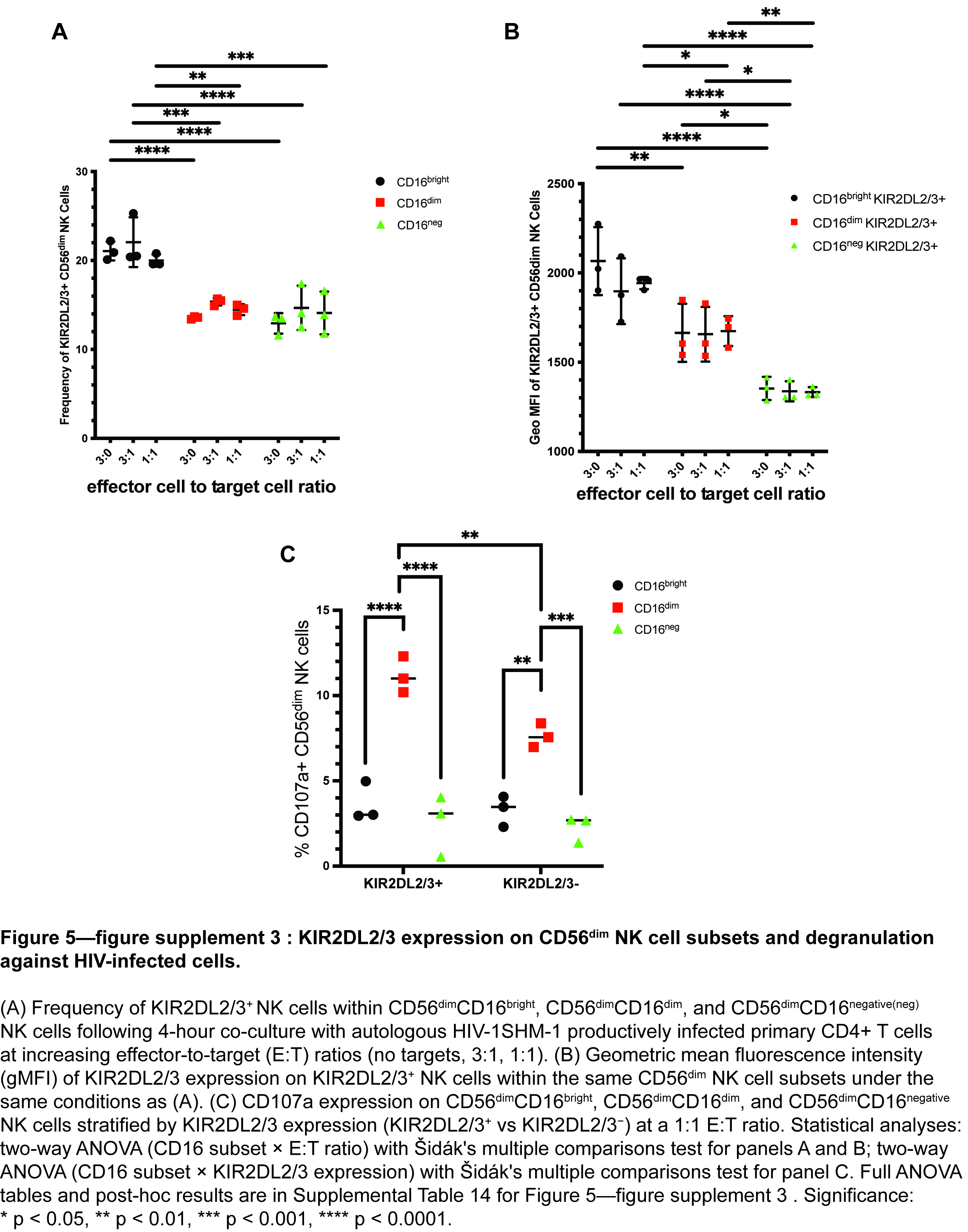

### Figure 5-figure supplement 4

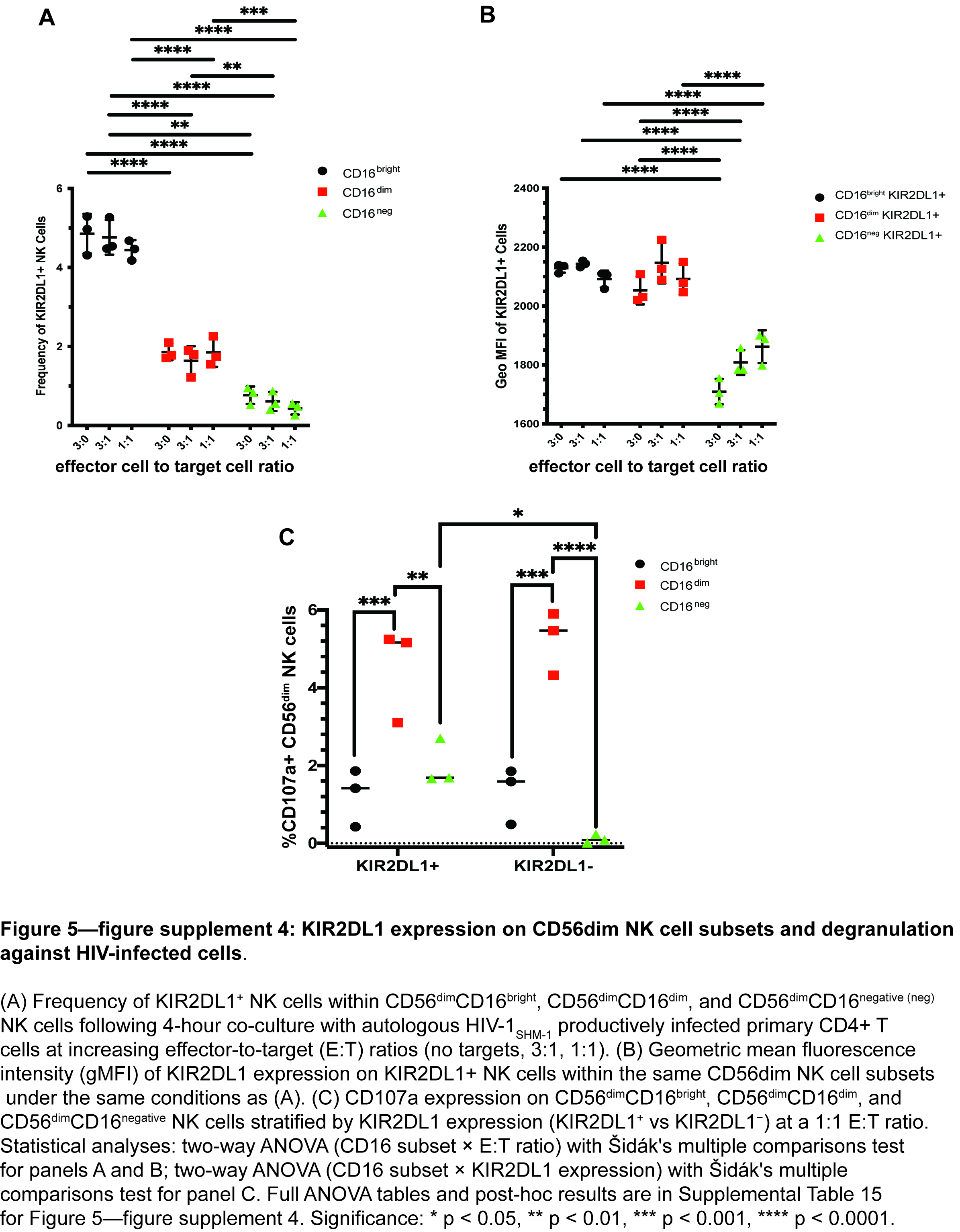

### Figure 5-figure supplement 5

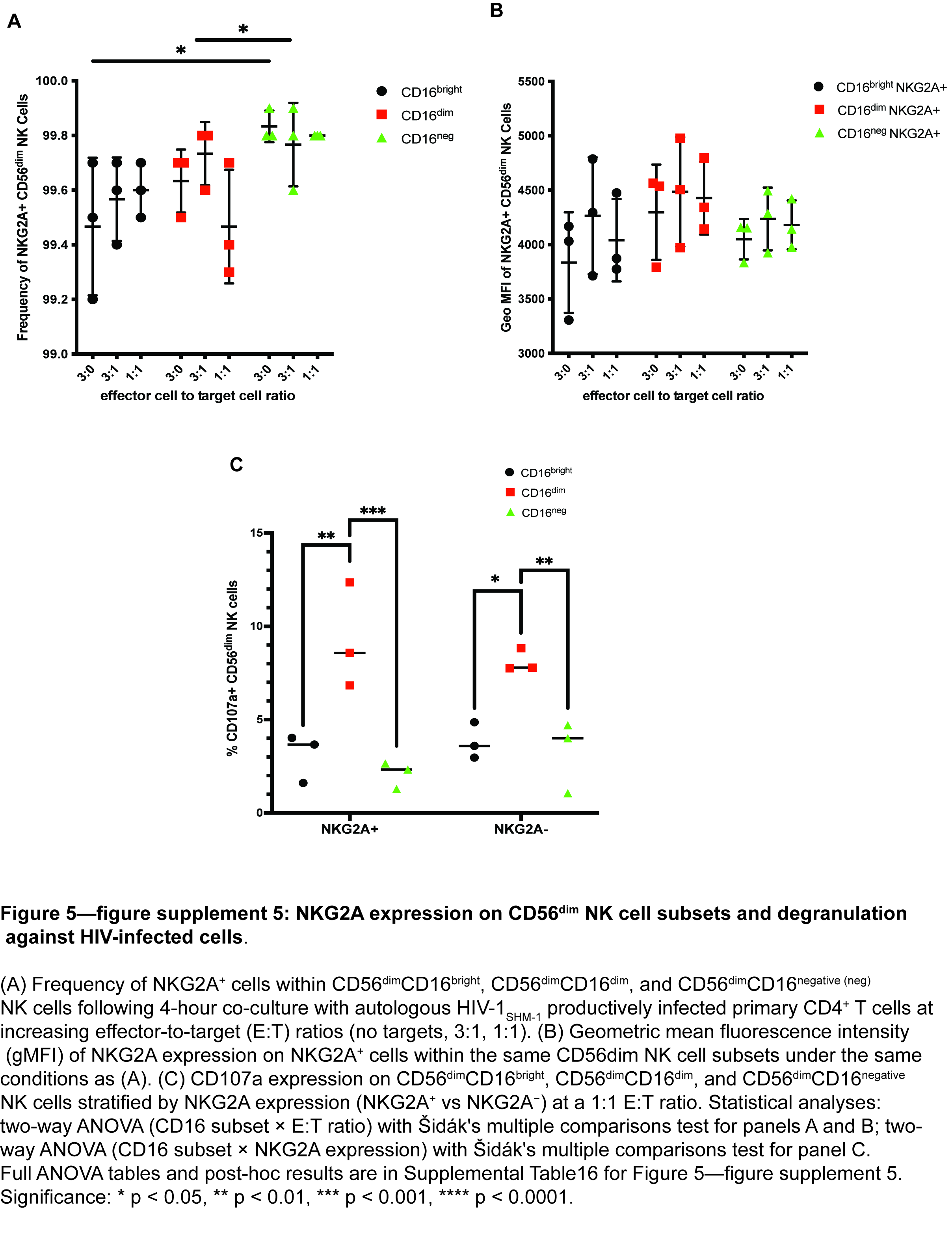

### Figure 7-figure supplement 1

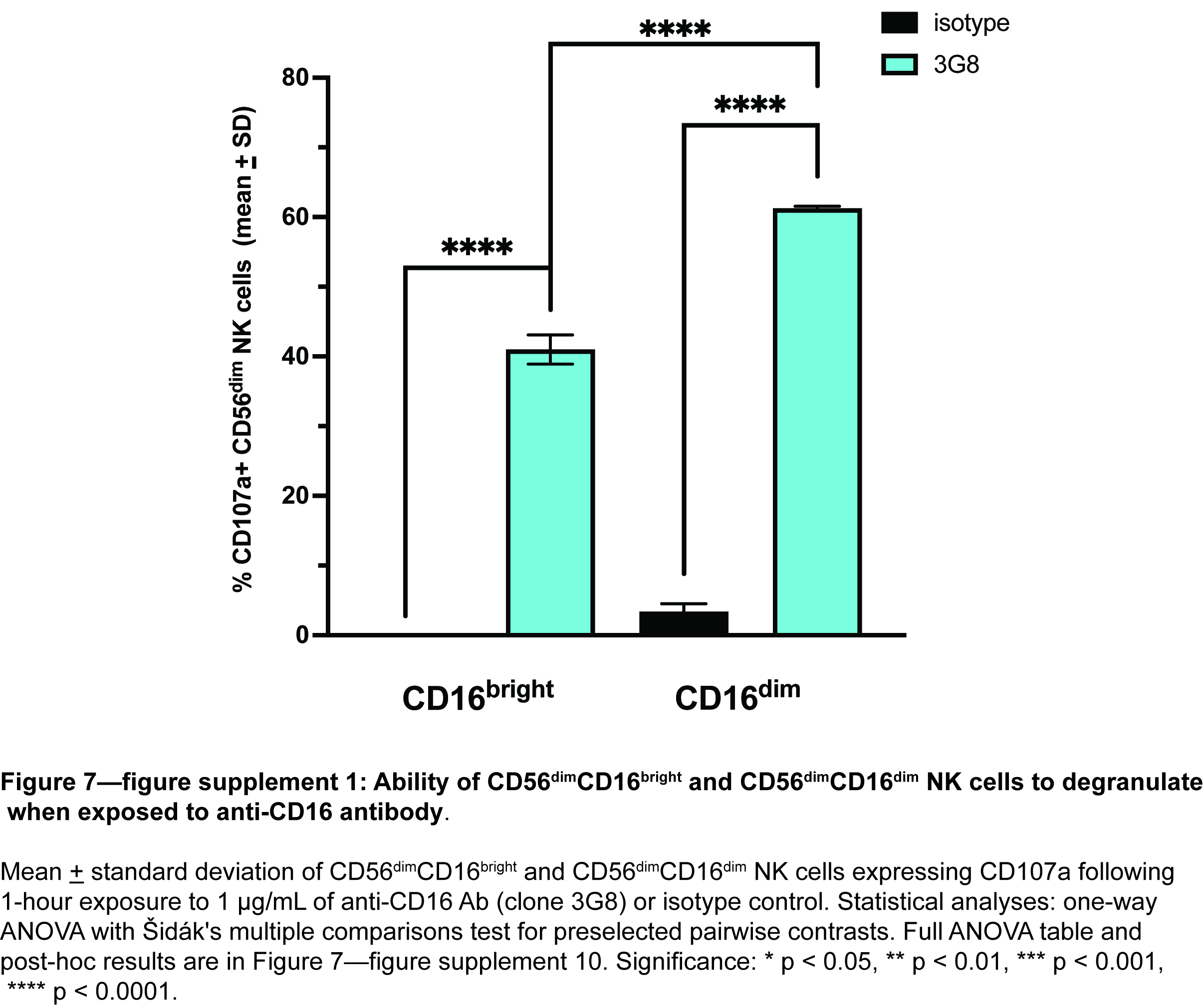
